# Multivariate optimization of k for k-nearest-neighbor feature selection with dichotomous outcomes: complex associations, class imbalance, and application to RNA-Seq in Major Depressive Disorder

**DOI:** 10.1101/2022.05.19.492724

**Authors:** Bryan A. Dawkins, Brett A. McKinney

## Abstract

Optimization of nearest-neighbor feature selection depends on the number of samples and features, the type of statistical effect, the feature scoring algorithm, and class imbalance. We recently reported a fixed-k for Nearest-neighbor Projected-Distance Regression (NPDR) that addresses each of these parameters, except for class imbalance. To remedy this, we parameterize our NPDR fixed-k by the minority class size (minority-class-k). We also introduce a class-adaptive fixed-k (hit-miss-k) to improve performance of Relief-based algorithms on imbalanced data. In addition, we present two optimization methods, including constrained variable-wise optimized k (VWOK) and a fixed-k derived with principal components analysis (kPCA), both of which are adaptive to class imbalance. Using simulated data, we show that our methods significantly improve feature detection across a variety of nearest-neighbor feature scoring metrics, and we demonstrate superior performance in comparison to random forest and ridge regression using consensus-nested cross-validation (cnCV) for feature selection. We applied cnCV to RNASeq expression data from a study of Major Depressive Disorder (MDD) using NPDR with minority-class-k, random forest, and cnCV-ridge regression for gene importance. Pathway analysis showed that NPDR with minority-class-k alone detected genes with clear relevance to MDD, suggesting that our new fixed-k formula is an effective rule-of-thumb.

## 1 INTRODUCTION

Nearest-neighbor feature selection is a filter-based class of methods for efficiently detecting main effects, interaction effects, and other complex variable-to-outcome associations in large datasets without the need for model specification [1]. However, optimization of their performance is a multifactorial problem, depending on factors like sample size, dimension of the space of predictors, the choice of feature scoring metric, and whether more weight should be given to interaction effects or main effects. Various approaches have been introduced to account for sample size and group imbalance [1], adjust for asymptotic moments of sample distances [2, 3], and tune neighbors for detecting interactions and main effects [4, 5]. In order to extend Relief-based feature selection to allow hypothesis testing for statistical significance of features, Statistical Inference Relief (STIR) reformulated the Relief scores into a pseudo t-test [4]. More recently, Nearest-neighbor Projected-Distance Regression (NPDR) was developed, in part, to extend the ability of STIR to do hypothesis testing for quantitative outcomes as well as case-control, while allowing for covariate adjustment (e.g., age, sex, and ethnicity) [3]. Despite the large variety of feature scoring methods, including many variations of Relief-based methods, STIR, and NPDR, the problem of generalized neighborhood optimization across feature scoring methods has not been addressed.

Previously, we introduced an analytical fixed-k [2, 3] that is a highly accurate asymptotic approximation to the average neighborhood size in Multiple instance Spatially Uniform ReliefF (MultiSURF) algorithm, which uses an adaptive-radius informed by moments of the target distance distribution [1]. Contrary to the adaptive-radius of MultiSURF, this fixed-k is based on a normal approximation to the target distance distribution, providing more efficient neighborhood calculations without sacrificing performance in feature detection. Using simulated data, our distance distributionally informed fixed-k shows considerable improvement over the naïve rule-of-thumb k=10 for detecting interaction effects and main effects within the context of ReliefF feature selection [2]; similarly, this fixed-k with NPDR yielded significantly better performance than both random forest and ReliefF [3]. However, we did not address the problem of class imbalance for classification in our previous work. In the present study, we extended our previous fixed-k for NPDR to account for group imbalance by parameterizing our formulation by the minority class size (minority-class-k), and thereby addressed an all too common issue in the classification setting. Applying NPDR to imbalanced data simulations, we showed significantly better performance for our minority-class-k than not only its predecessor [2, 3] but also the adaptive-radius neighborhoods used in MultiSURF [1]. We also introduced a class-adaptive fixed-k formulation (hit-miss-k) that allows different quantities of within-class neighbors (hits) and between-class neighbors (misses), dramatically improving performance over the single fixed-k ReliefF [1] and STIR [4] algorithms on imbalanced data.

Typically, one thinks of neighborhood adaptability at the level of samples or instances, as in the case of MultiSURF [1]. However, a variable-wise optimized k (VWOK) approach, developed for gene expression data [5], allows each independent variable/feature to have its own value of fixed-k that maximizes its importance score. This requires calculation of feature scores for all possible values of fixed-k, resulting in a feature-by-k score matrix from which maximal scores are derived. Although this type of adaptability leads to improved recall for main effects and interaction effects, it is computationally intensive and does not address the issue of group imbalance for classification. To improve efficiency and maintain variable-wise adaptability, while also providing adjustment for class imbalance, we used our minority-class-k to create a constrained grid of fixed-k values over which optimization is performed that shifts according to the degree of group imbalance. In this case, feature scores are maximized across a proper subset of all possible values of fixed-k. On the other hand, rather than choosing values of fixed-k that maximize feature scores, relying on the assumption that maximum scores are in fact optimal, one may instead desire to optimize fixed-k based on explained variance of feature scores in the feature-by-k score matrix. As an alternative to our constrained VWOK method, we introduced an approach that uses principal component analysis, choosing the value of fixed-k with the largest contribution to the first principal component (kPCA). This approach could be useful if the distribution of maximum feature scores deviates from the truly optimal feature ranking.

Finally, we applied NPDR with our new minority-class-k to real RNA sequencing data with class imbalance from a study of Major Depressive Disorder (MDD) [6]. The goal of feature selection is to find features that are relevant to the outcome of interest, however feature relevance is not necessarily reflected in classification accuracy [2]. Therefore, we relied on biological pathways from ConsensusPathDB [7] to make comparisons between gene sets from different feature selection methods, finding that NPDR with minority-class-k was the best method overall for detecting genes involved in central nervous system pathways. Thus, our novel approaches, now including adjustment for class imbalance, provide an ideal candidate for a rule-of-thumb neighborhood method for nearest-neighbor feature selection that can lead to discovery of biologically relevant predictors of complex diseases.

## 2 MATERIALS AND METHODS

Imbalanced data poses a challenge to feature selection algorithms in general. When data is balanced, nearest-neighbor feature importance scores have similar behavior for performance metrics like Area Under the Precision-Recall Curve (AUPRC) as neighborhood size (k) increases (Figs. 1 and 2 – top row). When data is imbalanced, nearest-neighbor feature importance scores remain largely unchanged, but the fixed-k that maximizes AUPRC shifts significantly to the left in favor of smaller neighborhood size (Figs. 1 and 2 – bottom row).

**Fig. 1.**
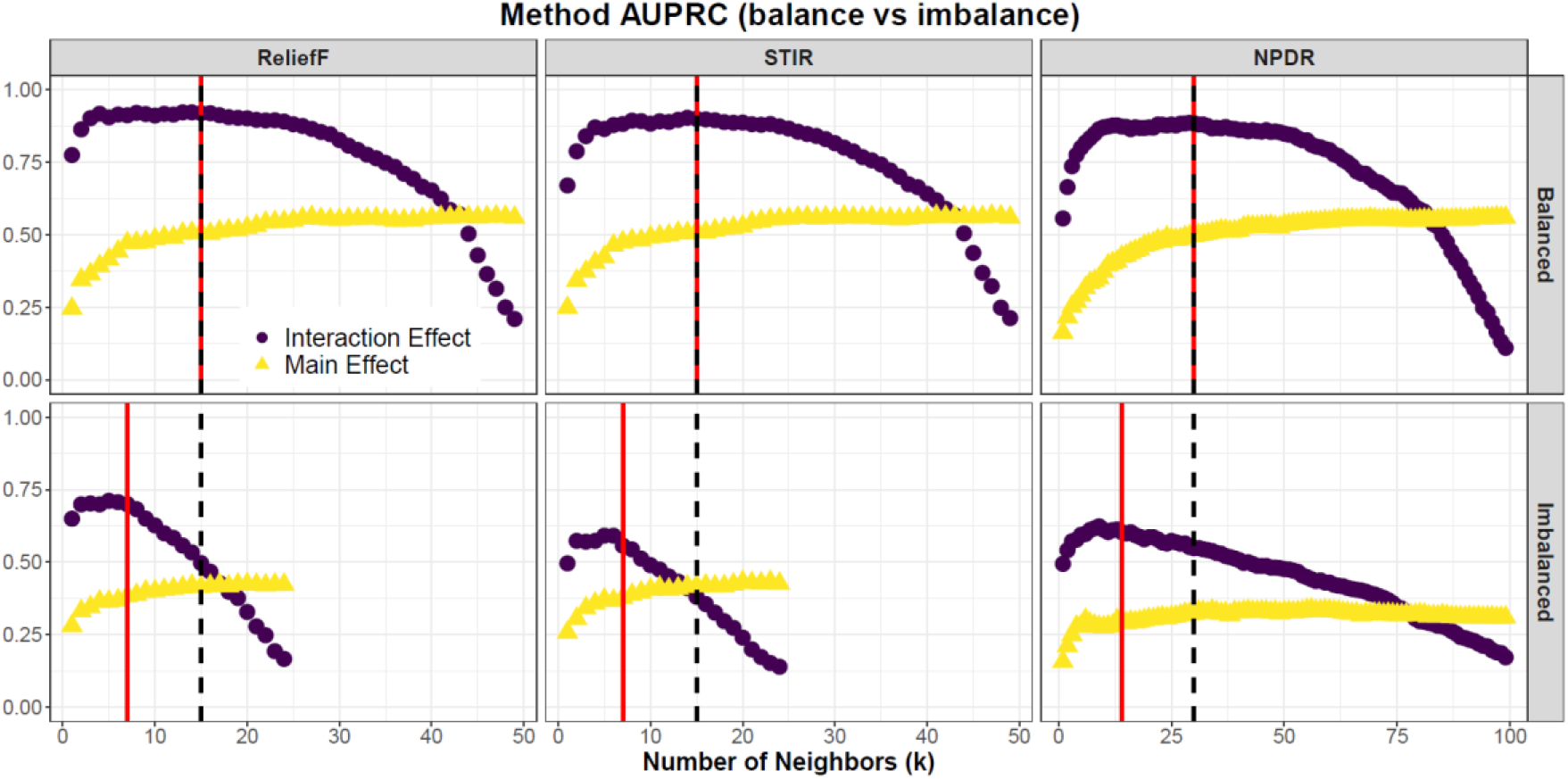
Comparison of method performance for balanced and imbalanced simulated data. Interaction effects and main effects in the presence of noise are simulated separately, where each simulated dataset has m=100 samples, p=100 features, with 10 functional variables and 90 irrelevant (noise) variables. Black-dashed and red-solid vertical lines are the non-adjusted fixed-k (Eq. 1) and imbalance-adjusted fixed-k (Eq. 2), respectively. Imbalance simulations have outcome with ratio of 25:75 (case:control). Each point is the average Area Under the Precision-Recall Curve (AUPRC) for ReliefF, STIR, and NPDR, respectively, for 20 simulation replicates. The optimal choice of k for detecting interaction effects shifts to the left in the presence of imbalance, whereas we see a flattening in the AUPRC curve for main effects. Our imbalance-adjusted fixed-k (vertical, red-solid line) shifts to the left, improving overall detection of functional features.

**Fig. 2.**
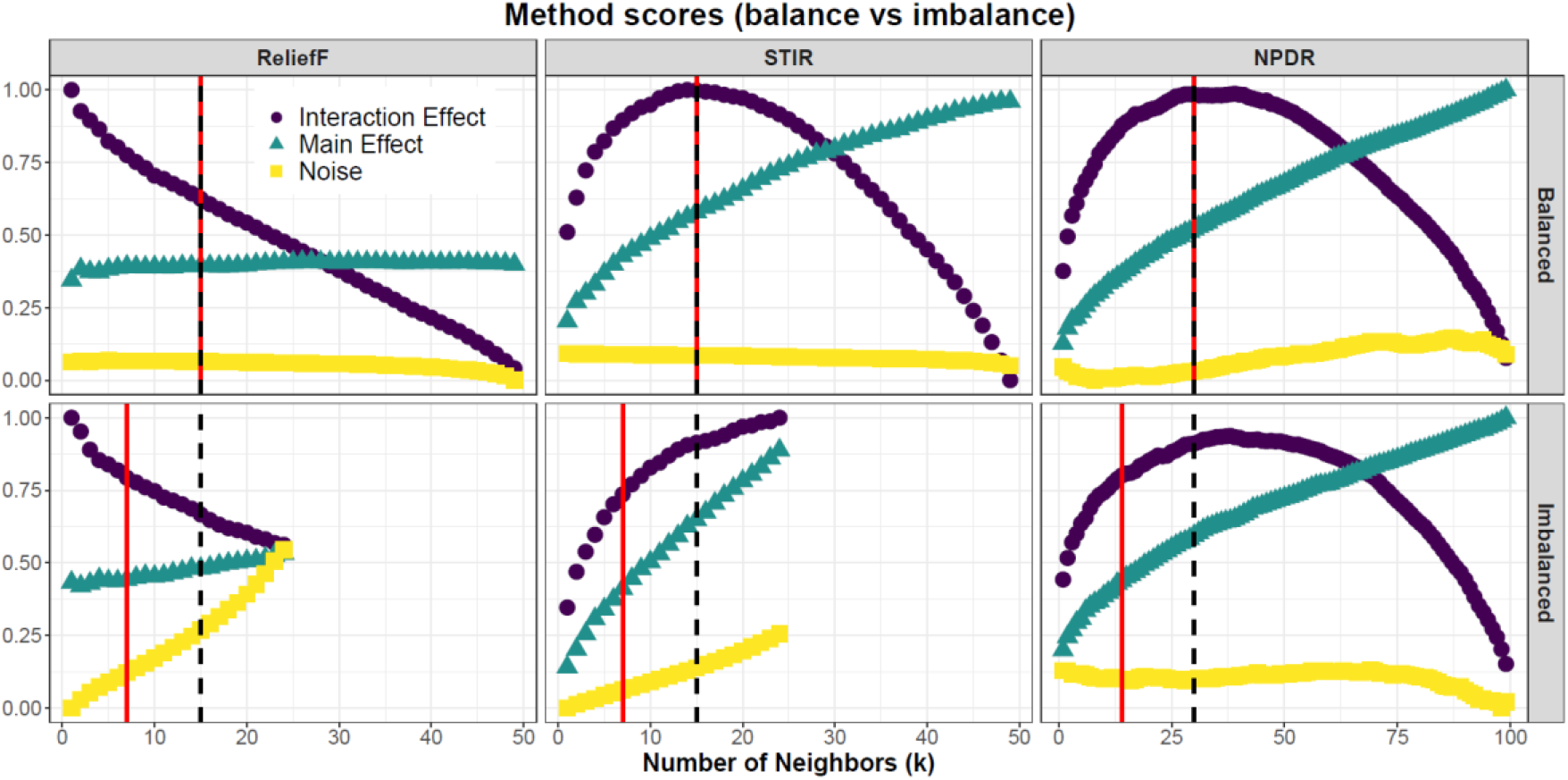
Comparison of method importance scores for balanced and imbalanced simulated data. Interaction effects and main effects in the presence of noise are simulated separately, where each simulated dataset has m=100 samples, p=100 features, with 10 functional variables and 90 irrelevant (noise) variables. Black-dashed and red-solid vertical lines are the non-adjusted fixed-k (Eq. 1) and imbalance-adjusted fixed-k (Eq. 2), respectively. Imbalance simulations have outcome with ratio of 25:75 (case:control). Each point is the average importance score for ReliefF, STIR, and NPDR, respectively, for 20 simulation replicates. With the exception of NPDR, importance scores are sensitive to imbalance. Our imbalance-adjusted fixed-k (vertical, red-solid line) shifts to the left, resulting in reduced variable scores relative to the non-adjusted fixed-k (vertical, black-dashed line) but also superior performance (Fig.1).

One adaptive neighborhood approach is to only allow neighbors of an instance that are within a certain radius. The adaptive *α*-radius for instance *i*. is given by

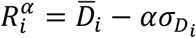

where *D* is the *m*×*m* sample distance matrix,

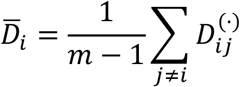

is the average distance for subject *i*. with respect to all other subjects,

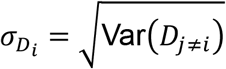

is the standard deviation, and *α* is the fraction or number of standard deviations away from the mean that the neighborhood radius can reach. Previously we derived the fixed-k that asymptotically approximates the *α*-radius [2, 3], given by the following function

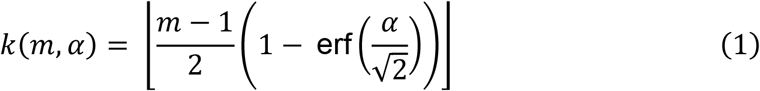

where *m* is the total number of training instances, erf is the Gaussian error function. For example, *α* = 0.5 would approximate the MultiSURF [1] neighborhood. This value of fixed-k leads to similar performance as MultiSURF radii with NPDR feature selection [3], where MultiSURF neighborhoods have been shown to successfully detect both interactions and main effects in simulated data [4].

### 2.1 IMBALANCE ADJUSTMENT FOR FIXED-K

However, *k*(*m,α*) (Eq.1) does not adjust for imbalance because it is parameterized by the total number of training instances *m*. In order to adjust fixed-k for imbalance in the case of NPDR, we let *k*(*m,α*) be a functional composition

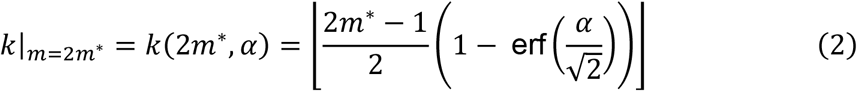

where *m*\* is the minority class size in a binary response (minority-class-k). That is, if *m*_1_ and *m*_2_ (*m* = *m*_1_ + *m*_2_) are the class sizes in a binary response, then *m*\* = min{*m*_1_, *m*_2_}. When data is balanced (i.e. *m*_1_ = *m*_2_), we have equivalence *k*(*m, α*) = *k*(2*m*, α*), however, imbalance causes *k*|_*m*=2*m*\*_ to shift leftward relative to our original *k*(*m, α*) formulation of fixed-k (Eq. 1). The motivation for using the minority class sample size for our new fixed-k (Eq. 2) in case-control data was derived from examination of AUPRC (Fig. 1) and importance scores (Fig. 2) as the level of imbalance in simulated data became more extreme. We found significant improvement in performance of ReliefF, STIR, and NPDR (Fig. 1 – bottom row) because the minorit-class-k (Eq. 2) shifted to be within a range of optimal AUPRC, whereas our original formulation (Eq. 1) fell outside the optimal AUPRC range. When the outcome was balanced, AUPRC and feature importance were positively correlated across all possible values of fixed-k (Fig. 1 and 2 – top row), indicating that maximizing feature scores for interaction or main effects leads to improved performance. When the outcome was no longer balanced, AUPRC and feature importance decorrelated (Fig. 1 and 2 – bottom row), implying that fixed-k that optimized AUPRC was no longer equivalent to that which maximized importance scores.

For each target instance in the training data, relief-based algorithms calculate the difference of the average miss and hit magnitude differences with respect to a given feature, with weighting of the magnitude differences (e.g., diff) of the target instance’s miss-neighbors to adjust scores for imbalance [1]. However, ReliefF uses the same value of fixed-k for hit and miss neighborhoods, which excludes much of the information contained in the full neighbor distribution. To allow class-adaptive fixed-k in ReliefF and STIR (hit-miss-k), we introduced the following equations:

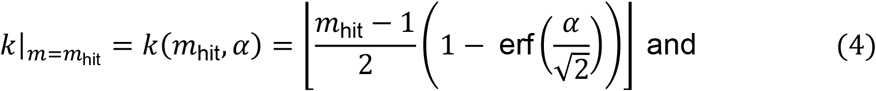

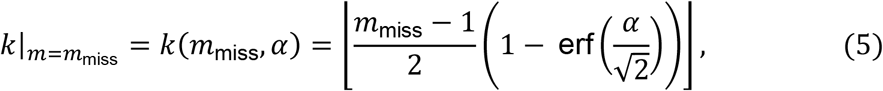

where *m*_hit_ and *m*_miss_ are the total number of possible hit and miss neighbors, respectively, for a given target instance.

### 2.2 CONSTRAINED VARIABLE-WISE OPTIMIZED K (VWOK)

The adaptive-radius neighborhood discussed previously is instance-wise adaptive [1], and we have shown that this can be approximated by a fixed-k (Eqs. 1 and 2) [2, 3]. Another important way to adapt the neighborhood is with respect to the features or variables (i.e., variable-wise). On average, it is possible to improve recall (or sensitivity) in detecting functional effects by applying the VWOK algorithm for maximizing feature scores as a function of k. However, using unconstrained k-grids is computationally intensive and leads to a loss in precision, allowing selection of irrelevant features that attain higher importance scores. Another limitation of VWOK is that it does not adapt to imbalance, resulting in diminished performance due to maximal feature scores decorrelating from maximal AUPRC (Figs. 1 & 2 – bottom row). To adjust VWOK for imbalance in the case of NPDR, we maximized standardized NPDR coefficients over the following grid of fixed-k values:

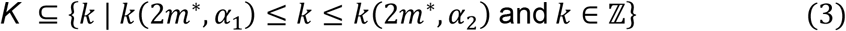

where *k*(*2m*, α_i_*) is the minority-class-k (Eq. 2) at *α*-radius (*i* ∈ {1,2}), 0.5 ≤ *α*_2_ < *α*_1_ ≤ 1, and |*K*| = *L*. We found empirically that setting *α*_1_ = 1 and *α*_2_ = 0.5 yielded optimal performance with extensive simulation analyses, experimenting with interaction and main effect sizes and various degrees of imbalance in dichotomous outcomes. The size of the k-grid (*L*) depends on the bounds *k*(2*m*, α*) (*i* ∈ {1,2}) and the chosen step size, however we used step size greater than 1 for efficiency. Calculation of maximal scores first requires generation of the full feature-by-*L* matrix of feature scores. Features are then ranked with respect to these maximal importance scores.

The standard VWOK neighborhood algorithm with ReliefF or STIR feature scoring uses a single grid of fixed-k values, where there is an equal number of hit and miss neighbors for each value of fixed-k in the grid [5]. To allow for class-adaptive VWOK in ReliefF and STIR, we introduced the following equations:

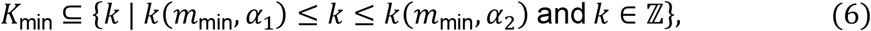

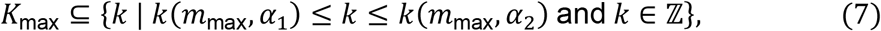

where *m*_min_ and *m*_max_ are the sizes of the smallest and largest class group, respectively, in the data, *k*(·, *α_i_*) is our analytical fixed-k (Eq. 1) evaluated at *α*-radius *α_i_* (*i* ∈ {1,2}), and 0.5 ≤ *α*_2_ < *α*_1_ ≤ 1. There is a one-to-one correspondence between grid values in *K*_min_ and *K*_max_ (i.e. |*k*_min_| = |*K*_max_ | = *L*), implying that the maximum score for each feature will occur at one of the ordered pairs 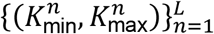. As in the case of the VWOK k-grid for NPDR, the size (*L*) of the class-adaptive k-grid depends on the bounds *k*(·, *α_i_*) (*i* ∈ {1,2}) and the chosen step size.

### 2.3 FIXED-K DERIVED WITH PCA (KPCA)

Ideal choices of fixed-k allow for partitioning relevant feature scores and irrelevant feature scores into distinct clusters. That is, the average relevant feature score is significantly larger than the average irrelevant feature score for the ideal value of fixed-k. In the case of NPDR, we denote the matrix of standardized feature importance scores from our constrained k-grid (Eq. 3) by *Ŝ*^*p*×*L*^, where *p* is the total number of features and *L* is the size of the k-grid (Fig. 3A). The first principal component of Ŝ^*p*×*L*^ (Fig. 3B) explains the largest proportion of the variance in feature importance scores across all values of fixed-k. The percent contribution of each value of fixed-k is calculated as the maximum squared variable loading on the first principal component of Ŝ^*p*×*L*^ (Fig. 3C). The value of fixed-k with the largest percent contribution (Fig. 3D) takes into account not only outcome imbalance but also explained variance in importance scores across k-grid values.

**Fig. 3.**
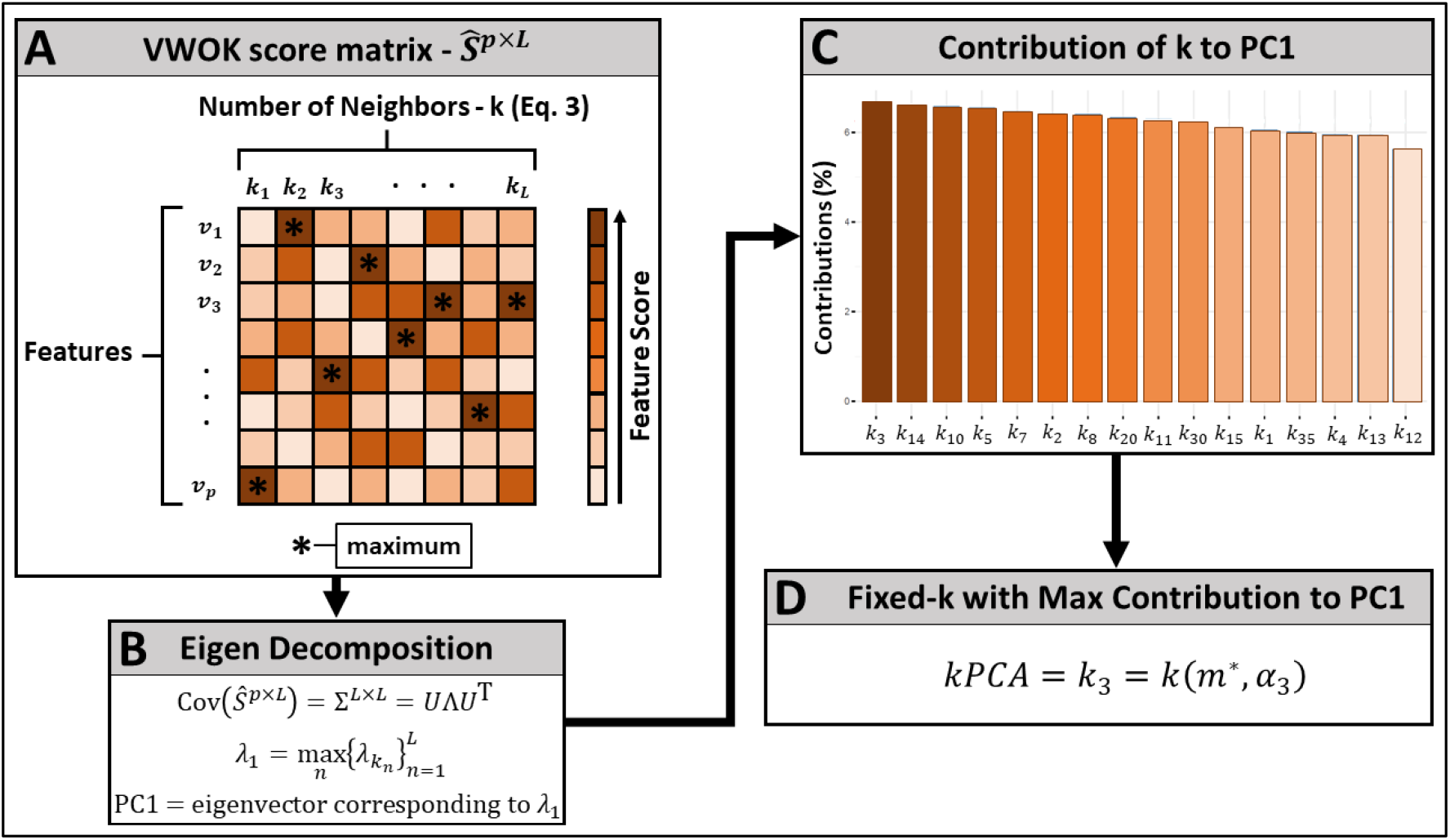
Choosing k by maximum feature score and by contribution to first principal component of feature-by-k score matrix. (**A**) Feature scores are calculated for each value of fixed-k in our constrained, imbalanced-adaptive k-grid (Eq. 3). VWOK scores are calculated from the unstandardized score matrix *Ŝ*^*p*×*L*^, which is the maximum score as a function of k for each feature (*v_a_, a* = 1,…,*p*). (**B**) The first principal component is determined for the standardized score matrix *Ŝ*^*P*×*L*^. (**C**) For each k in the constrained k-grid (Eq. 3), percent contribution to the first principal component is calculated as the squared loading on the first principal component. (**D**) The value of fixed-k with the largest percent contribution to the first principal component of the standardized score matrix *(kPCA),* is used to extract the corresponding feature scores from the unstandardized score matrix *S*^*p*×*L*^, which is *k*_3_, in this toy example.

Due to the one-to-one correspondence between the hit-miss-k grids for ReliefF and STIR (Eqs. 6 and 7), the standardized matrix of importance scores is denoted by *Ŝ*^*p*×*L*^ as in the case of NPDR described previously, where *p* is the total number of features and *L* is the size of each k-grid (Fig. 3A). The ordered pair of class-adaptive grid values 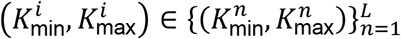 that had the largest percentage contribution to the first principal component of the standardized score matrix (Fig. 3B-C), along with the associated ReliefF or STIR importance scores, was chosen for further analysis (Fig. 3D).

### 2.4 RNA-SEQ DATA FOR ANALYSIS OF MDD

We used publicly available RNA-Seq data that was described previously [6]. This dataset consists of 15,231 total genes that were identified in individuals with Major Depressive Disorder (MDD, n = 463) and healthy controls (HC, n = 452). We used this dataset previously to identify genes associated with MDD, with adjustment for biological sex [3]. Because the data were approximately balanced, having about the same number of MDD and HC, we randomly sampled a subset of HC (n=115) to artificially create an approximately 4:1 ratio of MDD-to-HC. Genes that were detected for relevance to MDD were submitted to ConsensusPathDB [7] (version 35 – released June, 19 2021) for biological pathways, which is an integrated collection of molecular interaction data from 32 different public repositories.

### 2.5 SIMULATED DATA

Simulated case-control data contained a combination of functional (i.e. relevant to the outcome) and irrelevant features. We used previously published methodologies for simulating main effects [8] and network-based interactions [9] which have been employed for assessing feature selection and model classification performance in a settings common to high-dimensional bioinformatic data [2–4, 10], where the signal-to-noise ratio is often quite low. We used the createSimulation() function from an R package called ‘privateEC’ (https://github.com/insilico/privateEC) nfor generating all main effect and interaction effect simulations.

#### 2.5.1 FEATURE SELECTION PERFORMANCE METRICS

The performance of each method was determined by its ability to detect functional features (True Positives, TP) in simulated data. To this end we used precision, recall, Area Under the Precision-Recall Curve (AUPRC), and Matthew’s Correlation Coefficient (MCC) [11] as performance measures. As described elsewhere[3], AUPRC measures how well a given feature selection algorithm delineates between true and false positives simultaneously across a grid of call thresholds. Thus, AUPRC can be used as a global performance measure for a given set of ranked feature scores, allowing for comparisons between different scoring methods with respect to the entire feature ranking. When a fixed call threshold was used for selecting features, such as adjusted p-values from NPDR [3], we used precision, recall, and MCC to measure feature selection performance. Like AUPRC, MCC can be used to measure a feature selection method’s performance in detecting true positives while avoiding detection of false positives. Unlike AUPRC, MCC can be used for method comparison with respect to a single, fixed call threshold. For methods lacking a measure of statistical significance (e.g., ReliefF or random forest), we calculated the maximum value of MCC across a grid of call thresholds, selecting up to 20% of ranked features. For NPDR, we used Bonferroni corrected p-values as the call threshold for calculating MCC. We did this to compare the MCC of NPDR-selected features, with its corresponding p-value threshold, to the optimized MCC of other feature scoring methods.

#### 2.5.2 FEATURE SCORES AND AUPRC VERSUS FIXED-K

For assessing feature importance scores and the performance of nearest-neighbor feature selection algorithms as a function of fixed-k, we generated 20 simulation replicates with *m* = 100 samples and *p* = 100 features with 10% functional. In these comparisons, we analyzed detection of main effects and interaction effects separately. That is, one simulation scenario (20 replicates) had 10 features with main effect and 90 remaining features that were irrelevant to the outcome. The next scenario (20 replicates) had 10 features involved in feature-feature interactions and 90 remaining features that were irrelevant to the outcome. The effect size parameter *(bias)* for main effects was set to 0.8, which has been shown elsewhere to correspond to approximately 40% power [10]. The effect size parameter *(bias)* for interaction effects was set to 0.4, which is inversely proportional to the average functional feature-feature correlation in ‘controls’. These same correlations are destroyed in ‘cases’ through a process of random permutation, thus creating differential correlation between ‘cases’ and ‘controls’ [9].

#### 2.5.3 ALGORITHM PERFORMANCE COMPARISONS

Each simulated dataset used for assessing feature selection performance had *m* = 100 samples and *p* = 1000 features with 10% functional. The effect size parameter (bias) was set to 0.8 and 0.4 for main effects and interaction effects, respectively. Functional features were mixed, where a fixed proportion (*ρ*) were involved in only interaction effects with no main effect while the remaining proportion (1 – *ρ*) had only main effect. Three scenarios were tested: (i) *ρ* = 0.5 (50% interaction/50% main effect), (ii) *ρ* = 0.75 (75% interaction/25% main effect), and (iii) *ρ* = 0.25 (25% interaction/75% main effect). For brevity, we include here illustrations for only the first scenario (*ρ* = 0.5), leaving those remaining (*ρ* = 0.75 and *ρ* = 0.25) to supplemental materials. Comparisons between different feature selection methodologies were made with respect to raw feature scores and within consensus-features nested cross-validation (cnCV) [12]. We used 100 simulation replicates for raw method comparisons outside cnCV, while 50 replicates were used within cnCV. The cnCV method was implemented using the function consensus_nestedCV() from an R package called ‘cncv’ (https://github.com/insilico/cncv). For comparisons within cnCV, we first split each dataset into training and test sets (70% training/30% test). We used 5 folds (ncv_folds = c(5, 5)) for both inner and outer loops of the nested procedure, selecting the top 30% (inner_selection_percent = 0.3) of ranked features within inner training folds. Random forest, as implemented in the R package ‘randomForest’, was the learning method (learning_method = “rf”) used in combination with each feature importance algorithm.

## 3 RESULTS

### 3.1 VWOK FEATURE SCORES OPTIMIZE ALL METHODS

We investigated the quality of raw feature scores for NPDR, ReliefF, and STIR for each of our imbalance-adjusted approaches using simulated data; simulation scenarios below are partitioned into two types: equal (*ρ* = 0.5) and unequal (*ρ* = 0.75,0.25) proportion, respectively, of functional main effects and interaction effects.

#### 3.1.1 EQUAL PROPORTION MAIN AND INTERACTION

Using simulated case-control data of equal proportions main effect and interaction effect (*ρ* = 0.5: 50% interaction/50% main), we compared the feature-based AUPRC (Fig. 4A) and MCC (Fig. 4B) between minority-class-k (Eq. 2), constrained VWOK, and kPCA with NPDR feature scoring. The AUPRC (Fig. 4A) is approximately the same between all three methods for both balanced and imbalanced dichotomous outcome, suggesting that the full list of ranked features has about the same quality for each neighborhood-NPDR pairing. On balanced data, the MCC for constrained VWOK is significantly higher than minority-class-k (Eq. 2) and kPCA (Fig. 4B – left panel), however all methods have about the same MCC for imbalanced data (Fig. 4B – right panel). This increase in MCC for VWOK suggests its modest superiority over the two other novel approaches when using adjusted NPDR p-values. Using our hit-miss-k (Eqs. 4 – 5), constrained VWOK (Eqs. 6 – 7), and kPCA, we found similar results for ReliefF (Supplementary Fig. 1) and STIR (Supplementary Fig. 2).

**Fig. 4.**
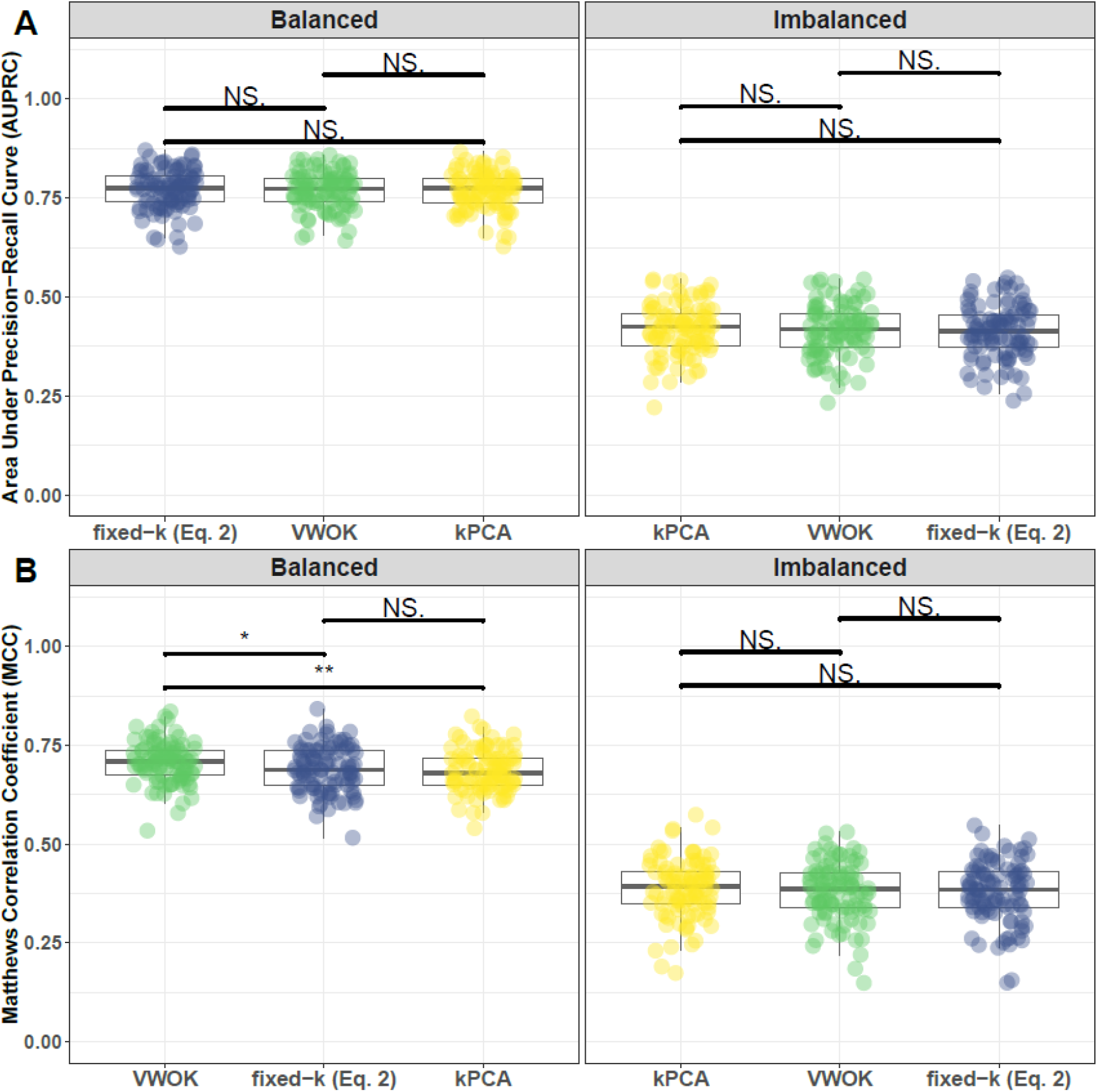
Performance comparison for fixed-k (Eq. 2), VWOK, and kPCA with NPDR feature scoring. Performance of feature selection was measured for 100 simulation replicates. Each simulated data set had *m* = 100 instances and *p* = 1000 features with 100 functional. Functional features included 50 that with main effect (bias_main_ = 0.8) only and the remaining 50 were involved in network interactions (bias_int_ = 0.4) and had no main effect. Imbalanced simulations had class ratio of 25:75 (cases:controls). (**A**) Area Under Precision-Recall Curve (AUPRC) for each method, sorted by decreasing mean AUPRC. (**B**) Matthews Correlation Coefficient (MCC) for each method, sorted by decreasing mean MCC. Comparisons were made with Mann-Whitney U test (* *P* < 0.05 and ** *P* < 0.01).

#### 3.1.2 UNEQUAL PROPORTION MAIN AND INTERACTION

We analyzed performance of each pairing of neighborhood algorithm and feature scoring method for two additional scenarios. These were (i) *ρ* = 0.75: 75% interaction/25% main and (ii) *ρ* = 0.25: 25% interaction/75% main. Overall, we found that VWOK was modestly superior to fixed-k (Eq. 2) and kPCA in the case of 75% interaction effect (Supplementary Figs. 8 – 10) and 25% interaction effect (Supplementary Figs. 17 – 19) simulations; this was true for NPDR (Supplementary Figs. 8 & 17), ReliefF (Supplementary Figs. 9 & 18), and STIR (Supplementary Figs. 10 & 19).

### 3.2 FIXED-K OUTPERFORMS ADAPTIVE RADIUS

We compared performance of minority-class-k (Eq. 2), non-adjusted fixed-k (Eq. 1), and the MultiSURF adaptive-radius with respect to raw NPDR feature scores. In addition, we compared hit-miss-k (Eqs. 4 – 5), non-adjusted fixed-k (Eq. 1), and MultiSURF adaptive-radius with respect to ReliefF and STIR feature scores. As in previous sections (3.1 – 3.2), results are partitioned according to the proportions of main effects and interaction effects in simulations.

#### 3.2.1 EQUAL PROPORTION MAIN AND INTERACTION

Using the same case-control simulation replicates mentioned previously (3.1), we compared the feature-based AUPRC (Fig. 5A) and MCC (Fig. 5B) between minority-class-k (Eq. 2), non-adjusted fixed-k (Eq. 1), and MultiSURF adaptive-radius with NPDR feature scoring. With balanced classes, performance was approximately the same from each pairing of neighborhood algorithm and NPDR feature scoring (Fig. 5 – left panels). On the contrary, both AUPRC and MCC were significantly higher for minority-class-k (Eq. 2) when classes were imbalanced (Fig. 5 – right panels). Comparison of neighborhood methods within the context of ReliefF and STIR feature scores differed from those previously mentioned for NPDR, particularly with respect to the MultiSURF adaptive-radius. Our hit-miss-k (Eqs. 4 – 5) resulted in significantly higher AUPRC and MCC on imbalanced data compared to non-adjusted fixed-k (Eq. 1) with ReliefF (Supplementary Fig. 4) and STIR (Supplementary Fig. 5). However, hit-miss-k (Eqs. 4 – 5) and MultiSURF adaptive-radius were roughly equivalent for both balanced and imbalanced designs Supplementary Figs. 4 – 5).

**Fig. 5.**
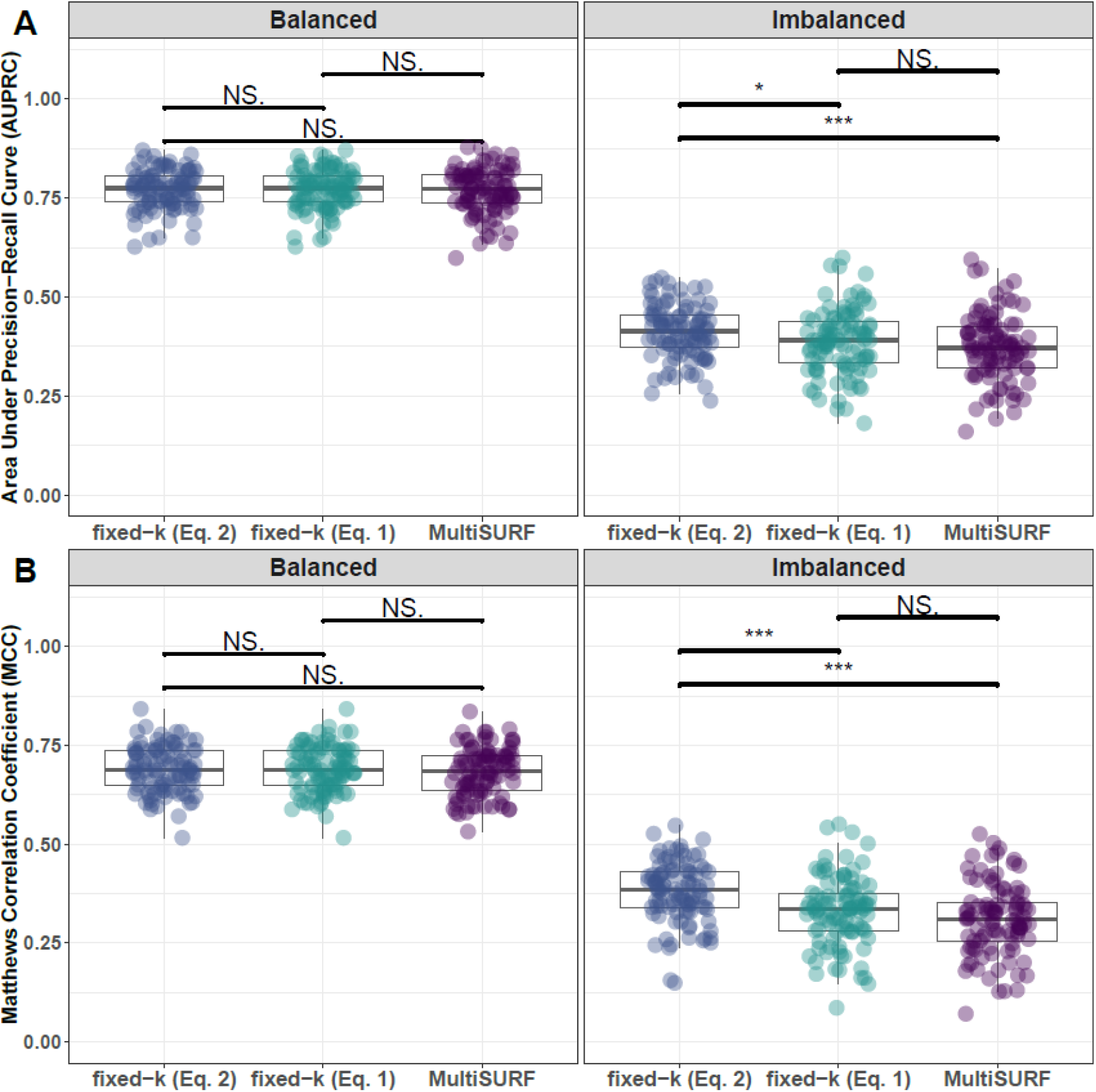
Performance comparison of imbalance-adjusted fixed-k (Eq. 2), non-adjusted fixed-k (Eq. 1), and MultiSURF with NPDR feature scoring. Performance of feature selection was measured for 100 simulation replicates. Each simulated data set had *m* = 100 instances and *p* = 1000 features with 100 functional. Functional features included 50 that with main effect (bias_main_ = 0.8) only and the remaining 50 were involved in network interactions (bias_int_ = 0.4) and had no main effect. Imbalanced simulations had class ratio of 25:75 (cases:controls). (**A**) Area Under Precision-Recall Curve (AUPRC) for each method, sorted by decreasing mean AUPRC. (**B**) Matthews Correlation Coefficient (MCC) for each method, sorted by decreasing mean MCC. Comparisons were made with Mann-Whitney U test (* *P* < 0.05 and *** *P* < 0.001).

#### 3.2.2 UNEQUAL PROPORTION MAIN AND INTERACTION

Varying the ratio of interaction effects to main effects as mentioned previously (i.e., *ρ* = 0.75: 75% interaction/25% main to *ρ* = 0.25: 25% interaction/75% main), we found similar results. That is, there was significantly higher AUPRC and MCC for minority-class-k (Eq. 2) compared to non-adjusted fixed-k (Eq. 1) and MultiSURF adaptive-radius for NPDR feature scores on imbalanced data (Supplementary Figs. 11 & 20). Similarly, hit-miss-k (Eqs. 4 – 5) and MultiSURF adaptive-radius had approximately the same performance, both of which had significantly higher AUPRC and MCC for ReliefF (Supplementary Fig. 12 & 21) and STIR (Supplementary Figs. 13 & 22) on imbalanced data.

### 3.3 NPDR, RELIEFF, AND STIR OUTPERFORM RANDOM FOREST AND Ridge REGRESSION ON SIMULATED DATA

Using the same case-control simulation replicates mentioned in preceding subsections (3.1 – 3.2), we compared the performance of minority-class-k (Eq. 2) for NPDR and hit-miss-k (Eqs. 4 – 5) for ReliefF and STIR to random forest and ridge regression. When there were at least as many interacting features as those with main effect only (*ρ* = 0.5,0.75), the feature-based AUPRC and MCC for NPDR with minority-class-k (Eq. 2) was significantly higher than for random forest and ridge regression for balanced and imbalanced data (Fig. 6 and Supplementary Figs. 14 & 17), indicating a general superiority of NPDR over these two commonly used methods. The same was true for hit-miss-k (Eqs. 4 – 5) for ReliefF (Supplementary Fig. 15) and STIR (Supplementary Fig. 16) compared to random forest and ridge regression.

**Fig. 6.**
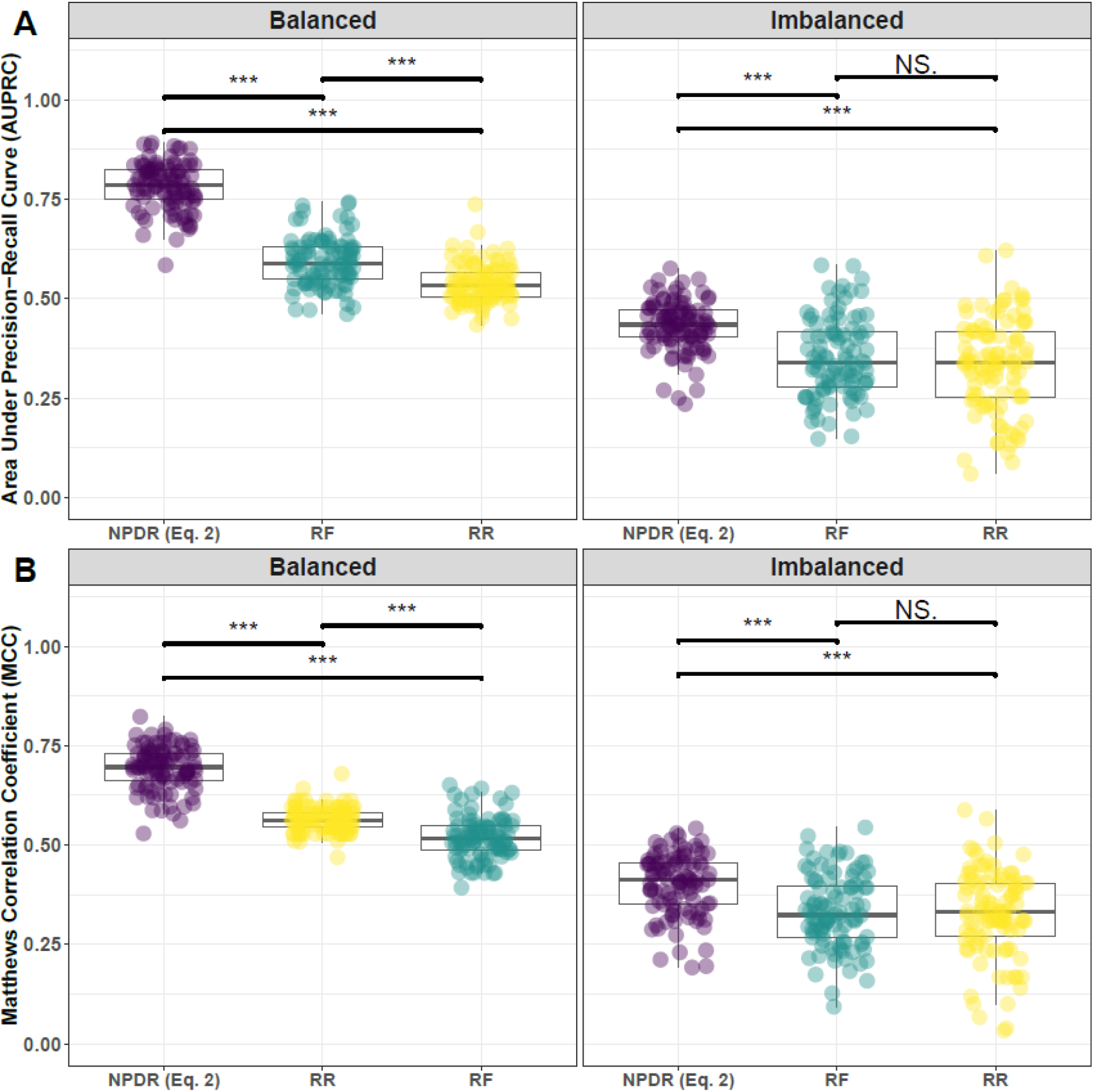
Performance comparison of NPDR with imbalance-adjusted fixed-k (Eq. 2), Random Forest (RF), and Ridge Regression (RR). Performance of feature selection was measured for 100 simulation replicates. Each simulated data set had *m* = 100 instances and *p* = 1000 features with 100 functional. Functional features included 50 that with main effect (bias_main_ = 0.8) only and the remaining 50 were involved in network interactions (bias_int_ = 0.4) and had no main effect. Imbalanced simulations had class ratio of 25:75 (cases:controls). (**A**) Area Under Precision-Recall Curve (AUPRC) for each method, sorted by decreasing mean AUPRC. (**B**) Matthews Correlation Coefficient (MCC) for each method, sorted by decreasing mean MCC. Comparisons were made with Mann-Whitney U test (* *P* < 0.05 and *** *P* < 0.001).

These same comparisons showed completely different results when there were fewer interacting features than main effects (*ρ* = 0.25). In the case of balanced data, NPDR with minority-class-k (Eq. 2) and ridge regression had approximately equal AUPRC, however ridge regression had significantly higher MCC; both minority-class-k (Eq. 2) NPDR and ridge regression had significantly higher AUPRC and MCC than random forest (Supplementary Fig. 23 – left panel). For imbalanced data, ridge regression had significantly higher AUPRC and MCC than both minority-class-k (Eq. 2) NPDR and random forest (Supplementary Fig. 23 – right panel). ReliefF with hit-miss-k (Eqs. 4 – 5) had significantly higher AUPRC and MCC than both ridge regression and random forest for balanced data (Supplementary Fig. 24 – left panel). On the contrary, ridge regression performed best on imbalanced data, attaining significantly higher AUPRC and slightly higher MCC than ReliefF with hit-miss-k (Eqs. 4 – 5) (Supplementary Fig. 24 – right panel). STIR with hit-miss-k (Eqs. 4 – 5) performed about the same as NPDR with minority-class-k (Eq. 2) (Supplementary Fig. 25).

### 3.4 NEIGHBORHOOD PERFORMANCE WITHIN CNCV

In addition to investigating the quality of raw feature scores for NPDR, ReliefF, and STIR for each of our imbalance-adjusted approaches, we used the same simulation scenarios mentioned in the preceding subsection (3.1) for comparison within consensus-features nested cross-validation (cnCV). This was done because nearest-neighbor feature selection is typically wrapped within cross-validation to avoid over-fitting.

#### 3.4.1 EQUAL PROPORTION MAIN AND INTERACTION

For simulations with equal parts interaction effect and main effect, minority-class-k (Eq. 2), constrained VWOK (Eq. 3), and kPCA had approximately the same feature-based MCC, precision, and recall for balanced (Fig. 7) and imbalanced (Fig. 8) data. In addition to assessing the quality of detected features from cnCV for each neighborhood-NPDR pairing, consensus features were used to train a random forest classifier for each simulation replicate, resulting in approximately equal balanced classification accuracy on training and validation sets (Figs. 7 – 8). Using our hit-miss-k (Eqs. 4 – 5), constrained VWOK (Eqs. 6 – 7), and kPCA, we found similar results for ReliefF (Supplementary Figs. 26 – 27) and STIR (Supplementary Figs. 29 – 30).

**Fig. 7.**
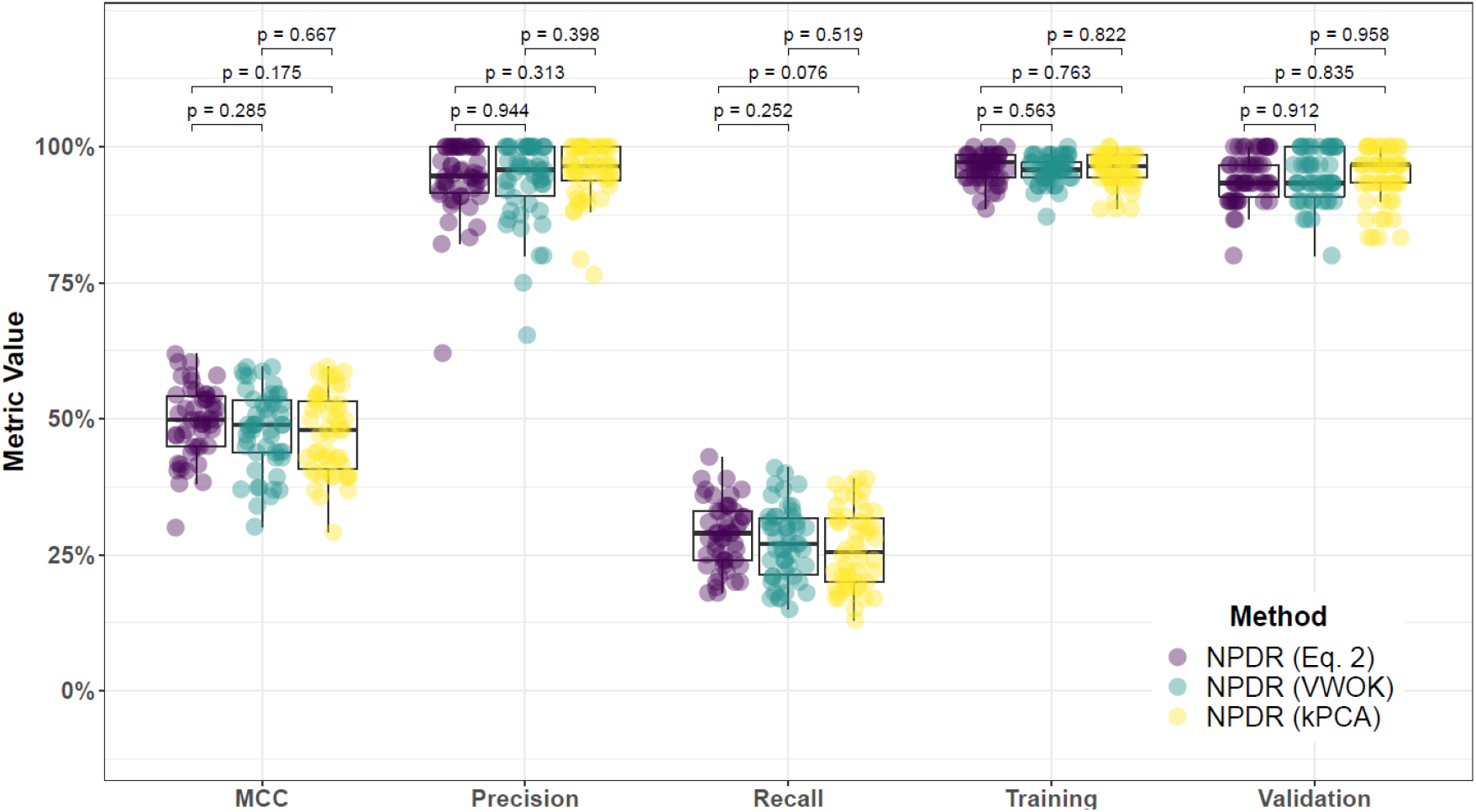
Performance comparison for fixed-k (Eq. 2), VWOK, and kPCA with NPDR feature scoring and consensus-features nested Cross-Validation (cnCV) on balanced data. Performance of feature selection was measured for 50 simulation replicates. Each simulated data set had *m* = 100 instances and *p* = 1000 features with 100 functional. Each simulated data set had balanced class groups with 50 ‘case’ and 50 ‘control’. Functional features included 50 with main effect (bias_main_ = 0.8) only and the remaining 50 were involved in network interactions (bias_int_ = 0.4) and had no main effect. Matthew’s Correlation Coefficient (MCC), precision, and recall were calculated based on detection of functional features (i.e. true positives). Training and validation represent the balanced classification accuracy of the random forest model that was fit using consensus features from the cnCV {Parvandeh, 2020 #11} on full training data (70 samples) and independent test data (30 samples), respectively.

**Fig. 8.**
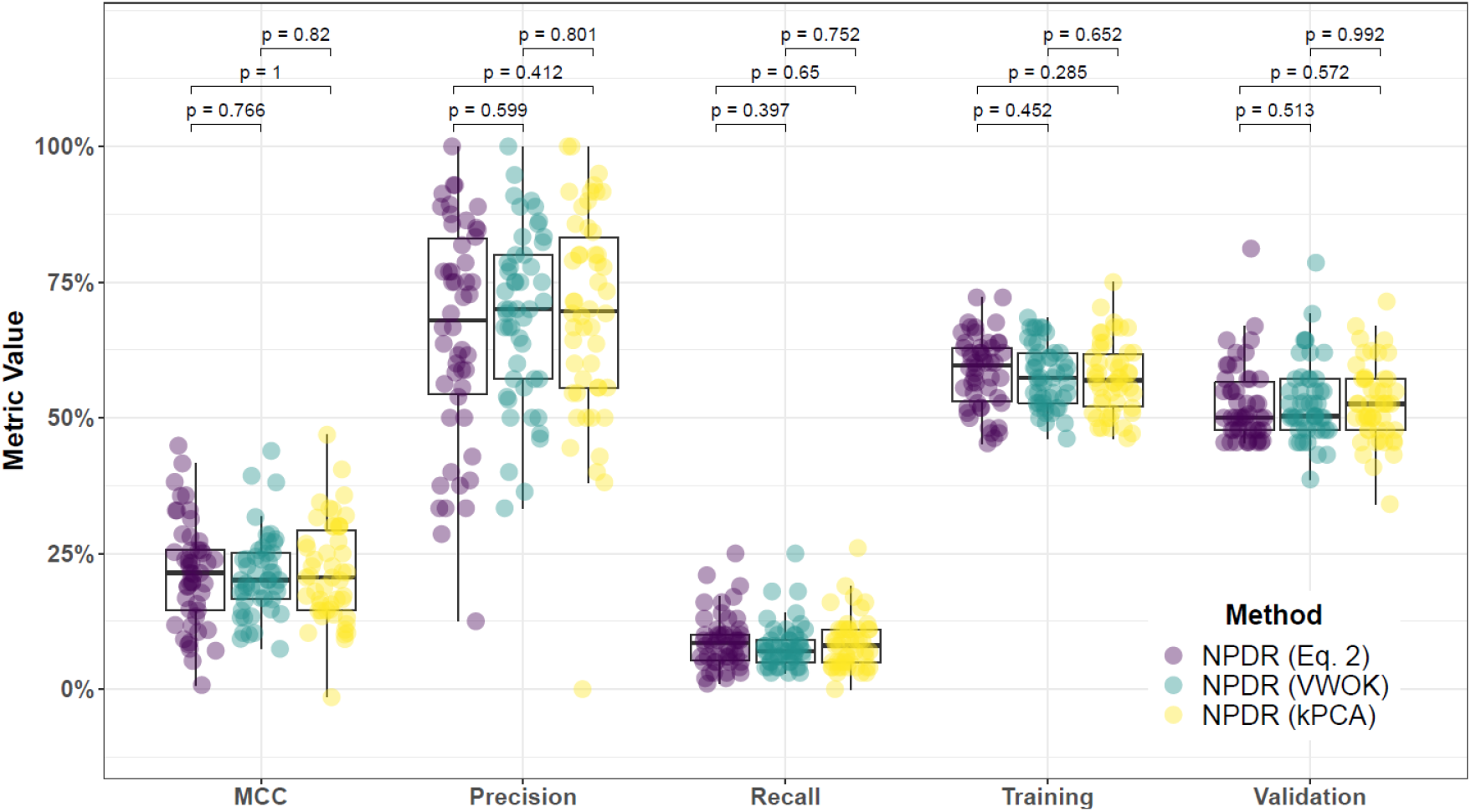
Performance comparison for fixed-k (Eq. 2), VWOK, and kPCA with NPDR feature scoring and consensus-features nested Cross-Validation (cnCV) on imbalanced data. Performance of feature selection was measured for 50 simulation replicates. Each simulated data set had *m* = 100 instances and *p* = 1000 features with 100 functional. Each simulated data set had imbalanced class groups with 25 ‘case’ and 75 ‘control’. Functional features included 50 with main effect (bias_main_ = 0.8) only and the remaining 50 were involved in network interactions (bias_int_ = 0.4) and had no main effect. Matthew’s Correlation Coefficient (MCC), precision, and recall were calculated based on detection of functional features (i.e. true positives). Training and validation represent the balanced classification accuracy of the random forest model that was fit using consensus features from the cnCV {Parvandeh, 2020 #11} on full training data (70 samples) and independent test data (30 samples), respectively.

#### 3.4.2 UNEQUAL PROPORTION MAIN AND INTERACTION

Varying the ratio of interaction effect to main effect (i.e., *ρ* = 0.75: 75% interaction/25% main to *ρ* = 0.25: 25% interaction/75% main) resulted in the same overall conclusions. Our minority-class-k (Eq. 2), constrained VWOK (Eq. 3), and kPCA methods had approximately equal performance within cnCV for NPDR (Supplementary Figs. 38 – 39 & 56 – 57). Hit-miss-k (Eqs. 4 – 5), constrained VWOK (Eqs. 6 – 7), and kPCA yielded approximately equal performance measures for ReliefF (Supplementary Figs. 40 – 41 & 58 – 59) and STIR (Supplementary Figs. 42 – 43 & 60 – 61). These results suggest that minority-class-k (Eq. 2) for NPDR and hit-miss-k fixed-k (Eqs. 4 – 5) for ReliefF or STIR should be used for the sake of efficiency within cross-validation.

### 3.5 CNCV-NPDR WITH MINORITY-CLASS-K DETECTS MDD ASSOCIATED GENES IN NERVOUS SYSTEM PATHWAYS

We applied cnCV to real RNA-Seq data from a study of MDD [6], calculating gene importance within inner training folds using NPDR with minority-class-k (Eq. 2), NPDR with non-adjusted fixed-k (Eq. 1), random forest, and ridge regression, respectively. Because the relationship between genes and outcome is not known a priori in real data, our goal was to compare functional relevance of gene sets (Table 1, Supplementary Table 2) between the different feature selection approaches. Specifically, we compared GO terms associated with genes detected by each combination of cnCV and feature scoring method (Fig. 9 & Table 2). We did not derive GO terms for cnCV-random forest because only a single gene (RAB26) was detected using this approach (Table 1). Even though total number of genes detected with each approach was comparable, with the exception of cnCV-random forest, genes detected using cnCV-NPDR with minority-class-k (Eq. 2) were associated with far more GO terms than those detected by cnCV-NPDR with non-adjusted fixed-k (Eq. 1) and cnCV-ridge regression (Fig. 9); this was true for biological pathways from other databases as well (Supplementary Fig. 74). We also found more GO terms corresponding directly to the nervous system for cnCV-NPDR with minority-class-k (Eq. 2) when compared to the cnCV-NPDR with non-adjusted fixed-k (Eq. 1) and -ridge regression (Fig. 9 – highlighted terms). However, the strongest GO term associations were found for genes detected by cnCV-ridge regression, corresponding to insulin-like growth factor binding. In addition, cnCV-ridge regression had the highest concordance between gene sets detected on the full (balanced) and sub-sampled (imbalanced) RNA-Seq data (Supplementary Table 2), suggesting less sensitivity to variation in training samples; there was very little concordance between gene sets detected by cnCV-NPDR (Eqs. 1 & 2) (Supplmentary Table 2).

**Figure 9.**
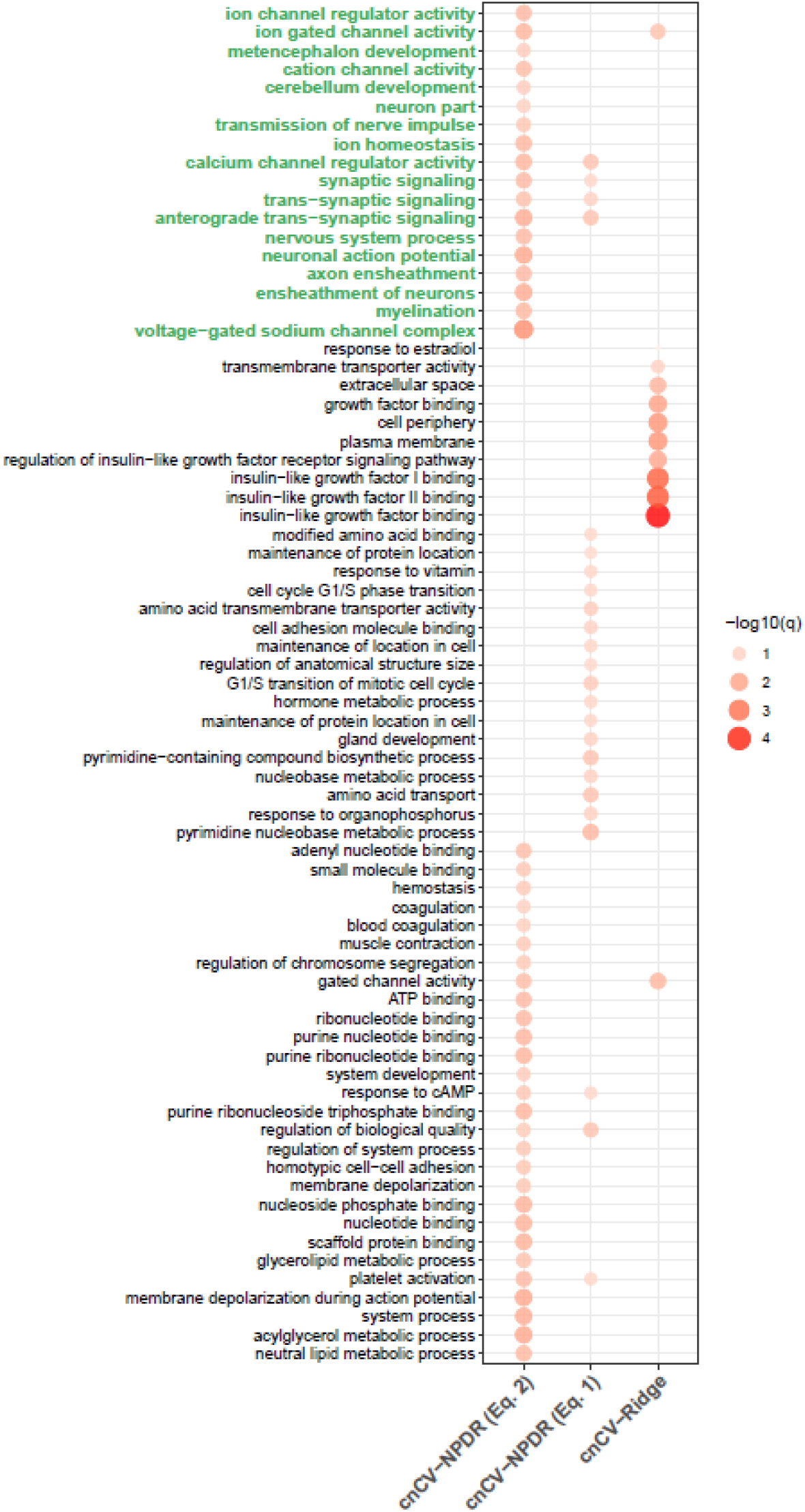
Comparison of gene ontology terms for MDD-associated genes detected using cnCV with importance scores calculated by NPDR (Eq. 1), NPDR (Eq. 2), and Ridge Regression. Point color and size is proportional to the -log10 False Discovery Rate q-value, denoted by -log10(q). Gene ontology (GO) terms are on the y-axis and feature selection methods are on the x-axis. GO terms with clear neurological relevance are highlighted in green; these include GO terms tracing back to the parent terms ‘nervous system process’ or ‘nervous system development’ and terms that are clearly specific to neuronal function

**Table 1.**
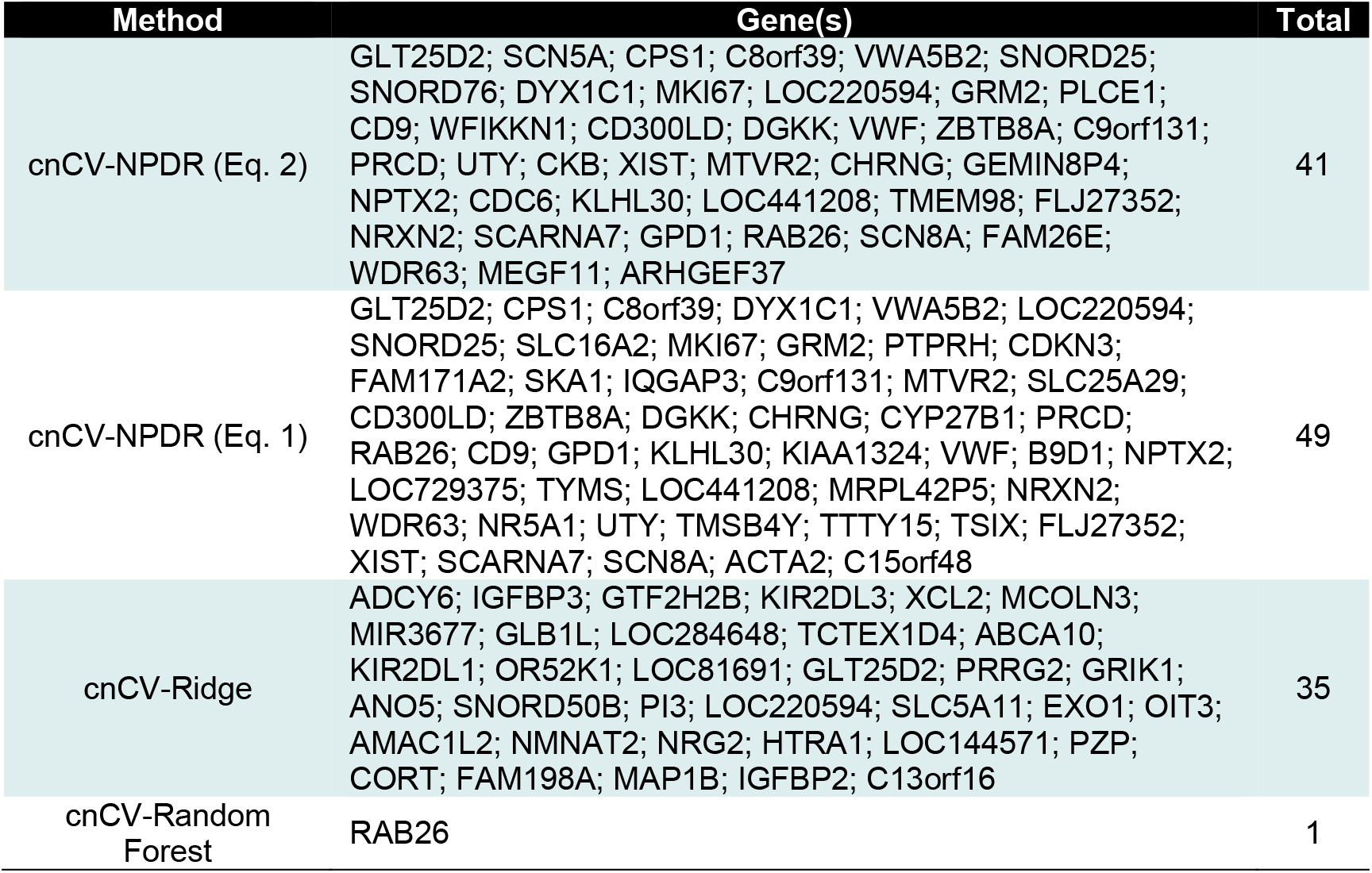
Genes detected by cnCV using NPDR with non-adjusted fixed-k (Eq. 1), NPDR with minority-class-k (Eq. 2), random forest, and ridge regression for relevance to MDD.

**Table 2.** Gene ontology terms for cnCV using NPDR with non-adjusted fixed-k (Eq. 1), NPDR with minority-class-k (Eq. 2), and ridge regression.

| Method(s) | GO Term | q-value<sup>†</sup><br>(FDR) | Genes<sup>‡</sup> | Size<br>†† |
| --- | --- | --- | --- | --- |
| cnCV-NPDR (Eq. 2) | platelet activation | **0.0293** | DGKK; VWF; CD9 | 158 |
| cnCV-NPDR (Eq. 1) |  | 0.1246 | DGKK; VWF; CD9 |  |
| cnCV-NPDR (Eq. 2) | anterograde trans-synaptic signaling | **0.0151** | RAB26; NRXN2; CHRNG; GRM2; NPTX2 | 686 |
| cnCV-NPDR (Eq. 1) |  | **0.0452** | RAB26; NRXN2; CHRNG; GRM2; NPTX2 |  |
| cnCV-NPDR (Eq. 2) | trans-synaptic signaling | **0.0422** | RAB26; NRXN2; CHRNG; GRM2; NPTX2 | 693 |
| cnCV-NPDR (Eq. 1) |  | 0.0927 | RAB26; NRXN2; CHRNG; GRM2; NPTX2 |  |
| cnCV-NPDR (Eq. 2) | synaptic signaling | **0.0293** | RAB26; NRXN2; CHRNG; GRM2; NPTX2 | 702 |
| cnCV-NPDR (Eq. 1) |  | 0.1246 | RAB26; NRXN2; CHRNG; GRM2; NPTX2 |  |
| cnCV-NPDR (Eq. 2) | calcium channel regulator activity | **0.0238** | NRXN2; GRM2 | 47 |
| cnCV-NPDR (Eq. 1) |  | **0.0464** | NRXN2; GRM2 |  |
| cnCV-NPDR (Eq. 2) | regulation of biological quality | 0.0725 | CKB; SCN8A; NRXN2; CD9; DGKK; PLCE1; CHRNG; VWF; SCN5A; CPS1; GRM2 | 4060 |
| cnCV-NPDR (Eq. 1) |  | **0.0468** | ACTA2; TMSB4Y; SLC16A2; SCN8A; NRXN2; CD9; DGKK; CYP27B1; NR5A1; VWF; IQGAP3; CPS1; GRM2; CHRNG |  |
| cnCV-NPDR (Eq. 2) | response to cAMP | 0.0617 | GPD1; CPS1 | 94 |
| cnCV-NPDR (Eq. 1) |  | 0.1294 | GPD1; CPS1 |  |
| cnCV-NPDR (Eq. 2) | ion gated channel activity | **0.0238** | SCN5A; SCN8A; CHRNG | 330 |
| cnCV-Ridge |  | **0.0429** | GRIK1; MCOLN3; ANO5 |  |
| cnCV-NPDR (Eq. 2) | gated channel activity | **0.0363** | SCN5A; SCN8A; CHRNG | 340 |
| cnCV-Ridge |  | **0.0241** | GRIK1; MCOLN3; ANO5 |  |
| cnCV-NPDR (Eq. 2) | voltage-gated sodium channel complex | **0.0041** | SCN5A; SCN8A | 17 |
| cnCV-NPDR (Eq. 2) | myelination | **0.0277** | TMEM98; SCN8A; CD9 | 123 |
| cnCV-NPDR (Eq. 2) | ensheathment of neurons | **0.0149** | TMEM98; SCN8A; CD9 | 125 |
| cnCV-NPDR (Eq. 2) | axon ensheathment | **0.0293** | TMEM98; SCN8A; CD9 | 125 |
| cnCV-NPDR (Eq. 2) | neutral lipid metabolic process | **0.0277** | DGKK; PLCE1; CPS1 | 135 |
| cnCV-NPDR (Eq. 2) | acylglycerol metabolic process | **0.0132** | DGKK; PLCE1; CPS1 | 135 |
| cnCV-NPDR (Eq. 2) | system process | **0.0149** | TMEM98; SCN8A; NRXN2; PRCD; PLCE1; NPTX2; SCN5A; CPS1; CHRNG | 2174 |
| cnCV-NPDR (Eq. 2) | neuronal action potential | **0.0132** | SCN5A; SCN8A | 33 |
| cnCV-NPDR (Eq. 2) | membrane depolarization during action potential | **0.0132** | SCN5A; SCN8A | 35 |
| cnCV-NPDR (Eq. 2) | nervous system process | **0.0293** | TMEM98; SCN8A; NRXN2; PRCD; CHRNG; SCN5A; NPTX2 | 1429 |
| cnCV-NPDR (Eq. 2) | glycerolipid metabolic process | **0.0427** | PLCE1; GPD1; CPS1; DGKK | 435 |
| cnCV-NPDR (Eq. 2) | scaffold protein binding | **0.0196** | SCN5A; GRM2 | 58 |
| cnCV-NPDR (Eq. 2) | ion homeostasis | **0.0252** | SCN5A; PLCE1; CPS1; GRM2; CKB | 810 |
| cnCV-NPDR (Eq. 2) | transmission of nerve impulse | 0.0574 | SCN5A; SCN8A | 67 |
| cnCV-NPDR (Eq. 2) | nucleotide binding | **0.0196** | MKI67; RAB26; CKB; SCN8A; DGKK; GPD1; CPS1; CDC6 | 2148 |
| cnCV-NPDR (Eq. 2) | nucleoside phosphate binding | **0.0196** | MKI67; RAB26; CKB; SCN8A; DGKK; GPD1; CPS1; CDC6 | 2149 |
| cnCV-NPDR (Eq. 2) | neuron part | 0.0922 | RAB26; CKB; SCN8A; NRXN2; PRCD; CHRNG; GRM2 | 1747 |
| cnCV-NPDR (Eq. 2) | membrane depolarization | 0.0586 | SCN5A; SCN8A | 84 |
| cnCV-NPDR (Eq. 2) | homotypic cell-cell adhesion | 0.0586 | CD9; MEGF11 | 86 |
| cnCV-NPDR (Eq. 2) | regulation of system process | 0.0586 | TMEM98; SCN5A; PLCE1; NPTX2 | 576 |
| cnCV-NPDR (Eq. 2) | purine ribonucleoside triphosphate binding | **0.0238** | CKB; SCN8A; NRXN2; CD9; DGKK; PLCE1; CHRNG; VWF; SCN5A; CPS1; GRM2 | 4060 |
| cnCV-NPDR (Eq. 2) | cerebellum development | 0.0796 | MKI67; RAB26; CKB; SCN8A; DGKK; CPS1; CDC6 | 1829 |
| cnCV-NPDR (Eq. 2) | system development | 0.0796 | SCN5A; CKB | 94 |
| cnCV-NPDR (Eq. 2) | purine ribonucleotide binding | **0.0238** | MKI67; RAB26; CKB; SCN8A; DGKK; CPS1; CDC6 | 1896 |
| cnCV-NPDR (Eq. 2) | purine nucleotide binding | **0.0238** | MKI67; RAB26; CKB; SCN8A; DGKK; CPS1; CDC6 | 1911 |
| cnCV-NPDR (Eq. 2) | ribonucleotide binding | **0.0299** | MKI67; RAB26; CKB; SCN8A; DGKK; CPS1; CDC6 | 1912 |
| cnCV-NPDR (Eq. 2) | cation channel activity | **0.0363** | SCN5A; SCN8A; CHRNG | 317 |
| cnCV-NPDR (Eq. 2) | metencephalon development | 0.0796 | SCN5A; CKB | 101 |
| cnCV-NPDR (Eq. 2) | ATP binding | **0.0299** | MKI67; CKB; SCN8A; DGKK; CPS1; CDC6 | 1486 |
| cnCV-NPDR (Eq. 2) | regulation of chromosome segregation | 0.0675 | MKI67; CDC6 | 111 |
| cnCV-NPDR (Eq. 2) | muscle contraction | 0.0675 | SCN5A; PLCE1; CHRNG | 343 |
| cnCV-NPDR (Eq. 2) | blood coagulation | 0.0856 | DGKK; VWF; CD9 | 344 |
| cnCV-NPDR (Eq. 2) | coagulation | 0.0905 | DGKK; VWF; CD9 | 348 |
| cnCV-NPDR (Eq. 2) | hemostasis | 0.0675 | DGKK; VWF; CD9 | 348 |
| cnCV-NPDR (Eq. 2) | small molecule binding | 0.0606 | MKI67; RAB26; CKB; SCN8A; DGKK; GPD1; CPS1; CDC6 | 2565 |
| cnCV-NPDR (Eq. 2) | ion channel regulator activity | **0.0300** | NRXN2; GRM2 | 120 |
| cnCV-NPDR (Eq. 2) | adenyl nucleotide binding | **0.0363** | MKI67; CKB; SCN8A; DGKK; CPS1; CDC6 | 1559 |
| cnCV-NPDR (Eq. 1) | pyrimidine nucleobase metabolic process | **0.0255** | TYMS; CPS1 | 21 |
| cnCV-NPDR (Eq. 1) | response to organophosphorus | 0.0915 | TYMS; GPD1; CPS1 | 131 |
| cnCV-NPDR (Eq. 1) | amino acid transport | **0.0447** | SLC25A29; SLC16A2; GRM2 | 155 |
| cnCV-NPDR (Eq. 1) | nucleobase metabolic process | 0.0915 | TYMS; CPS1 | 40 |
| cnCV-NPDR (Eq. 1) | pyrimidine-containing compound biosynthetic process | **0.0452** | TYMS; CPS1 | 53 |
| cnCV-NPDR (Eq. 1) | gland development | 0.0927 | TYMS; CPS1; NR5A1; IQGAP3 | 414 |
| cnCV-NPDR (Eq. 1) | maintenance of protein location in cell | 0.1246 | TMSB4Y; NR5A1 | 63 |
| cnCV-NPDR (Eq. 1) | hormone metabolic process | 0.1162 | SLC16A2; CYP27B1; NR5A1 | 244 |
| cnCV-NPDR (Eq. 1) | G1/S transition of mitotic cell cycle | 0.0658 | CDKN3; TYMS; IQGAP3 | 246 |
| cnCV-NPDR (Eq. 1) | regulation of anatomical structure size | 0.1365 | ACTA2; TMSB4Y; CPS1; IQGAP3 | 492 |
| cnCV-NPDR (Eq. 1) | maintenance of location in cell | 0.1162 | TMSB4Y; NR5A1 | 82 |
| cnCV-NPDR (Eq. 1) | cell adhesion molecule binding | 0.0967 | VWF; NRXN2; CD9; PTPRH | 516 |
| cnCV-NPDR (Eq. 1) | amino acid transmembrane transporter activity | 0.0758 | SLC25A29; SLC16A2 | 84 |
| cnCV-NPDR (Eq. 1) | cell cycle G1/S phase transition | 0.1294 | CDKN3; TYMS; IQGAP3 | 266 |
| cnCV-NPDR (Eq. 1) | response to vitamin | 0.1294 | TYMS; CYP27B1 | 88 |
| cnCV-NPDR (Eq. 1) | maintenance of protein location | 0.1594 | TMSB4Y; NR5A1 | 93 |
| cnCV-NPDR (Eq. 1) | modified amino acid binding | 0.1341 | TYMS; CPS1 | 95 |
| cnCV-Ridge | insulin-like growth factor binding | **0.0001** | IGFBP2; IGFBP3; HTRA1 | 29 |
| cnCV-Ridge | insulin-like growth factor II binding | **0.0004** | IGFBP2; IGFBP3 | 8 |
| cnCV-Ridge | insulin-like growth factor I binding | **0.0006** | IGFBP2; IGFBP3 | 13 |
| cnCV-Ridge | regulation of insulin-like growth factor receptor signaling pathway | **0.0121** | IGFBP2; IGFBP3 | 22 |
| cnCV-Ridge | plasma membrane | **0.0053** | KIR2DL3; ADCY6; NRG2; PRRG2; OR52K1; GRIK1; EXO1; MAP1B; IGFBP2; KIR2DL1; SLC5A11; MCOLN3; ANO5; PI3; HTRA1 | 5769 |
| cnCV-Ridge | cell periphery | **0.0053** | KIR2DL3; ADCY6; NRG2; PRRG2; OR52K1; GRIK1; EXO1; MAP1B; IGFBP2; KIR2DL1; SLC5A11; MCOLN3; ANO5; PI3; HTRA1 | 5879 |
| cnCV-Ridge | growth factor binding | **0.0094** | IGFBP2; IGFBP3; HTRA1 | 138 |
| cnCV-Ridge | extracellular space | **0.0220** | IGFBP2; NRG2; PRRG2; XCL2; PZP; CORT; IGFBP3; PI3; HTRA1; OIT3 | 3414 |
| cnCV-Ridge | transmembrane transporter activity | 0.0999 | ABCA10; GRIK1; SLC5A11; MCOLN3; ANO5 | 1067 |
| cnCV-Ridge | response to estradiol | 0.4770 | IGFBP2; MAP1B | 125 |

<sup>†</sup>P-value for over-representation calculated from hypergeometric test, based on the number of genes in predefined set and user-input set. Multiple testing correction performed using false discovery rate (q).
<sup>‡</sup>Genes overlapping between user-input set and predefined set.
<sup>§</sup>Size, the number of genes in the predefined gene set.

## 4 DISCUSSION

In this study, we presented novel neighborhood algorithms to optimize performance in detecting features relevant to dichotomous outcomes across multiple nearest-neighbor feature selection methods. We assessed performance using realistic high-dimensional simulated data with concentration on class imbalance. We showed that the optimal neighborhood for detecting relevant main effects and interaction effects depends heavily on the degree of class imbalance, favoring a shift toward reduced neighborhood sizes as imbalance becomes more extreme. Across a variety of different simulation scenarios, our new imbalance-adjusted fixed-k formulations led to significantly higher feature-based Area Under the Precision-Recall Curve and Matthews Correlation Coefficient. This improvement was seen relative not only to their non-adjusted predecessors but also relative to adaptive-radius methods and relative to random forest and ridge regression. We applied our minority-class-k to create a constrained k-grid, providing adaptive-k alternatives that led to the best performance overall for each nearest-neighbor scoring approach. Applying consensus-features nested cross-validation to real RNA-Seq data from a study of Major Depressive Disorder, genes detected using minority-class-k with NPDR feature scoring were associated with numerous pathways of particular relevance to the nervous system and neural signalling, more so than other feature scoring approaches.

Our minority-class-k performed significantly better than the MultiSURF adaptive-radius [1] within the context of NPDR, and hit-miss-k performed at least as well as MultiSURF adaptive-radius with respect to ReliefF and the pseudo t-test STIR. Although not shown here, we have observed significantly distinct feature rankings for NPDR, ReiefF, and STIR using the same neighborhood algorithm. For example, using the same value of fixed-k, feature rankings may differ dramatically between NPDR and ReliefF. The separate hit-miss-k that allowed for ReliefF and STIR to perform well on imbalanced data did not translate as well to NPDR; the single, minority-class-k performed better than hit-miss-k within the context of NPDR. This implied that NPDR performed better with smaller neighborhood sizes than ReliefF or STIR on imbalanced data, given that minority-class-k is smaller in magnitude than cumulative neighbors given by hit-miss-k when classes are imbalanced. Contrary to hit-miss-k, the single value of fixed-k used by NPDR allowed for stochasticity in the number of within- and between-class nearest neighbors, potentially being another contributing factor to the superiority of minority-class-k over hit-miss-k for NPDR.

In addition to comparing the quality of raw feature scores for each method, we also compared performance within consensus-features nested cross-validation [12] since filter-based approached are typically wrapped within cross-validation to limit over-fitting. Because consensus-features nested cross-validation only selects features that are among the highest in rank across all inner training folds, the variability of feature rankings for a given scoring method is an additional parameter for optimizing performance. Although our new constrained, imbalance-adjusted k-grid allowed variable-wise optimized k to achieve the best overall performance with respect to raw feature scores, this advantage was mitigated within consensus-features nested cross-validation; our imbalance-adjusted fixed-k approaches yielded similar feature-based Matthews Correlation Coefficient, precision, and recall across all simulation scenarios, suggesting fixed-k may yield less variable feature rankings across training folds. Thus, our imbalance-adjusted fixed-k approaches may be ideal for the sake of efficiency within the nested training procedure. However, our results suggested that methods like NPDR and STIR that allow for the quantification of statistical significance for feature importance may be optimized using the constrained, imbalance-adjusted variable-wise optimized k, minority-class-k for NPDR and hit-miss-k for STIR.

Random forest and penalized regression approaches, such as ridge regression, are commonly applied to high-dimensional biomedical data, considerably more often than nearest-neighbor feature selection. We compared the performance of NPDR, ReliefF, and STIR to random forest and ridge regression, finding that these nearest-neighbor feature selection approaches were all superior using our imbalance-adjusted methods. One exception occurred in simulation scenarios where functional features were predominantly composed of main effects, not interaction effects; ridge regression performed predictably well in this scenario, achieving the best performance particularly in the context of group imbalance. This was not surprising since ridge regression is explicitly designed for detecting minimally redundant main effects, whereas our neighborhood parameterization of each nearest-neighbor feature scoring method is meant to balance detection of interaction effects and main effects.

Feature selection for an RNA-Seq study of MDD revealed that the majority of the enriched biological pathways were derived from genes detected using NPDR with minority-class-k. NPDR with minority-class-k was the only feature scoring method having gene ontology terms linked directly to the nervous system, but ridge regression attained the strongest enrichment scores overall, specifically detecting genes in the insulin-like growth factor binding pathways that are known to be associated with Major Depressive Disorder [13–16]. Ridge regression also achieved the highest concordance between gene sets detected on the full (balanced) and sub-sampled (imbalanced) RNA-Seq data sets, respectively; NPDR showed very little concordance in the same comparison, suggesting a greater degree of variability of gene rankings across inner training folds. RAB26 was the only consensus gene detected using random forest, indicating significant variability of gene rankings across inner training folds, however RAB26 is highly expressed in the brain [17, 18] and RAB proteins are crucial to synaptic function [19]; RAB26 was also detected using NPDR both with minority-class-k and non-adjusted fixed-k. Cumulatively, these results suggest that consensus-features nested cross-validation using NPDR with minority-class-k was the best approach overall for detecting genes of likely relevance to Major Depressive Disorder in the presence of class imbalance.

## 5 CONCLUSION

In summary, this study addressed the problem of optimizing fixed-k neighborhoods across the spectrum of nearest-neighbor feature scoring algorithms for binary classification, particularly in the presence of group imbalance. Our imbalance-adjusted neighborhood algorithms achieved superior performance, not only when compared to non-adjusted nearest-neighbor approaches but also when compared to random forest and ridge regression, in a wide variety of data simulations having both interaction effects and main effects. The combination of superior performance on simulated data and the identification of neurologically linked genes associated with Major Depression in real RNA-Seq data demonstrated the validity of our approach and its ability to identify relevant predictors of complex diseases.

## ACKNOWLEDGEMENTS

This work was supported by the National Institute of Health [GM121312, GM103456 to B.M.].

## SUPPLEMENTARY INFORMATION

Supplementary Materials.pdf

