## Supplementary Materials for "Multivariate optimization of k for k-nearest-neighbor feature selection with dichotomous outcomes: complex associations, class imbalance, and application to RNA-Seq in Major Depressive Disorder"

#### Outline

1. Feature selection performance comparisons: equal main effect and interaction effect
   1.1 Comparing imbalance-adjusted fixed-k, VWOK, and kPCA
   1.2 Comparing imbalance-adjusted fixed-k, regular fixed-k, and MultiSURF
   1.3 Comparing imbalance-adjusted fixed-k, random forest, and ridge regression

2. Feature selection performance comparisons: 75% interaction effect/25% main effect
   2.1 Comparing imbalance-adjusted fixed-k, VWOK, and kPCA
   2.2 Comparing imbalance-adjusted fixed-k, regular fixed-k, and MultiSURF
   2.3 Comparing imbalance-adjusted fixed-k, random forest, and ridge regression

3. Feature selection performance comparisons: 25% interaction effect/75% main effect
   3.1 Comparing imbalance-adjusted fixed-k, VWOK, and kPCA
   3.2 Comparing imbalance-adjusted fixed-k, regular fixed-k, and MultiSURF
   3.3 Comparing imbalance-adjusted fixed-k, random forest, and ridge regression

4 Feature selection performance comparisons within consensus-features nested cross-validation (cnCV): equal main effect and interaction effect

   4.1 Comparing imbalance-adjusted fixed-k, VWOK, and kPCA
   4.2 Comparing imbalance-adjusted fixed-k, regular fixed-k, and MultiSURF
   4.3 Comparing imbalance-adjusted fixed-k, random forest, and ridge regression

5 Feature selection performance comparisons within consensus-features nested cross-validation (cnCV): 75% interaction effect/25% main effect

   5.1 Comparing imbalance-adjusted fixed-k, VWOK, and kPCA
   5.2 Comparing imbalance-adjusted fixed-k, regular fixed-k, and MultiSURF
   5.3 Comparing imbalance-adjusted fixed-k, random forest, and ridge regression

6 Feature selection performance comparisons within consensus-features nested cross-validation (cnCV): 25% interaction effect/75% main effect

   6.1 Comparing imbalance-adjusted fixed-k, VWOK, and kPCA
   6.2 Comparing imbalance-adjusted fixed-k, regular fixed-k, and MultiSURF
   6.3 Comparing imbalance-adjusted fixed-k, random forest, and ridge regression

### 1 Feature selection performance comparisons: equal main effect and interaction effect

#### 1.1 Comparing imbalance-adjusted fixed-k, VWOK, and kPCA

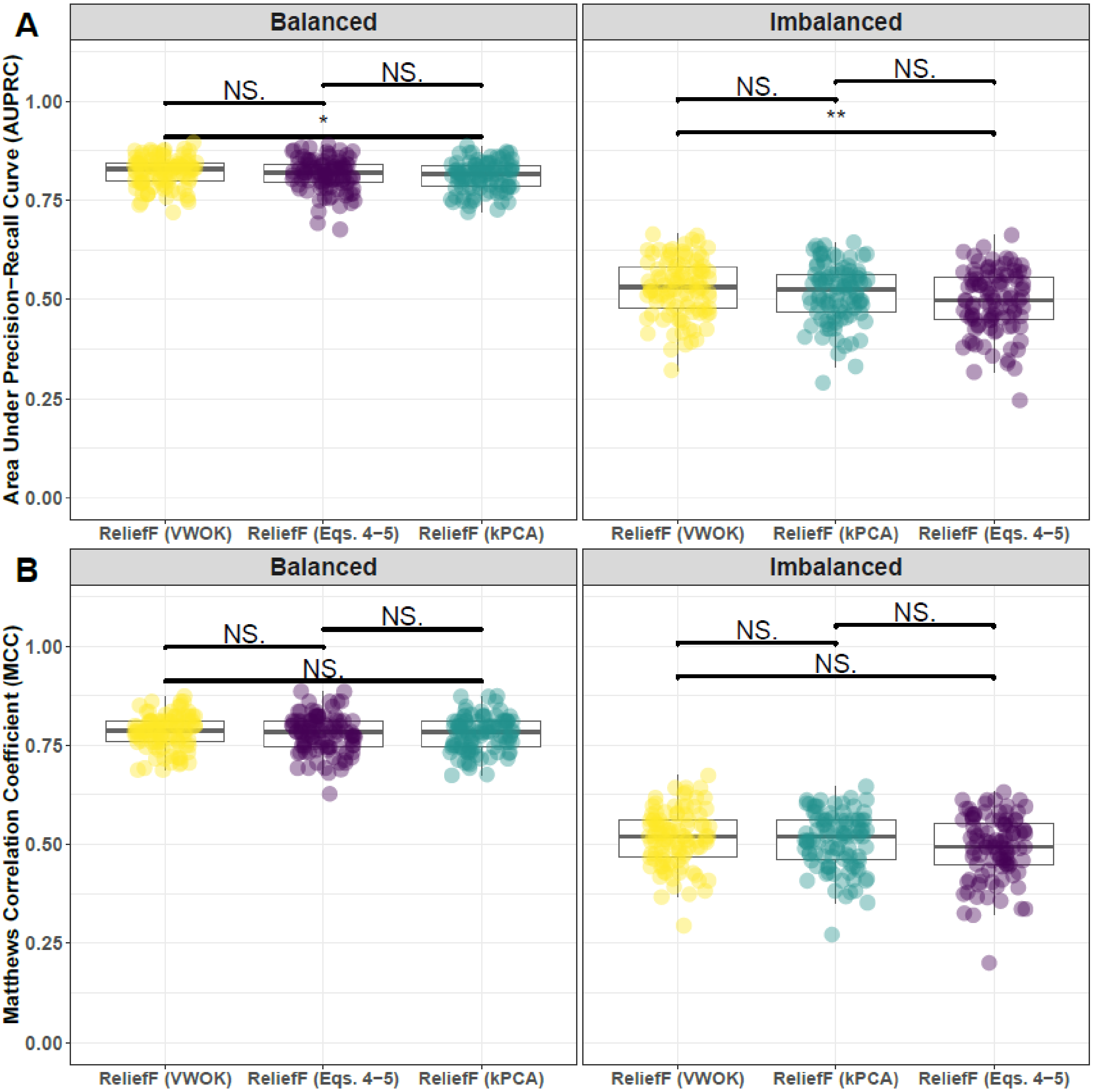


**Supplementary Fig. 1. Performance comparison for hit-miss-k (Eqs. 4 – 5), VWOK (Eqs. 6 – 7), and kPCA with ReliefF feature scoring.** Performance of feature selection was measured for 100 simulation replicates. Each simulated data set had $m = 100$ instances and $p = 1000$ features with 100 functional. Functional features included 50 that with main effect ($\text{bias}_{\text{main}} = 0.8$) only and the remaining 50 were involved in network interactions ($\text{bias}_{\text{int}} = 0.4$) and had no main effect. Imbalanced simulations had class ratio of 25:75 (cases:controls). (**A**) Area Under Precision-Recall Curve (AUPRC) for each method, sorted by decreasing mean AUPRC. (**B**) Matthews Correlation Coefficient (MCC) for each method, sorted by decreasing mean MCC. Comparisons were made with Mann-Whitney U test ($* \; P < 0.05$ and $** \; P < 0.01$)

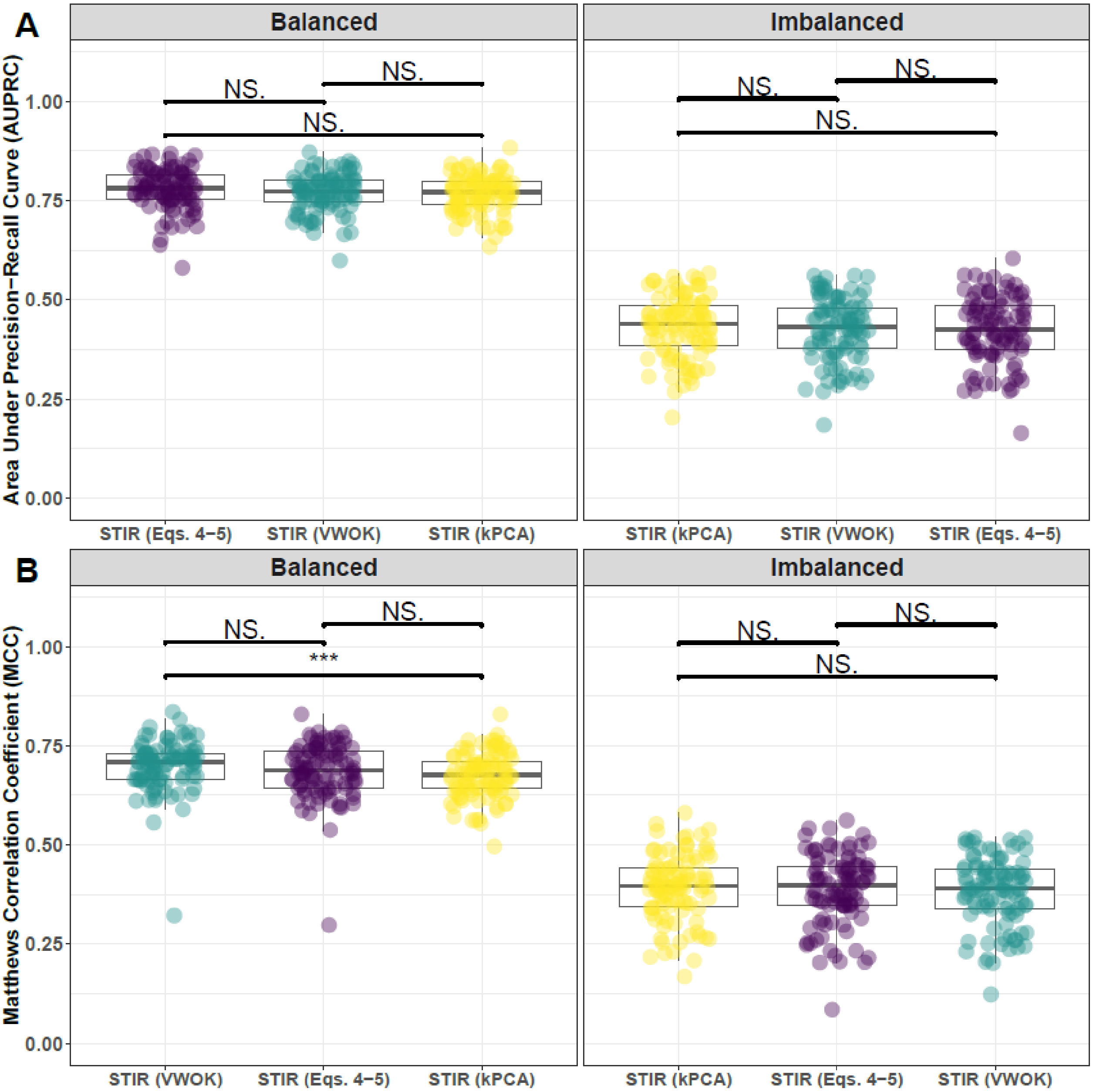


**Supplementary Fig. 2. Performance comparison for hit-miss-k (Eqs. 4 – 5), VWOK (Eqs. 6 – 7), and kPCA with STIR feature scoring.** Performance of feature selection was measured for 100 simulation replicates. Each simulated data set had $m = 100$ instances and $p = 1000$ features with 100 functional. Functional features included 50 that with main effect ($\text{bias}_{\text{main}} = 0.8$) only and the remaining 50 were involved in network interactions ($\text{bias}_{\text{int}} = 0.4$) and had no main effect. Imbalanced simulations had class ratio of 25:75 (cases:controls). (**A**) Area Under Precision-Recall Curve (AUPRC) for each method, sorted by decreasing mean AUPRC. (**B**) Matthews Correlation Coefficient (MCC) for each method, sorted by decreasing mean MCC. Comparisons were made with Mann-Whitney U test ($* \, P < 0.05$ and $** \, P < 0.01$).

#### 1.2 Comparing imbalance-adjusted fixed-k, regular fixed-k, and MultiSURF

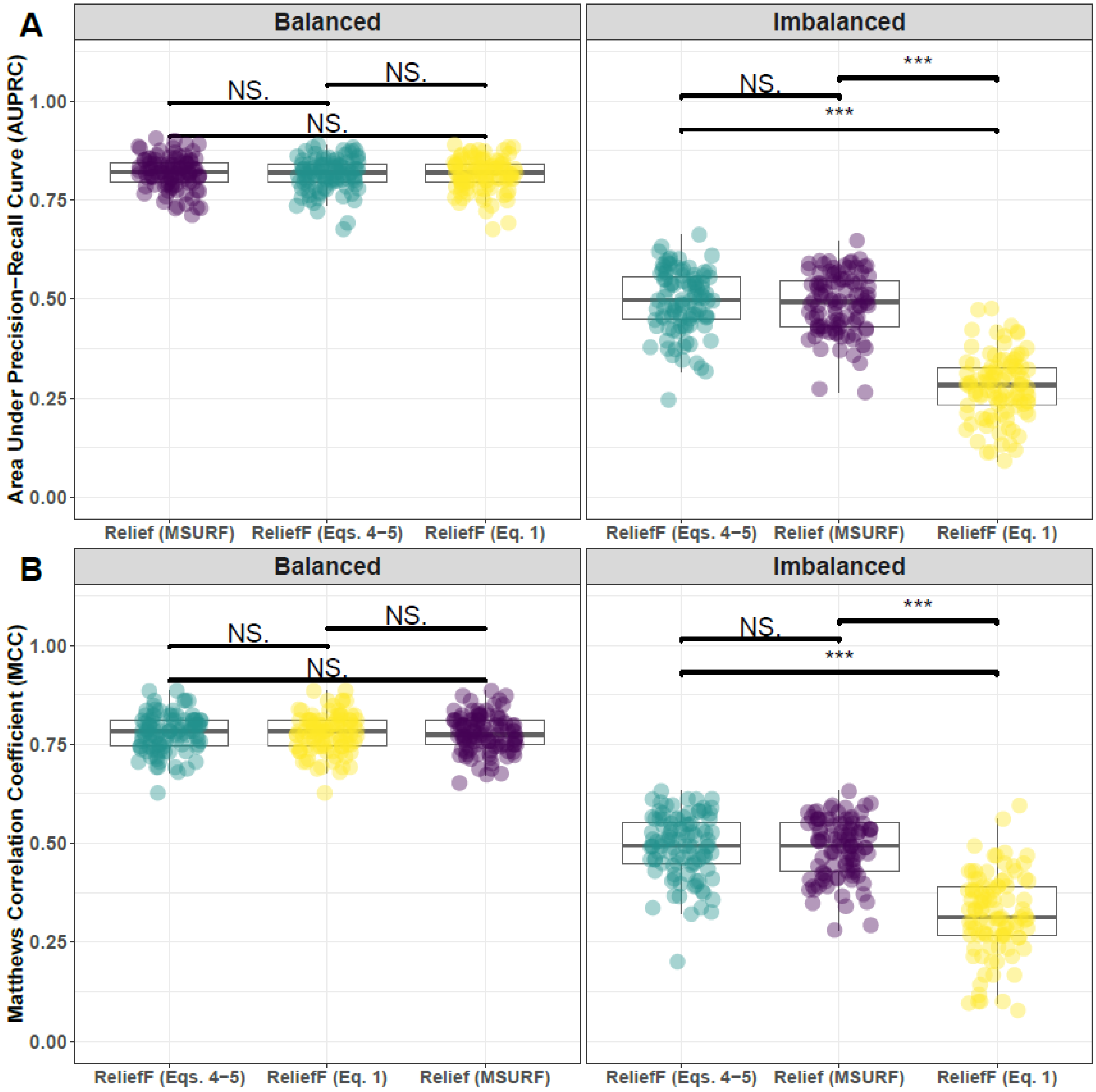


**Supplementary Fig. 4. Performance comparison of ReliefF with hit-miss-k (Eqs. 4 – 5), ReliefF with non-adjusted fixed-k (Eq. 1), and MultiSURF.** Performance of feature selection was measured for 100 simulation replicates. Each simulated data set had $m = 100$ instances and $p = 1000$ features with 100 functional. Functional features included 50 that with main effect ($\text{bias}_{\text{main}} = 0.8$) only and the remaining 50 were involved in network interactions ($\text{bias}_{\text{int}} = 0.4$) and had no main effect. Imbalanced simulations had class ratio of 25:75 (cases:controls). (**A**) Area Under Precision-Recall Curve (AUPRC) for each method, sorted by decreasing mean AUPRC. (**B**) Matthews Correlation Coefficient (MCC) for each method, sorted by decreasing mean MCC. Comparisons were made with Mann-Whitney U test ($* \, P < 0.05$ and $*** \, P < 0.001$).

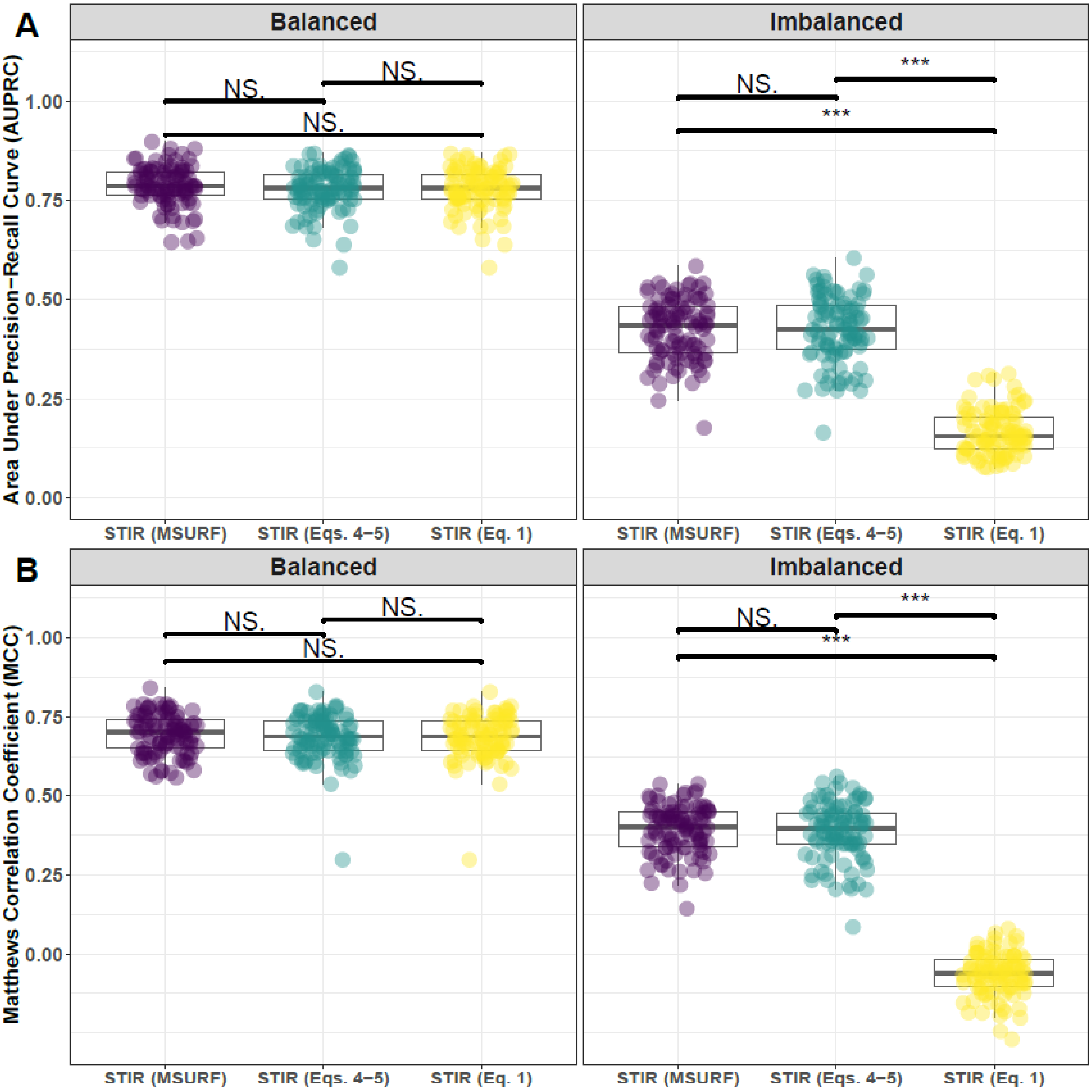


**Supplementary Fig. 5. Performance comparison of STIR with hit-miss-k (Eqs. 4 – 5), STIR with non-adjusted fixed-k (Eq. 1), and STIR (MultiSURF).** Performance of feature selection was measured for 100 simulation replicates. Each simulated data set had $m = 100$ instances and $p = 1000$ features with 100 functional. Functional features included 50 that with main effect ($\text{bias}_{\text{main}} = 0.8$) only and the remaining 50 were involved in network interactions ($\text{bias}_{\text{int}} = 0.4$) and had no main effect. Imbalanced simulations had class ratio of 25:75 (cases:controls). (**A**) Area Under Precision-Recall Curve (AUPRC) for each method, sorted by decreasing mean AUPRC. (**B**) Matthews Correlation Coefficient (MCC) for each method, sorted by decreasing mean MCC. Comparisons were made with Mann-Whitney U test ($* \, P < 0.05$ and $*** \, P < 0.001$).

#### 1.3 Comparing imbalance-adjusted fixed-k, random forest, and ridge regression

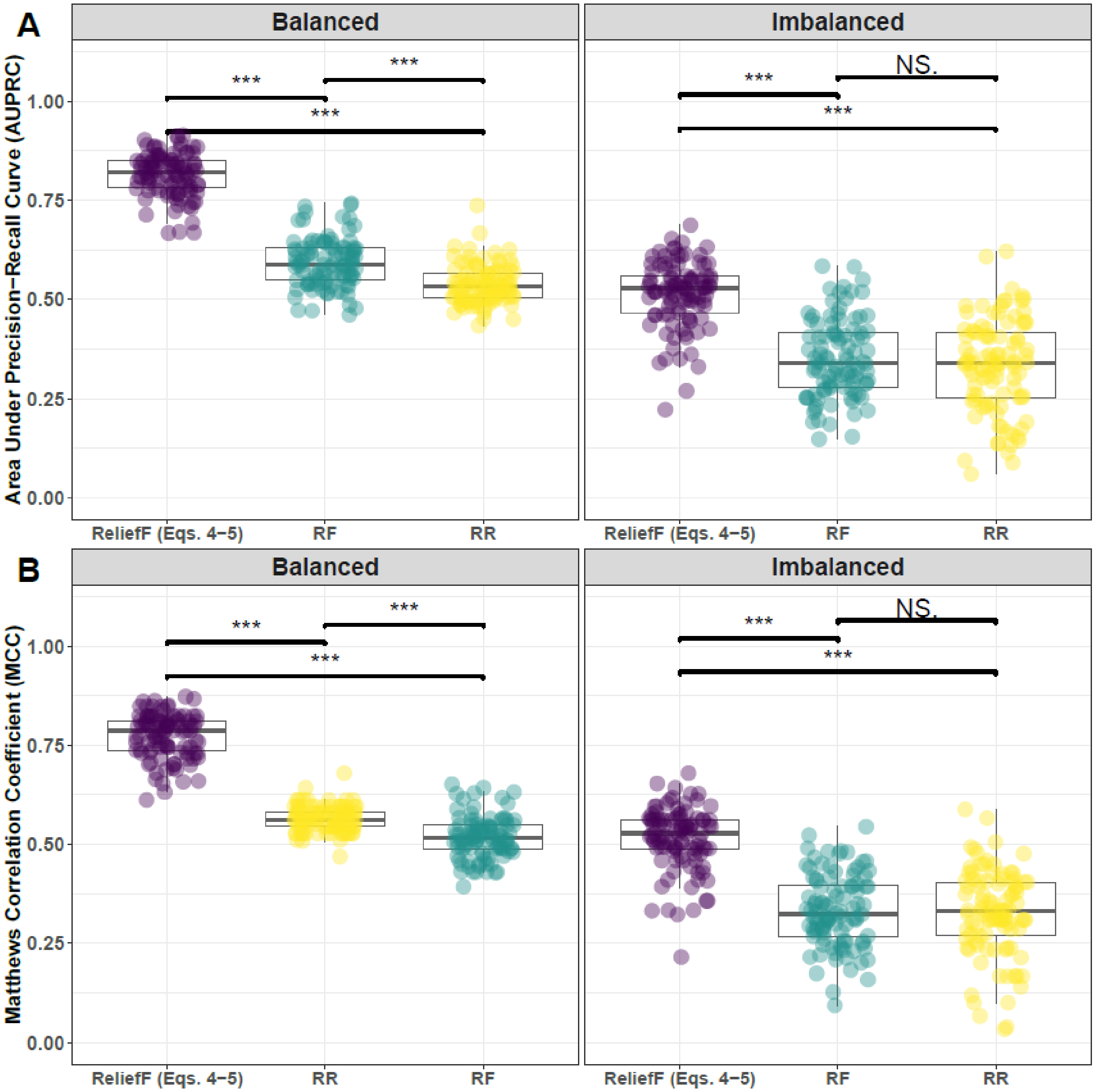


**Supplementary Fig. 6. Performance comparison of ReliefF with hit-miss-k (Eqs. 4 – 5), Random Forest (RF), and Ridge Regression (RR).** Performance of feature selection was measured for 100 simulation replicates. Each simulated data set had $m = 100$ instances and $p = 1000$ features with 100 functional. Functional features included 50 that with main effect ($\text{bias}_{\text{main}} = 0.8$) only and the remaining 50 were involved in network interactions ($\text{bias}_{\text{int}} = 0.4$) and had no main effect. Imbalanced simulations had class ratio of 25:75 (cases:controls). (**A**) Area Under Precision-Recall Curve (AUPRC) for each method, sorted by decreasing mean AUPRC. (**B**) Matthews Correlation Coefficient (MCC) for each method, sorted by decreasing mean MCC. Comparisons were made with Mann-Whitney U test ($* P < 0.05$ and $*** P < 0.001$).

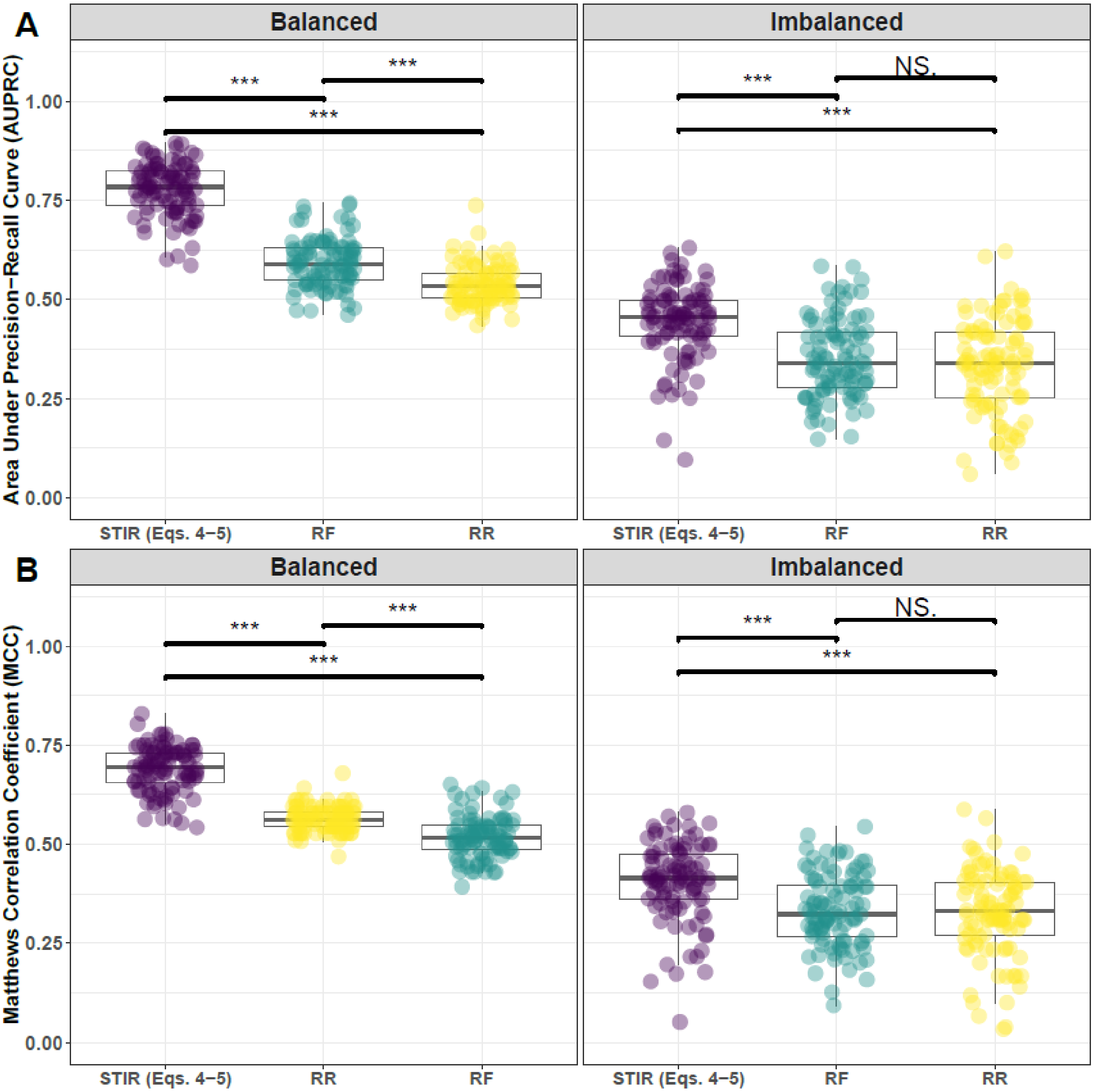


**Supplementary Fig. 7. Performance comparison of STIR with hit-miss-k (Eqs. 4 – 5), Random Forest (RF), and Ridge Regression (RR).** Performance of feature selection was measured for 100 simulation replicates. Each simulated data set had $m = 100$ instances and $p = 1000$ features with 100 functional. Functional features included 50 that with main effect ($\text{bias}_{\text{main}} = 0.8$) only and the remaining 50 were involved in network interactions ($\text{bias}_{\text{int}} = 0.4$) and had no main effect. Imbalanced simulations had class ratio of 25:75 (cases:controls). (**A**) Area Under Precision-Recall Curve (AUPRC) for each method, sorted by decreasing mean AUPRC. (**B**) Matthews Correlation Coefficient (MCC) for each method, sorted by decreasing mean MCC. Comparisons were made with Mann-Whitney U test ($* \ P < 0.05$ and $*** \ P < 0.001$)

### 2 Feature selection performance comparisons: 75% interaction effect/25% main effect

#### 2.1 Comparing imbalance-adjusted fixed-k, VWOK, and kPCA

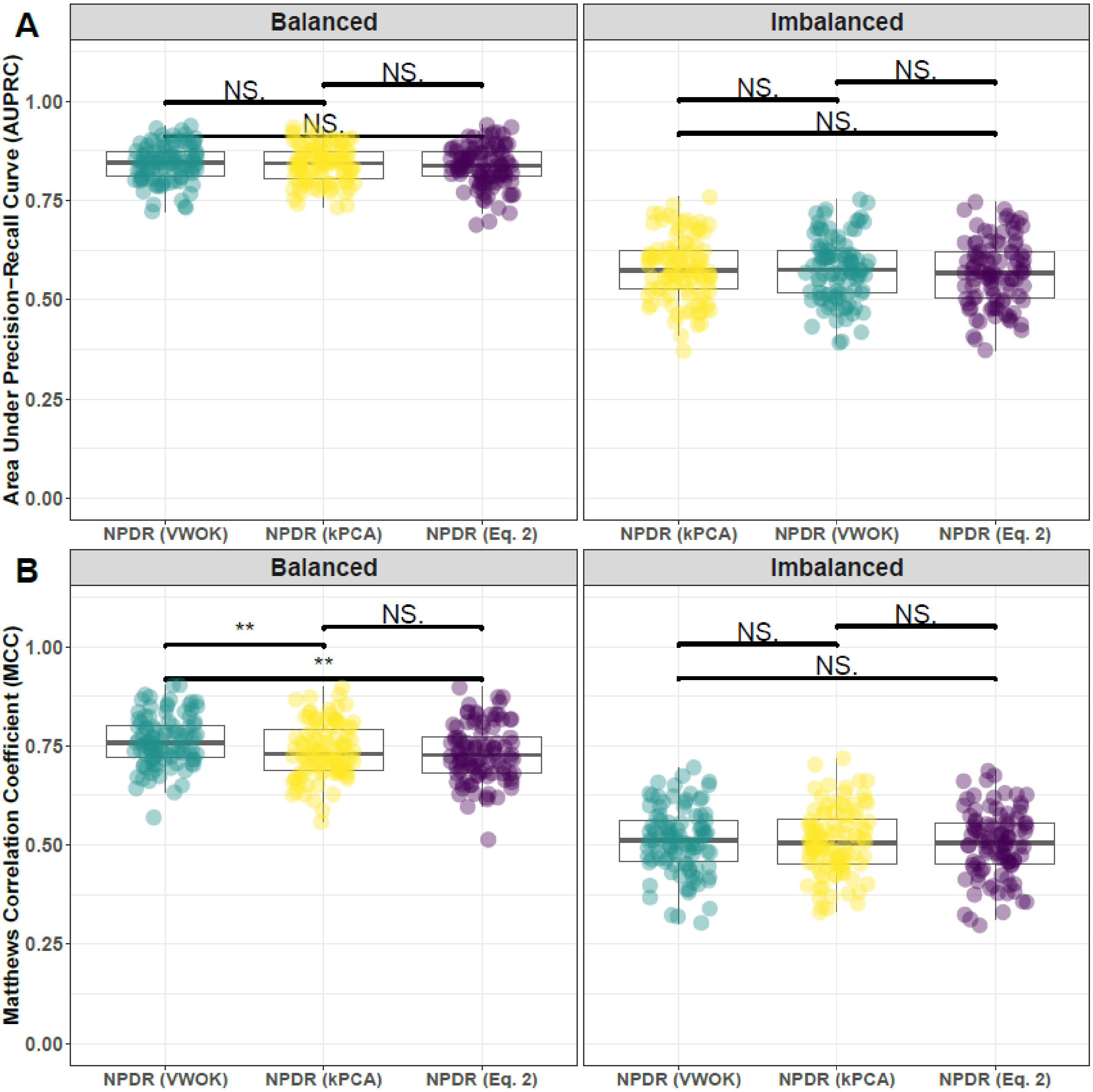


**Supplementary Fig. 8. Performance comparison for minority-class-k (Eq. 2), VWOK (Eq. 3), and kPCA with NPDR feature scoring.** Performance of feature selection was measured for 100 simulation replicates. Each simulated data set had $m = 100$ instances and $p = 1000$ features with 100 functional. Functional features included 25 that with main effect ($\text{bias}_{\text{main}} = 0.8$) only and the remaining 75 were involved in network interactions ($\text{bias}_{\text{int}} = 0.4$) and had no main effect. Imbalanced simulations had class ratio of 25:75 (cases:controls). (**A**) Area Under Precision-Recall Curve (AUPRC) for each method, sorted by decreasing mean AUPRC. (**B**) Matthews Correlation Coefficient (MCC) for each method, sorted by decreasing mean MCC. Comparisons were made with Mann-Whitney U test (NS. $P \geq 0.05$, $*\, P < 0.05$, and $**\, P < 0.01$).

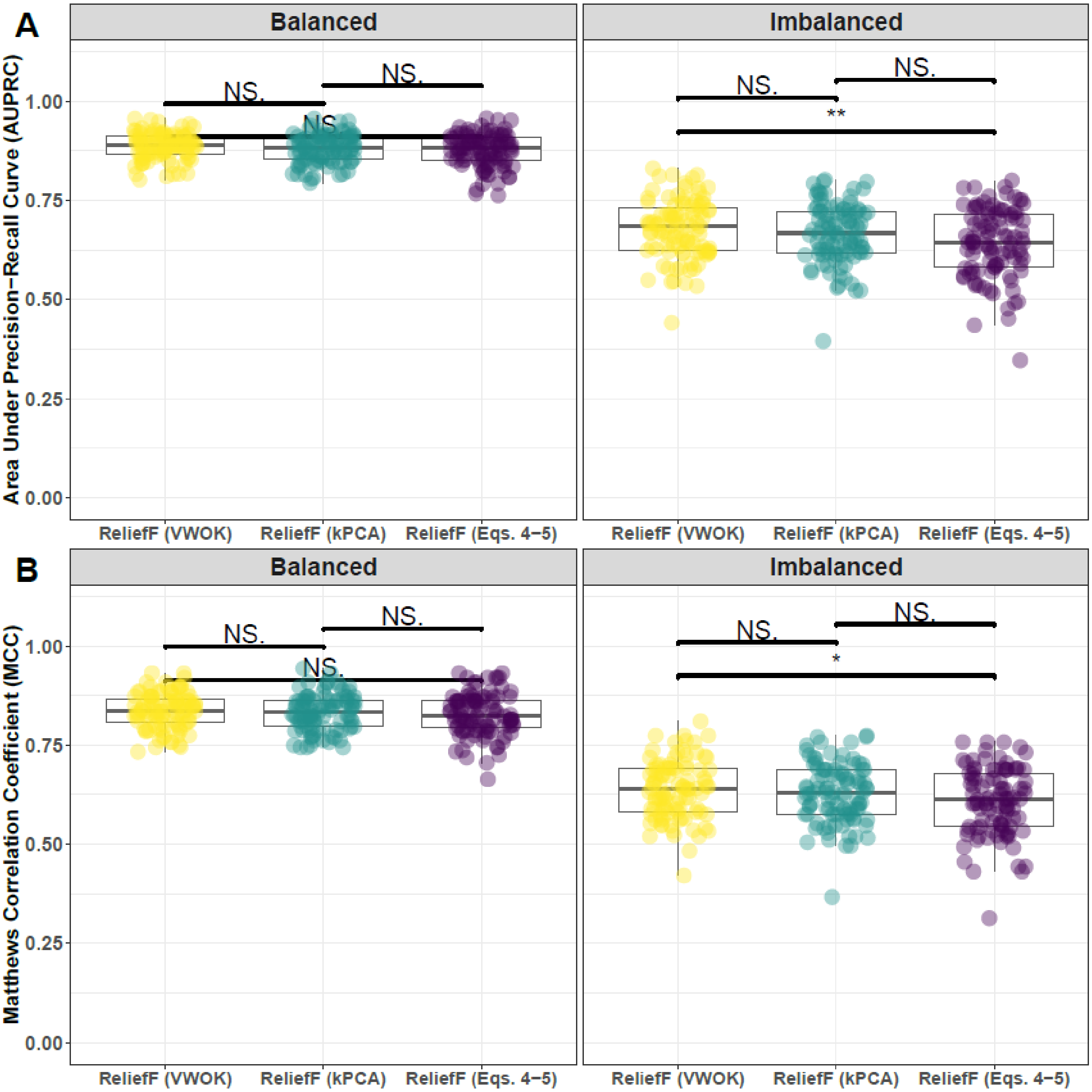


**Supplementary Fig. 9. Performance comparison for hit-miss-k (Eqs. 4 – 5), VWOK (Eqs. 6 – 7), and kPCA with ReliefF feature scoring.** Performance of feature selection was measured for 100 simulation replicates. Each simulated data set had $m = 100$ instances and $p = 1000$ features with 100 functional. Functional features included 25 that with main effect ($\text{bias}_{\text{main}} = 0.8$) only and the remaining 75 were involved in network interactions ($\text{bias}_{\text{int}} = 0.4$) and had no main effect. Imbalanced simulations had class ratio of 25:75 (cases:controls). (**A**) Area Under Precision-Recall Curve (AUPRC) for each method, sorted by decreasing mean AUPRC. (**B**) Matthews Correlation Coefficient (MCC) for each method, sorted by decreasing mean MCC. Comparisons were made with Mann-Whitney U test (NS. $P \geq 0.05$, $* \, P < 0.05$, and $** \, P < 0.01$).

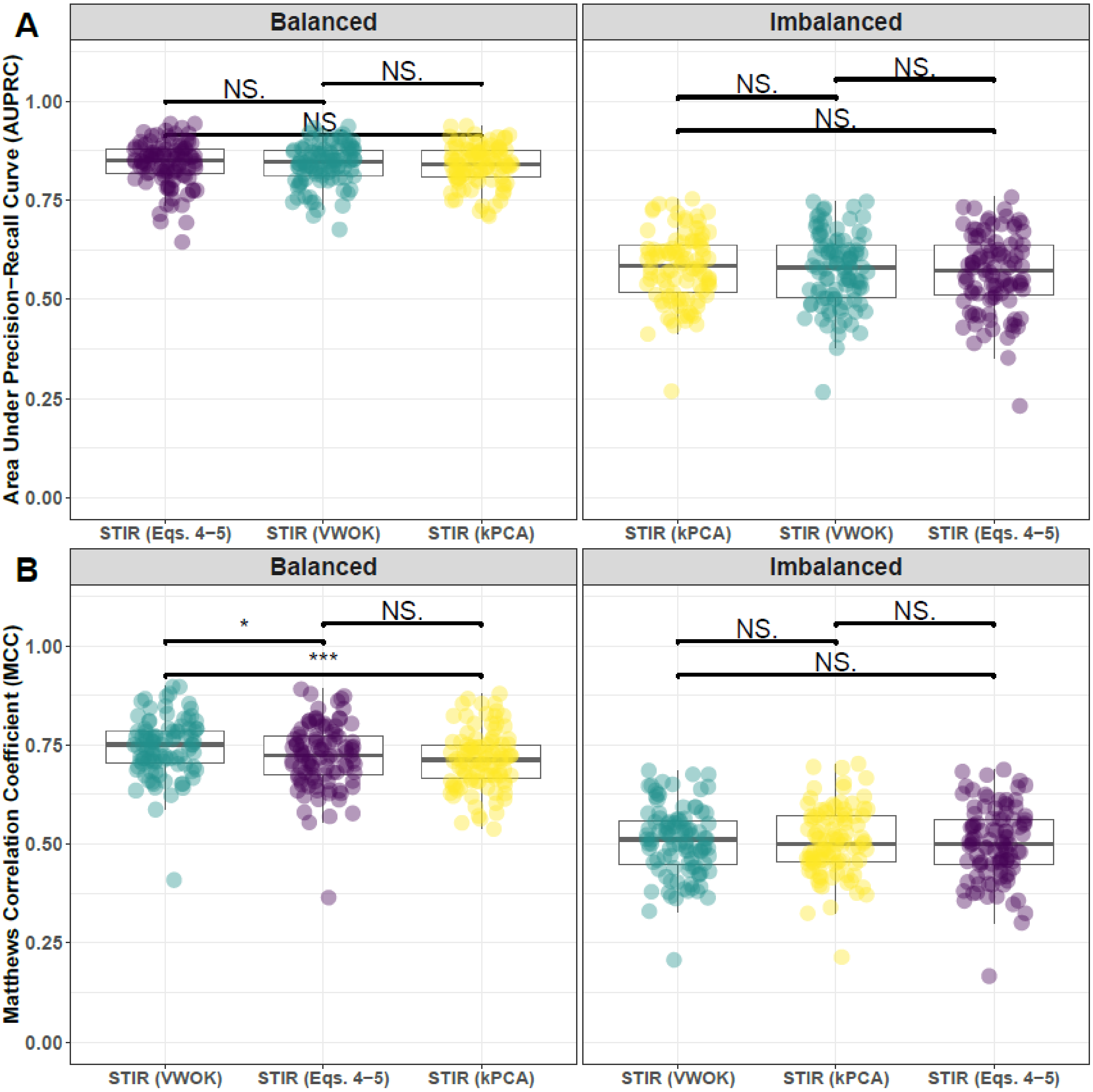


**Supplementary Fig. 10. Performance comparison for hit-miss-k (Eqs. 4 – 5), VWOK (Eqs. 6 – 7), and kPCA with STIR feature scoring.** Performance of feature selection was measured for 100 simulation replicates. Each simulated data set had $m = 100$ instances and $p = 1000$ features with 100 functional. Functional features included 25 that with main effect ($\text{bias}_{\text{main}} = 0.8$) only and the remaining 75 were involved in network interactions ($\text{bias}_{\text{int}} = 0.4$) and had no main effect. Imbalanced simulations had class ratio of 25:75 (cases:controls). (**A**) Area Under Precision-Recall Curve (AUPRC) for each method, sorted by decreasing mean AUPRC. (**B**) Matthews Correlation Coefficient (MCC) for each method, sorted by decreasing mean MCC. Comparisons were made with Mann-Whitney U test (NS. $P \geq 0.05$, $* \, P < 0.05$, and $** \, P < 0.01$).

#### 2.2 Comparing imbalance-adjusted fixed-k, regular fixed-k, and MultiSURF

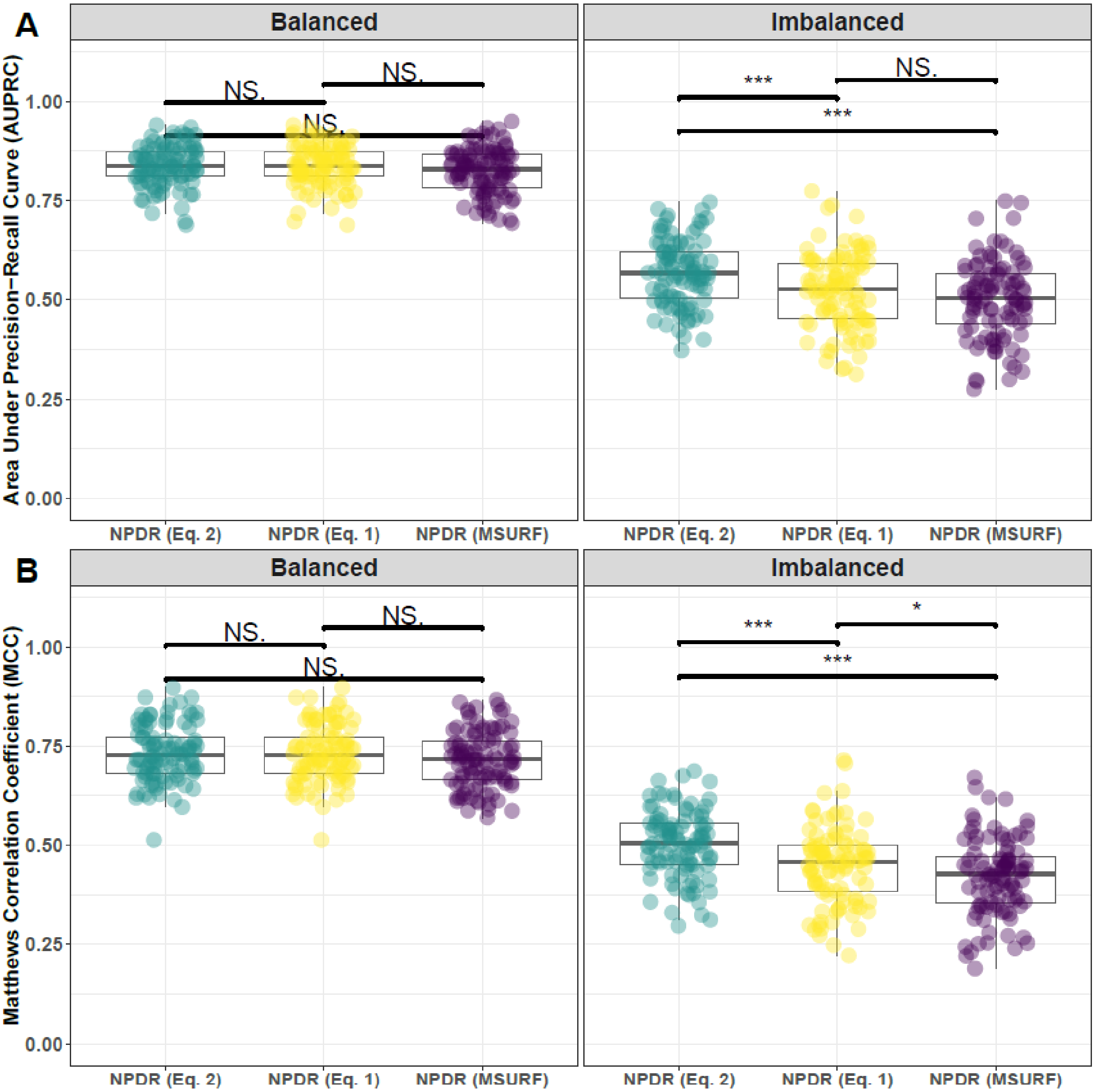


**Supplementary Fig. 11. Performance comparison minority-class-k (Eq. 2), non-adjusted fixed-k (Eq. 1), and MultiSURF with NPDR feature scoring.** Performance of feature selection was measured for 100 simulation replicates. Each simulated data set had $m = 100$ instances and $p = 1000$ features with 100 functional. Functional features included 25 that with main effect ($\text{bias}_{\text{main}} = 0.8$) only and the remaining 75 were involved in network interactions ($\text{bias}_{\text{int}} = 0.4$) and had no main effect. Imbalanced simulations had class ratio of 25:75 (cases:controls). (**A**) Area Under Precision-Recall Curve (AUPRC) for each method, sorted by decreasing mean AUPRC. (**B**) Matthews Correlation Coefficient (MCC) for each method, sorted by decreasing mean MCC. Comparisons were made with Mann-Whitney U test (NS. $P \geq 0.05$, $* \, P < 0.05$, and $*** \, P < 0.001$).

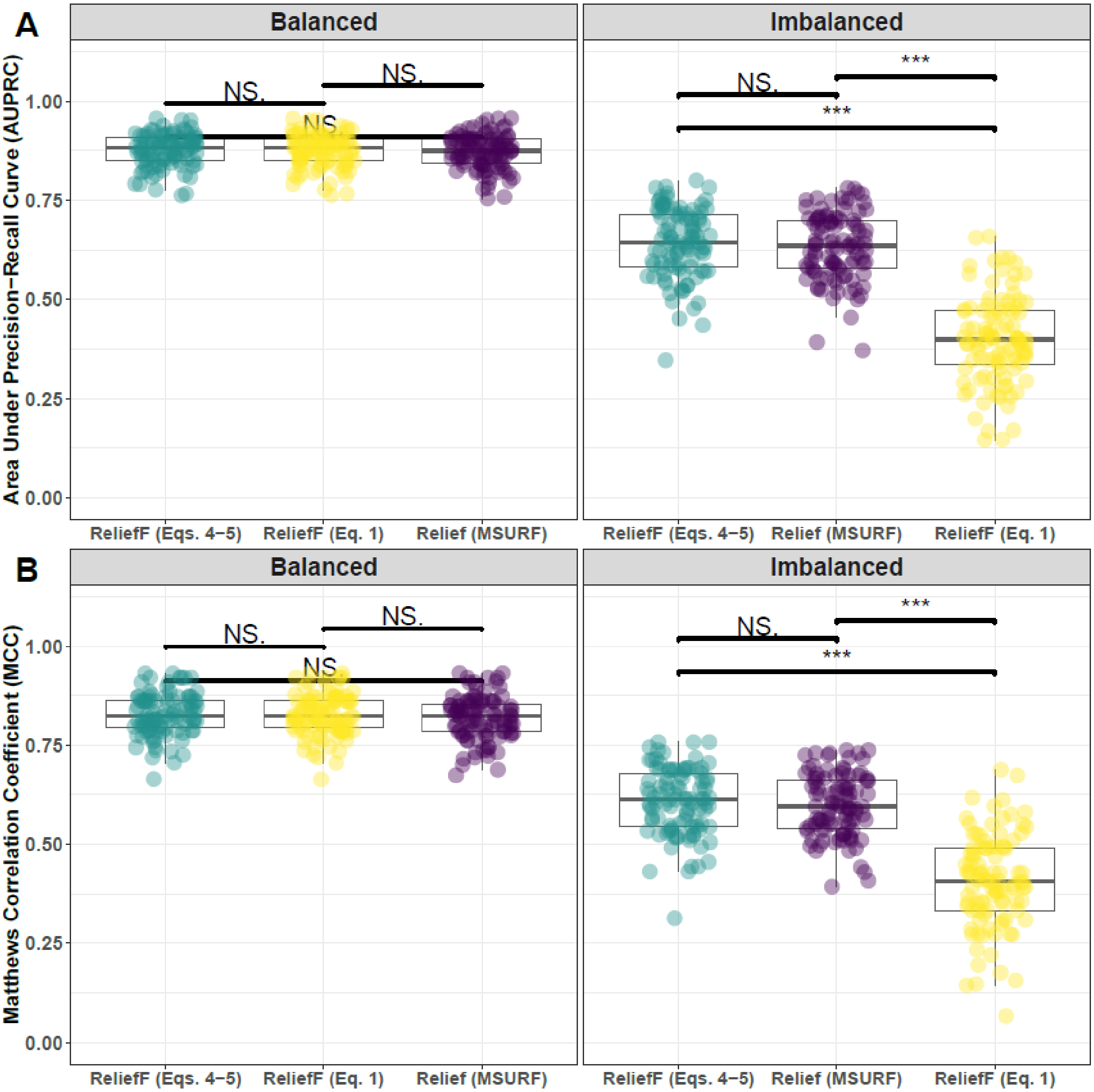


**Supplementary Fig. 12. Performance comparison of hit-miss-k (Eqs. 4 – 5), non-adjusted fixed-k (Eq. 1), and MultiSURF with ReliefF feature scoring.** Performance of feature selection was measured for 100 simulation replicates. Each simulated data set had $m = 100$ instances and $p = 1000$ features with 100 functional. Functional features included 25 that with main effect ($\text{bias}_{\text{main}} = 0.8$) only and the remaining 75 were involved in network interactions ($\text{bias}_{\text{int}} = 0.4$) and had no main effect. Imbalanced simulations had class ratio of 25:75 (cases:controls). (**A**) Area Under Precision-Recall Curve (AUPRC) for each method, sorted by decreasing mean AUPRC. (**B**) Matthews Correlation Coefficient (MCC) for each method, sorted by decreasing mean MCC. Comparisons were made with Mann-Whitney U test (NS. $P \geq 0.05$, $* \, P < 0.05$, and $*** \, P < 0.001$).

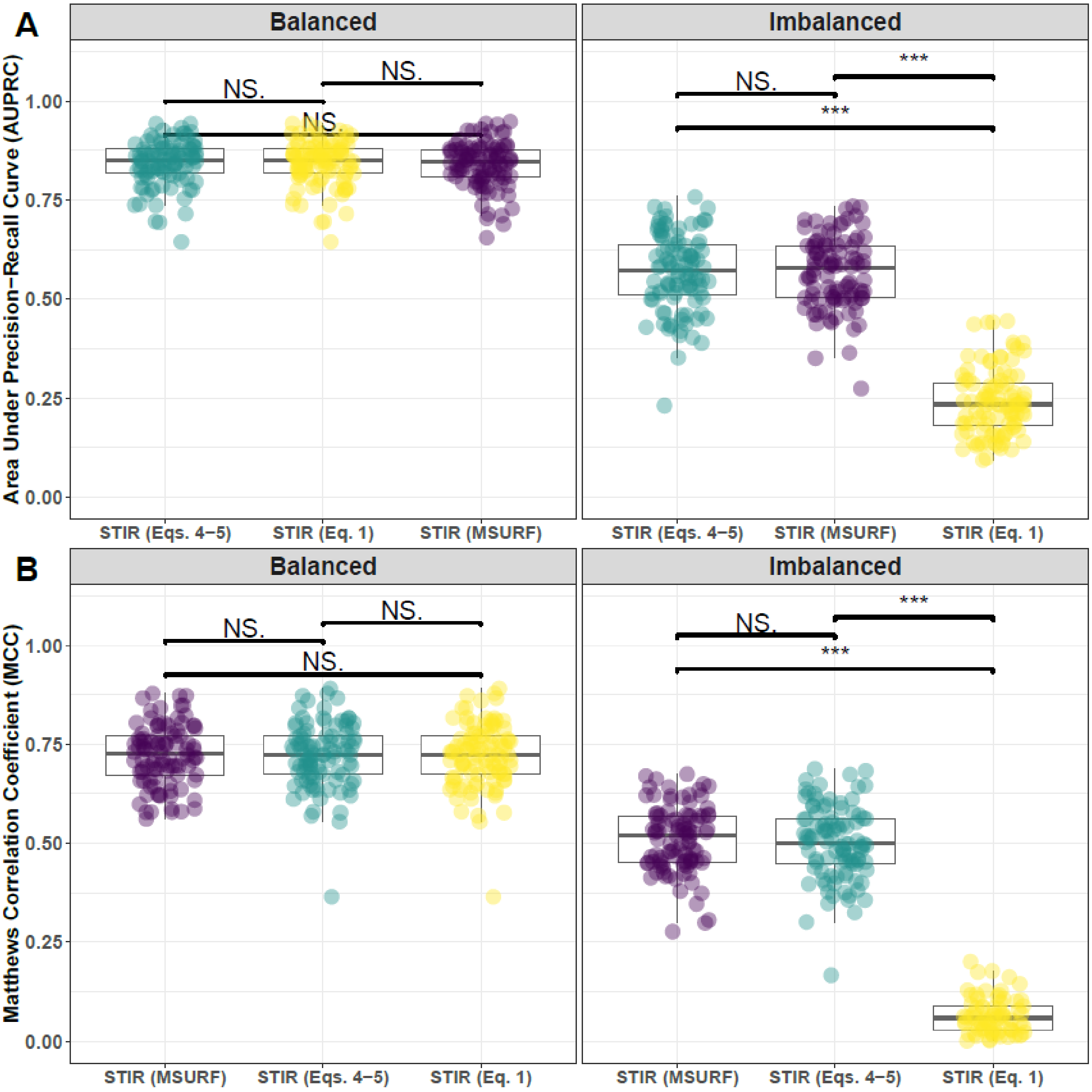


**Supplementary Fig. 13. Performance comparison of hit-miss-k (Eqs. 4 – 5), non-adjusted fixed-k (Eq. 1), and MultiSURF with STIR feature scoring.** Performance of feature selection was measured for 100 simulation replicates. Each simulated data set had $m = 100$ instances and $p = 1000$ features with 100 functional. Functional features included 25 that with main effect ($\text{bias}_{\text{main}} = 0.8$) only and the remaining 75 were involved in network interactions ($\text{bias}_{\text{int}} = 0.4$) and had no main effect. Imbalanced simulations had class ratio of 25:75 (cases:controls). (**A**) Area Under Precision-Recall Curve (AUPRC) for each method, sorted by decreasing mean AUPRC. (**B**) Matthews Correlation Coefficient (MCC) for each method, sorted by decreasing mean MCC. Comparisons were made with Mann-Whitney U test (NS. $P \geq 0.05$, $*\, P < 0.05$, and $***\, P < 0.001$).

#### 2.3 Comparing imbalance-adjusted fixed-k, random forest, and ridge regression

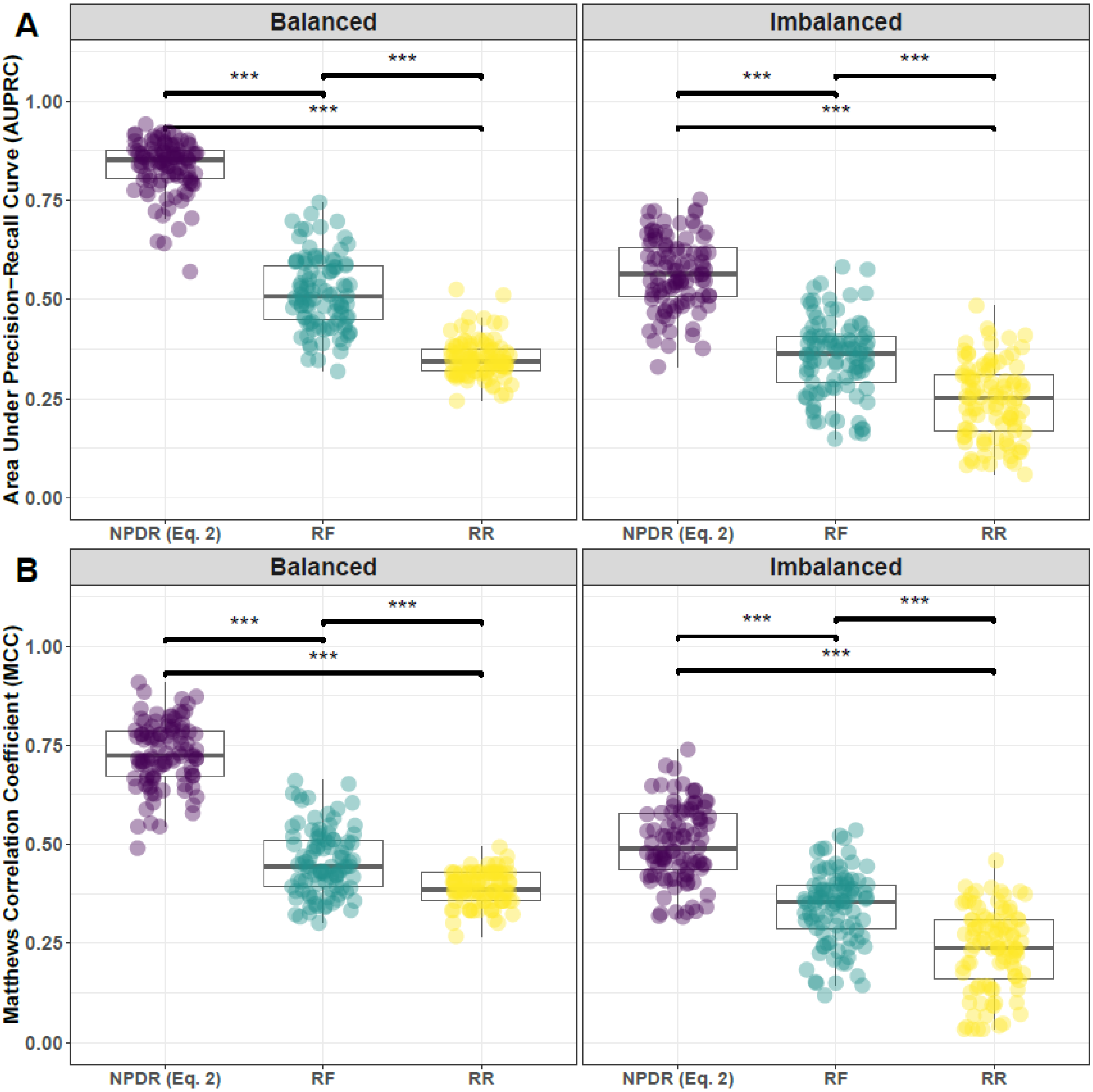


**Supplementary Fig. 14. Performance comparison of NPDR with minority-class-k (Eq. 2), Random Forest (RF), and Ridge Regression (RR).** Performance of feature selection was measured for 100 simulation replicates. Each simulated data set had $m = 100$ instances and $p = 1000$ features with 100 functional. Functional features included 25 that with main effect ($\text{bias}_{\text{main}} = 0.8$) only and the remaining 75 were involved in network interactions ($\text{bias}_{\text{int}} = 0.4$) and had no main effect. Imbalanced simulations had class ratio of 25:75 (cases:controls). (**A**) Area Under Precision-Recall Curve (AUPRC) for each method, sorted by decreasing mean AUPRC. (**B**) Matthews Correlation Coefficient (MCC) for each method, sorted by decreasing mean MCC. Comparisons were made with Mann-Whitney U test (NS. $P \geq 0.05$, $* \, P < 0.05$, and $*** \, P < 0.001$).

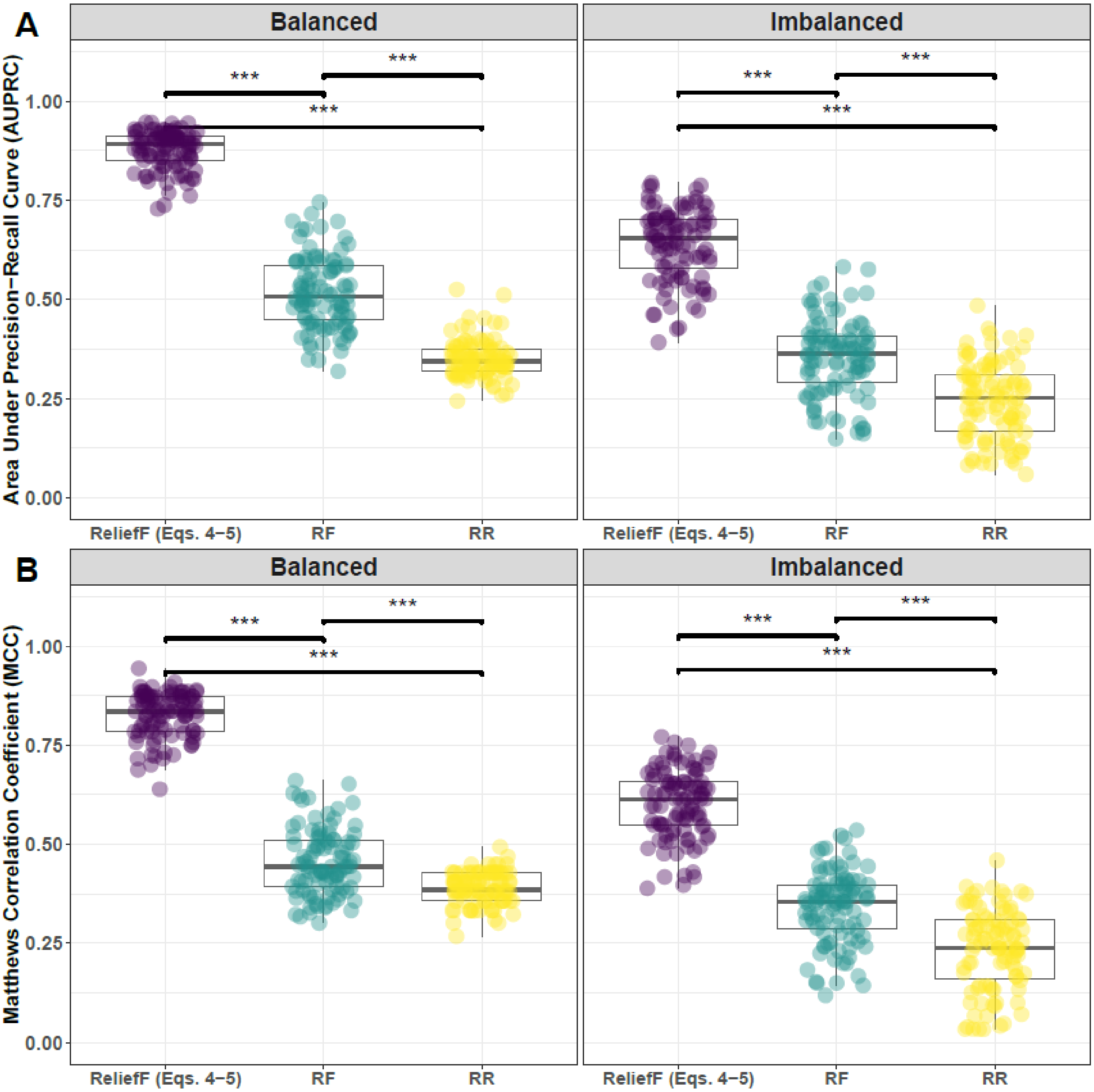


**Supplementary Fig. 15. Performance comparison of ReliefF with hit-miss-k (Eqs. 4-5), Random Forest (RF), and Ridge Regression (RR).** Performance of feature selection was measured for 100 simulation replicates. Each simulated data set had $m = 100$ instances and $p = 1000$ features with 100 functional. Functional features included 25 that with main effect ($\text{bias}_{\text{main}} = 0.8$) only and the remaining 75 were involved in network interactions ($\text{bias}_{\text{int}} = 0.4$) and had no main effect. Imbalanced simulations had class ratio of 25:75 (cases:controls). (**A**) Area Under Precision-Recall Curve (AUPRC) for each method, sorted by decreasing mean AUPRC. (**B**) Matthews Correlation Coefficient (MCC) for each method, sorted by decreasing mean MCC. Comparisons were made with Mann-Whitney U test (NS. $P \geq 0.05$, $*\, P < 0.05$, and $***\, P < 0.001$).

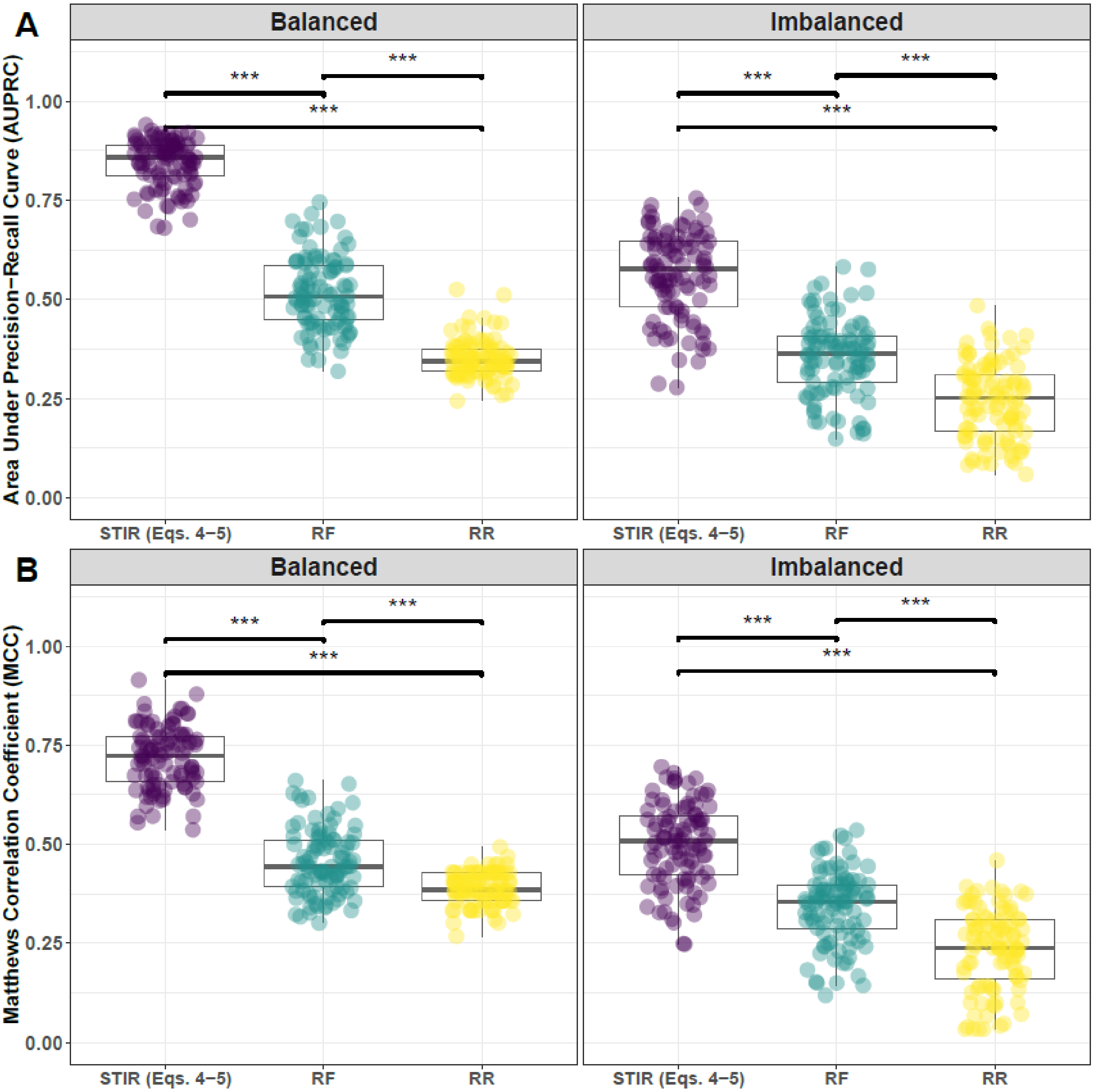


**Supplementary Fig. 16. Performance comparison of STIR with hit-miss-k (Eqs. 4-5), Random Forest (RF), and Ridge Regression (RR).** Performance of feature selection was measured for 100 simulation replicates. Each simulated data set had $m = 100$ instances and $p = 1000$ features with 100 functional. Functional features included 25 that with main effect ($\text{bias}_{\text{main}} = 0.8$) only and the remaining 75 were involved in network interactions ($\text{bias}_{\text{int}} = 0.4$) and had no main effect. Imbalanced simulations had class ratio of 25:75 (cases:controls). (**A**) Area Under Precision-Recall Curve (AUPRC) for each method, sorted by decreasing mean AUPRC. (**B**) Matthews Correlation Coefficient (MCC) for each method, sorted by decreasing mean MCC. Comparisons were made with Mann-Whitney U test (NS. $P \geq 0.05$, $*\, P < 0.05$, and $***\, P < 0.001$).

### 3 Feature selection performance comparisons: 25% interaction effect/75% main effect

#### 3.1 Comparing imbalance-adjusted fixed-k, VWOK, and kPCA

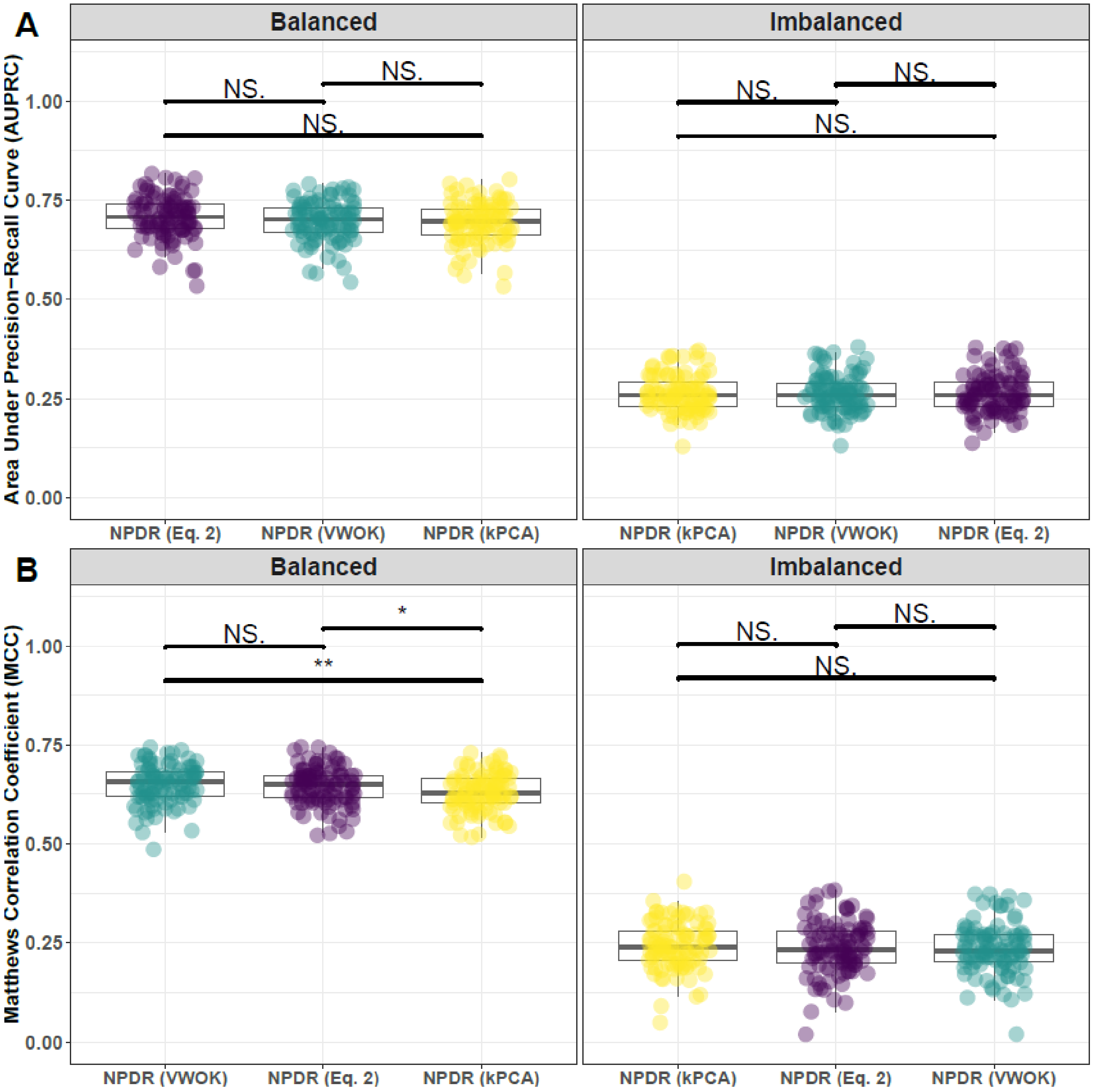


**Supplementary Fig. 17. Performance comparison for minority-class-k (Eq. 2), VWOK (Eq. 3), and kPCA with NPDR feature scoring.** Performance of feature selection was measured for 100 simulation replicates. Each simulated data set had $m = 100$ instances and $p = 1000$ features with 100 functional. Functional features included 75 that with main effect ($\text{bias}_{\text{main}} = 0.8$) only and the remaining 25 were involved in network interactions ($\text{bias}_{\text{int}} = 0.4$) and had no main effect. Imbalanced simulations had class ratio of 25:75 (cases:controls). (**A**) Area Under Precision-Recall Curve (AUPRC) for each method, sorted by decreasing mean AUPRC. (**B**) Matthews Correlation Coefficient (MCC) for each method, sorted by decreasing mean MCC. Comparisons were made with Mann-Whitney U test (NS. $P \geq 0.05$, $*\,P < 0.05$, and $**\,P < 0.01$).

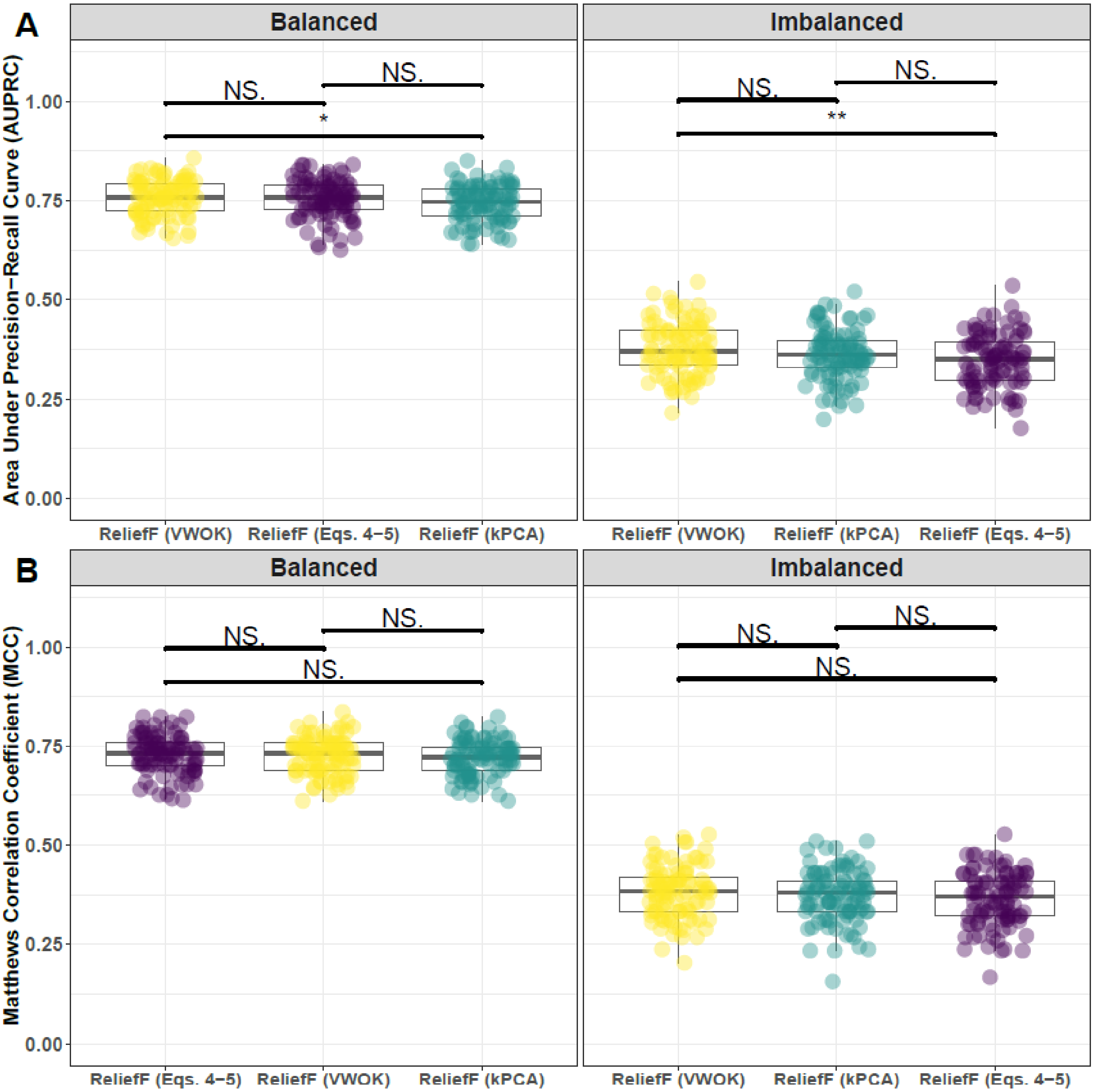


**Supplementary Fig. 18. Performance comparison for hit-miss-k (Eqs. 4 – 5), VWOK (Eqs. 6 – 7), and kPCA with ReliefF feature scoring.** Performance of feature selection was measured for 100 simulation replicates. Each simulated data set had $m = 100$ instances and $p = 1000$ features with 100 functional. Functional features included 75 that with main effect ($\text{bias}_{\text{main}} = 0.8$) only and the remaining 25 were involved in network interactions ($\text{bias}_{\text{int}} = 0.4$) and had no main effect. Imbalanced simulations had class ratio of 25:75 (cases:controls). (**A**) Area Under Precision-Recall Curve (AUPRC) for each method, sorted by decreasing mean AUPRC. (**B**) Matthews Correlation Coefficient (MCC) for each method, sorted by decreasing mean MCC. Comparisons were made with Mann-Whitney U test (NS. $P \geq 0.05$, $* \, P < 0.05$, and $** \, P < 0.01$).

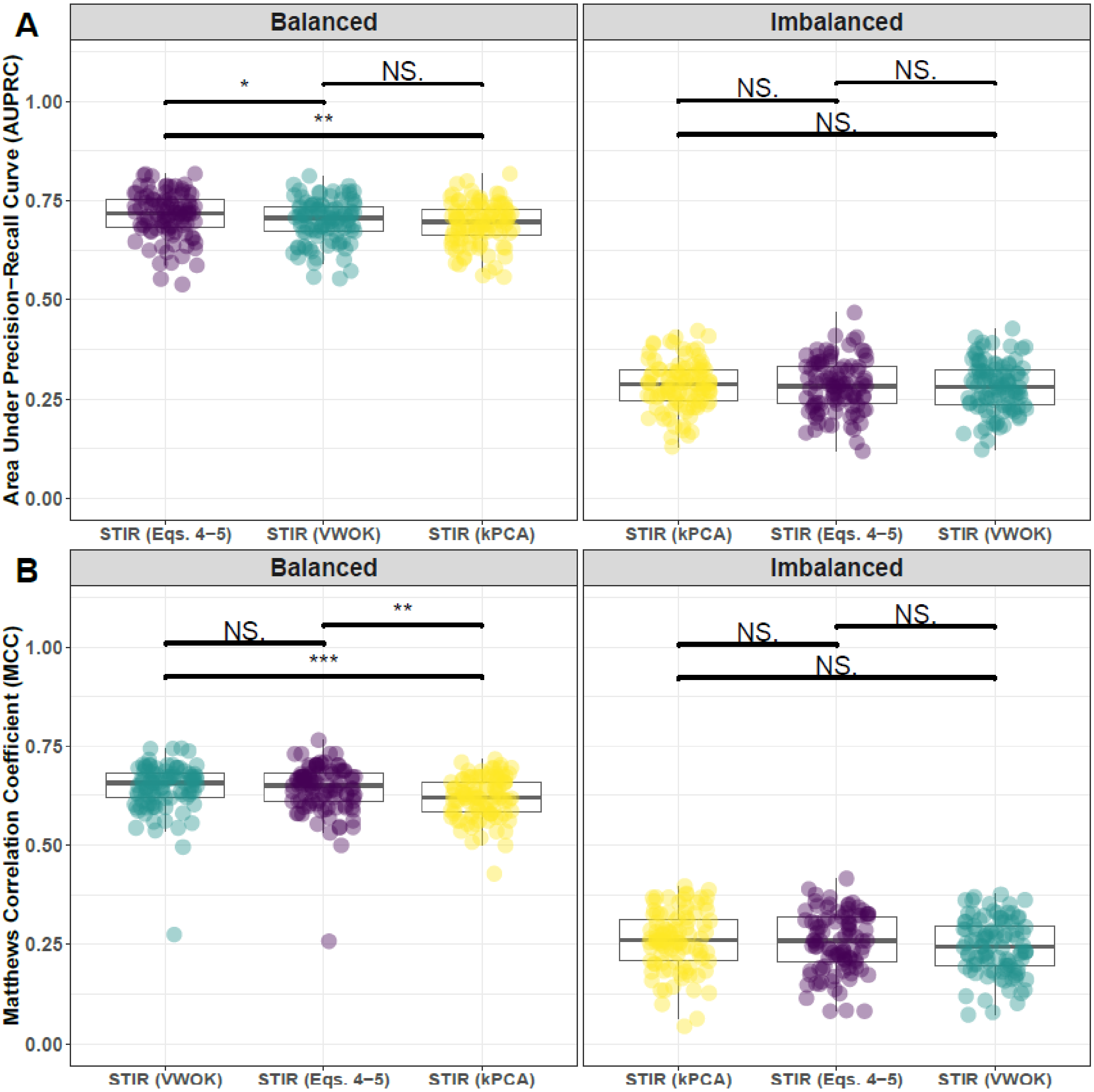


**Supplementary Fig. 19. Performance comparison for hit-miss-k (Eqs. 4 – 5), VWOK (Eqs. 6 – 7), and kPCA with STIR feature scoring.** Performance of feature selection was measured for 100 simulation replicates. Each simulated data set had $m = 100$ instances and $p = 1000$ features with 100 functional. Functional features included 75 that with main effect ($\text{bias}_{\text{main}} = 0.8$) only and the remaining 25 were involved in network interactions ($\text{bias}_{\text{int}} = 0.4$) and had no main effect. Imbalanced simulations had class ratio of 25:75 (cases:controls). (**A**) Area Under Precision-Recall Curve (AUPRC) for each method, sorted by decreasing mean AUPRC. (**B**) Matthews Correlation Coefficient (MCC) for each method, sorted by decreasing mean MCC. Comparisons were made with Mann-Whitney U test (NS. $P \geq 0.05$, $* \, P < 0.05$, and $** \, P < 0.01$).

#### 3.2 Comparing imbalance-adjusted fixed-k, regular fixed-k, and MultiSURF

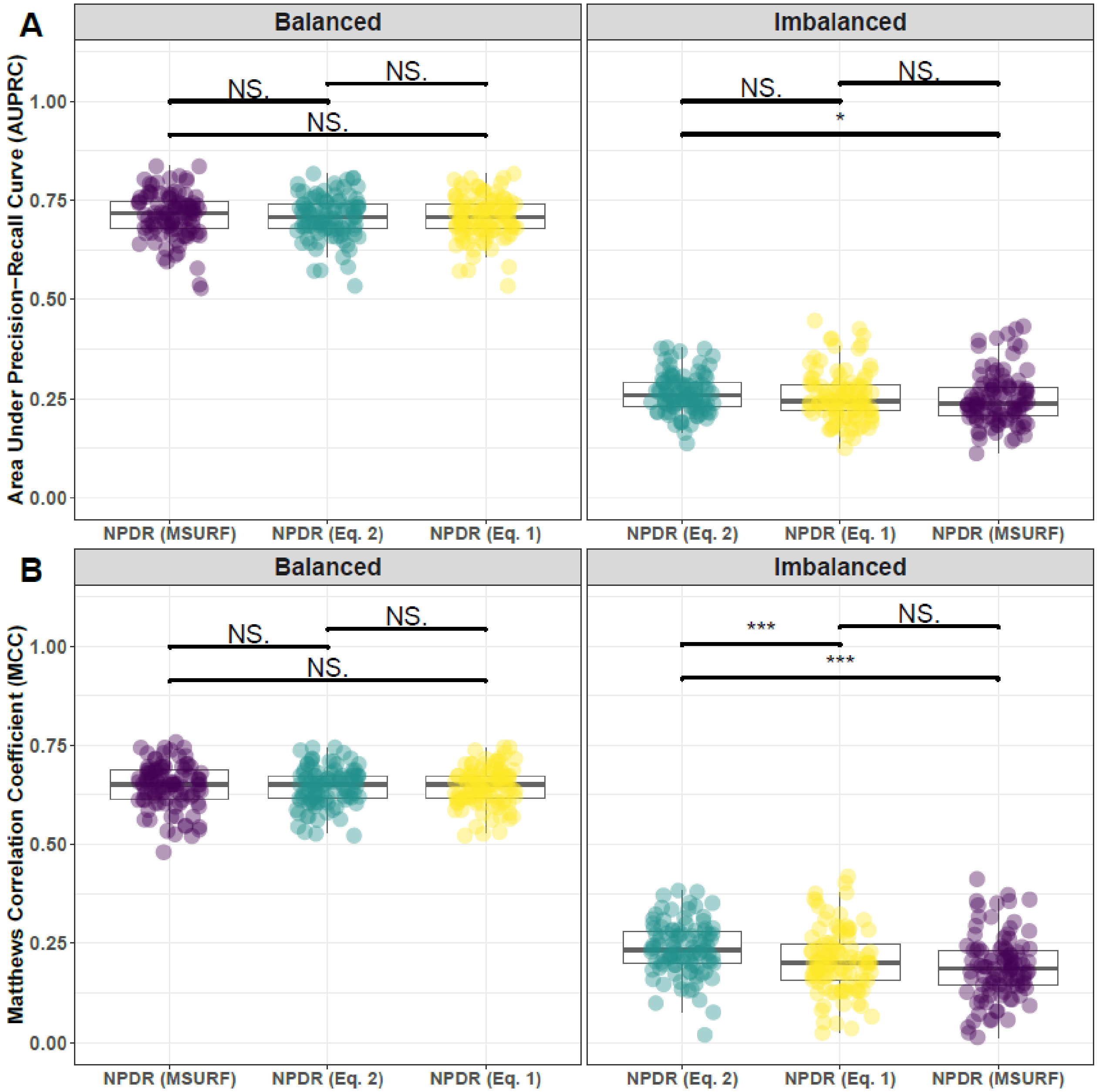


**Supplementary Fig. 20. Performance comparison of minority-class-k (Eq. 2), non-adjusted fixed-k (Eq. 1), and MultiSURF with NPDR feature scoring.** Performance of feature selection was measured for 100 simulation replicates. Each simulated data set had $m = 100$ instances and $p = 1000$ features with 100 functional. Functional features included 75 that with main effect ($\text{bias}_{\text{main}} = 0.8$) only and the remaining 25 were involved in network interactions ($\text{bias}_{\text{int}} = 0.4$) and had no main effect. Imbalanced simulations had class ratio of 25:75 (cases:controls). (**A**) Area Under Precision-Recall Curve (AUPRC) for each method, sorted by decreasing mean AUPRC. (**B**) Matthews Correlation Coefficient (MCC) for each method, sorted by decreasing mean MCC. Comparisons were made with Mann-Whitney U test (NS. $P \geq 0.05$, $* P < 0.05$, and $*** P < 0.001$).

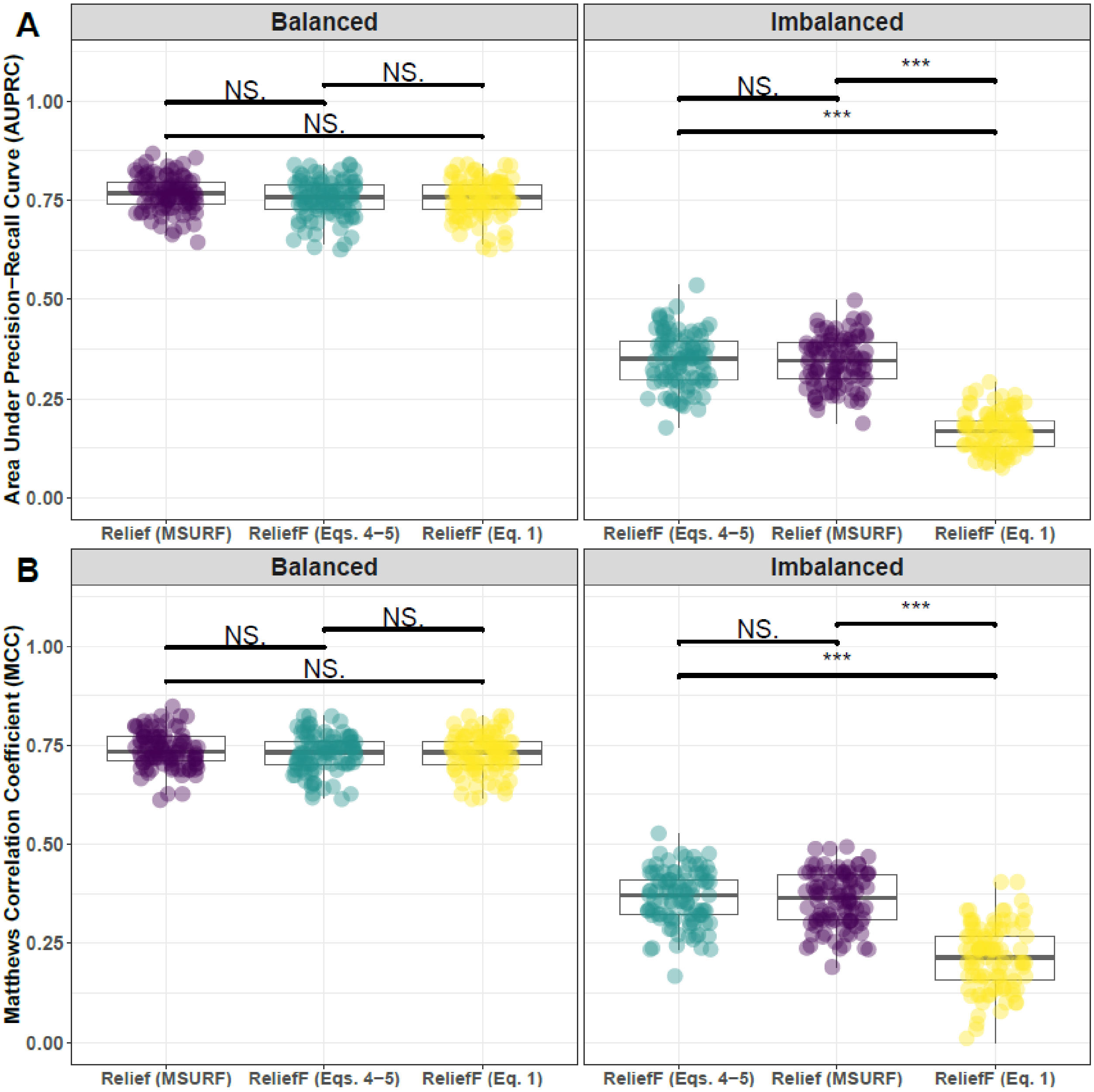


**Supplementary Fig. 21. Performance comparison of hit-miss-k (Eqs. 4 – 5), non-adjusted fixed-k (Eq. 1), and MultiSURF with ReliefF feature scoring.** Performance of feature selection was measured for 100 simulation replicates. Each simulated data set had $m = 100$ instances and $p = 1000$ features with 100 functional. Functional features included 75 that with main effect ($\text{bias}_{\text{main}} = 0.8$) only and the remaining 25 were involved in network interactions ($\text{bias}_{\text{int}} = 0.4$) and had no main effect. Imbalanced simulations had class ratio of 25:75 (cases:controls). (**A**) Area Under Precision-Recall Curve (AUPRC) for each method, sorted by decreasing mean AUPRC. (**B**) Matthews Correlation Coefficient (MCC) for each method, sorted by decreasing mean MCC. Comparisons were made with Mann-Whitney U test (NS. $P \geq 0.05$, $* P < 0.05$, and $*** P < 0.001$).

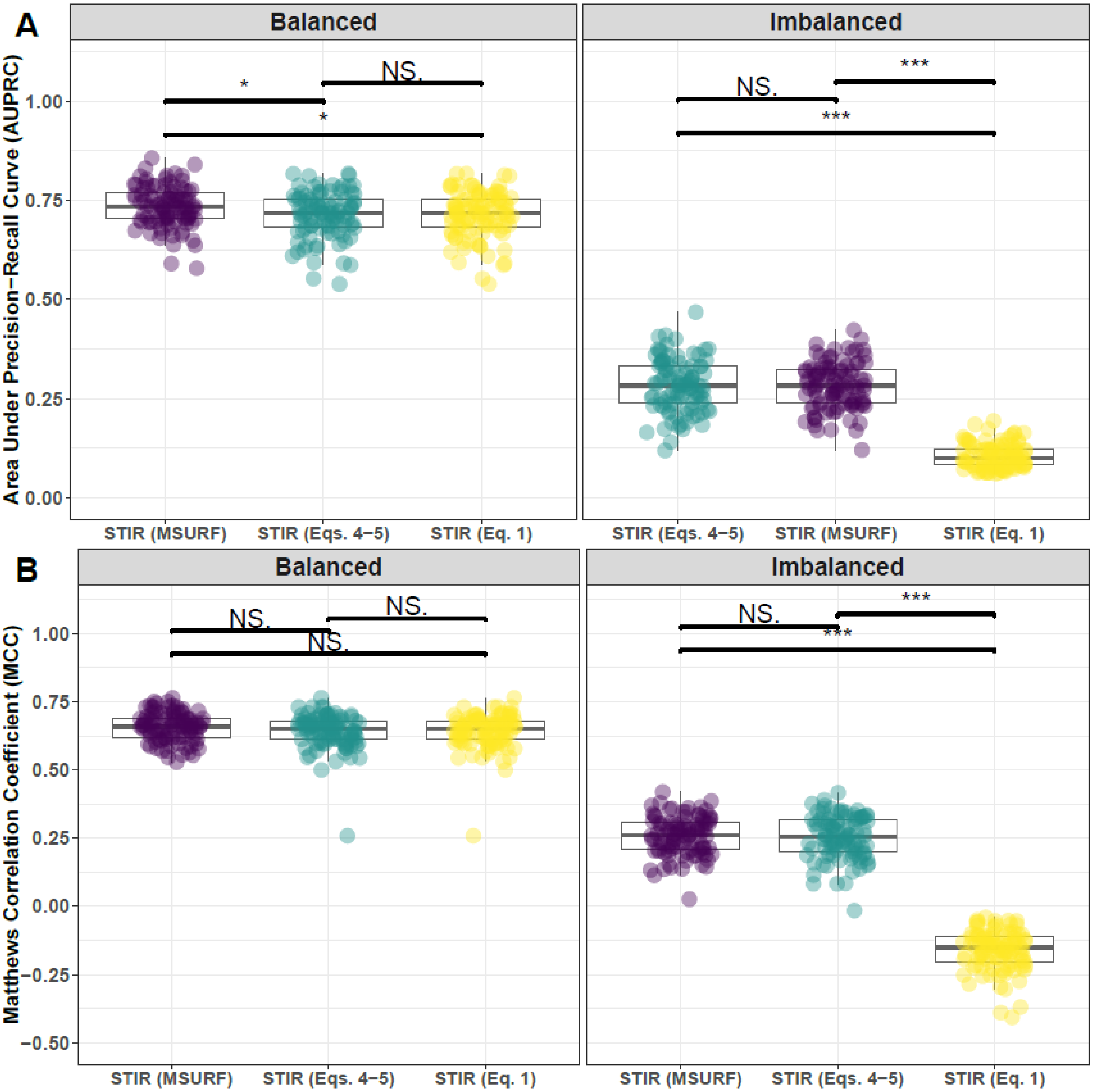


**Supplementary Fig. 22. Performance comparison of hit-miss-k (Eqs. 4 – 5), non-adjusted fixed-k (Eq. 1), and MultiSURF with STIR feature scoring.** Performance of feature selection was measured for 100 simulation replicates. Each simulated data set had $m = 100$ instances and $p = 1000$ features with 100 functional. Functional features included 75 that with main effect ($\text{bias}_{\text{main}} = 0.8$) only and the remaining 25 were involved in network interactions ($\text{bias}_{\text{int}} = 0.4$) and had no main effect. Imbalanced simulations had class ratio of 25:75 (cases:controls). (**A**) Area Under Precision-Recall Curve (AUPRC) for each method, sorted by decreasing mean AUPRC. (**B**) Matthews Correlation Coefficient (MCC) for each method, sorted by decreasing mean MCC. Comparisons were made with Mann-Whitney U test (NS. $P \geq 0.05$, $* P < 0.05$, and $*** P < 0.001$).

##### 3.3 Comparing imbalance-adjusted fixed-k, random forest, and ridge regression

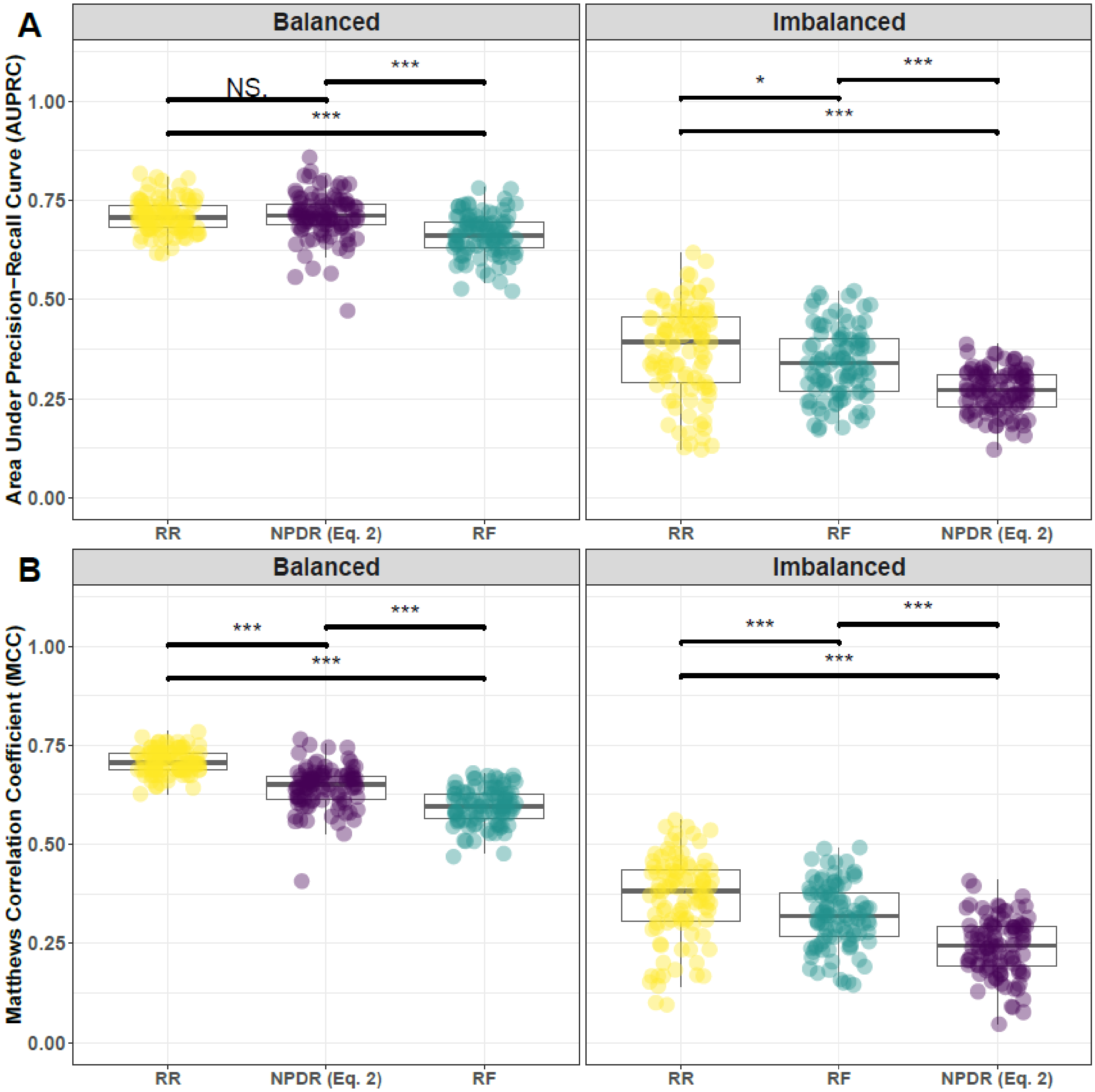


**Supplementary Fig. 23. Performance comparison of NPDR with minority-class-k (Eq. 2), Random Forest (RF), and Ridge Regression (RR).** Performance of feature selection was measured for 100 simulation replicates. Each simulated data set had $m = 100$ instances and $p = 1000$ features with 100 functional. Functional features included 75 that with main effect ($\text{bias}_{\text{main}} = 0.8$) only and the remaining 25 were involved in network interactions ($\text{bias}_{\text{int}} = 0.4$) and had no main effect. Imbalanced simulations had class ratio of 25:75 (cases:controls). (**A**) Area Under Precision-Recall Curve (AUPRC) for each method, sorted by decreasing mean AUPRC. (**B**) Matthews Correlation Coefficient (MCC) for each method, sorted by decreasing mean MCC. Comparisons were made with Mann-Whitney U test (NS. $P \geq 0.05$, $* \, P < 0.05$, and $*** \, P < 0.001$).

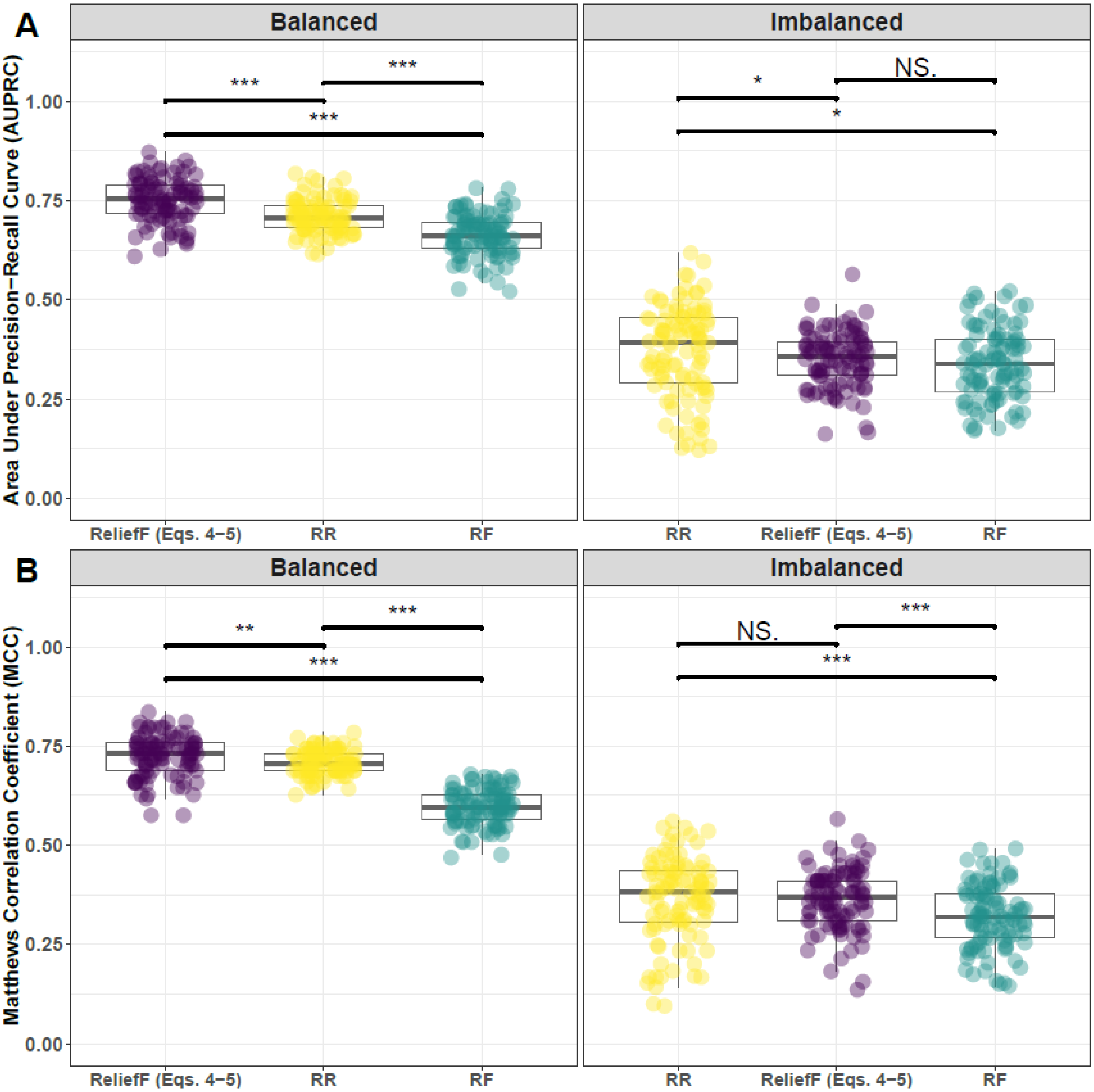


**Supplementary Fig. 24. Performance comparison of ReliefF with hit-miss-k (Eqs. 4 – 5), Random Forest (RF), and Ridge Regression (RR).** Performance of feature selection was measured for 100 simulation replicates. Each simulated data set had $m = 100$ instances and $p = 1000$ features with 100 functional. Functional features included 75 that with main effect ($\text{bias}_{\text{main}} = 0.8$) only and the remaining 25 were involved in network interactions ($\text{bias}_{\text{int}} = 0.4$) and had no main effect. Imbalanced simulations had class ratio of 25:75 (cases:controls). (**A**) Area Under Precision-Recall Curve (AUPRC) for each method, sorted by decreasing mean AUPRC. (**B**) Matthews Correlation Coefficient (MCC) for each method, sorted by decreasing mean MCC. Comparisons were made with Mann-Whitney U test (NS. $P \geq 0.05$, $* \, P < 0.05$, and $*** \, P < 0.001$).

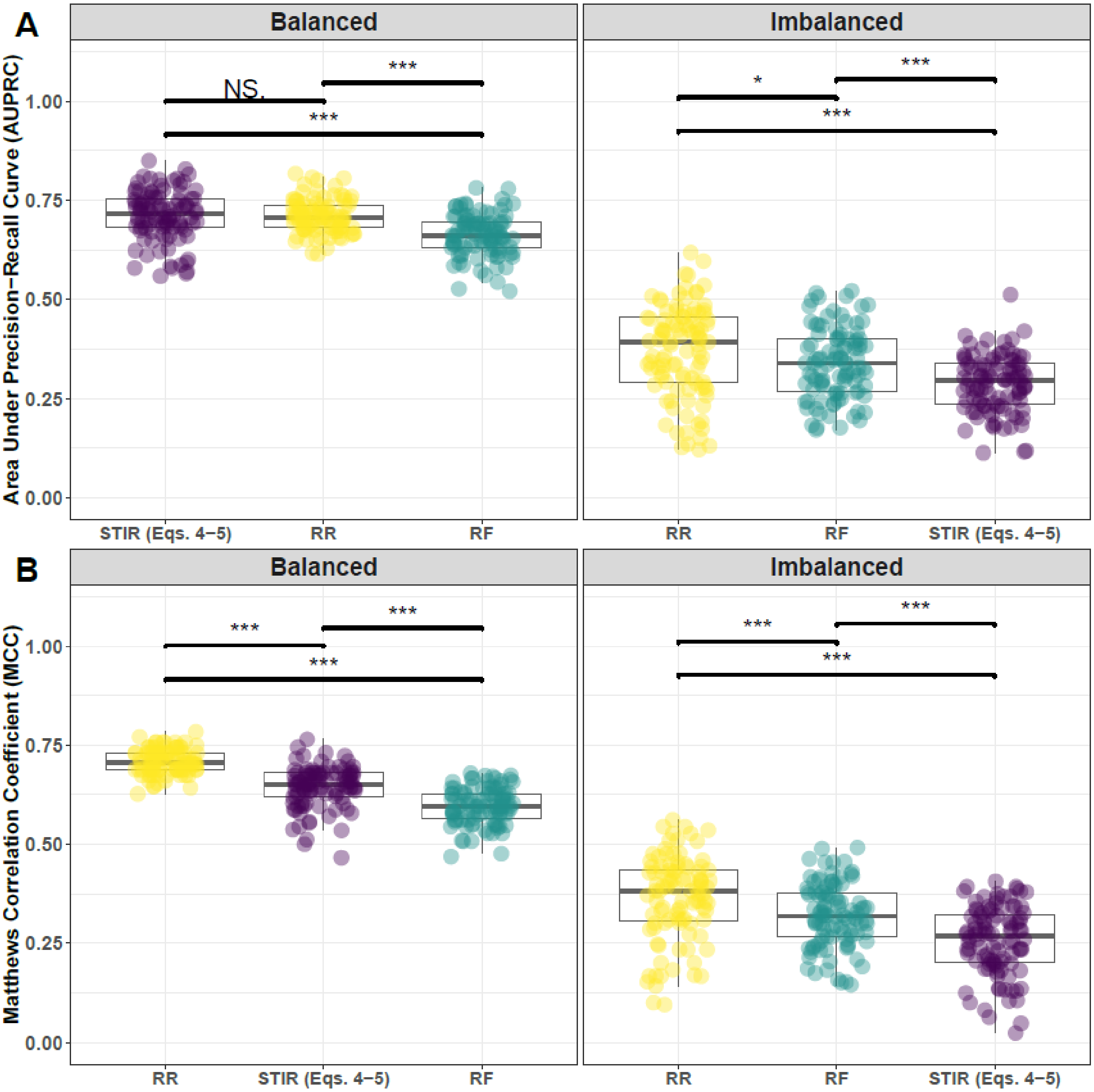


**Supplementary Fig. 25. Performance comparison of STIR with hit-miss-k (Eqs. 4 – 5), Random Forest (RF), and Ridge Regression (RR).** Performance of feature selection was measured for 100 simulation replicates. Each simulated data set had $m = 100$ instances and $p = 1000$ features with 100 functional. Functional features included 75 that with main effect ($\text{bias}_{\text{main}} = 0.8$) only and the remaining 25 were involved in network interactions ($\text{bias}_{\text{int}} = 0.4$) and had no main effect. Imbalanced simulations had class ratio of 25:75 (cases:controls). (**A**) Area Under Precision-Recall Curve (AUPRC) for each method, sorted by decreasing mean AUPRC. (**B**) Matthews Correlation Coefficient (MCC) for each method, sorted by decreasing mean MCC. Comparisons were made with Mann-Whitney U test (NS. $P \geq 0.05$, $* \, P < 0.05$, and $*** \, P < 0.001$).

### 4 Feature selection performance comparisons within consensus-features nested cross-validation (cnCV): equal main effect and interaction effect

#### 4.1 Comparing imbalance-adjusted fixed-k, VWOK, and kPCA

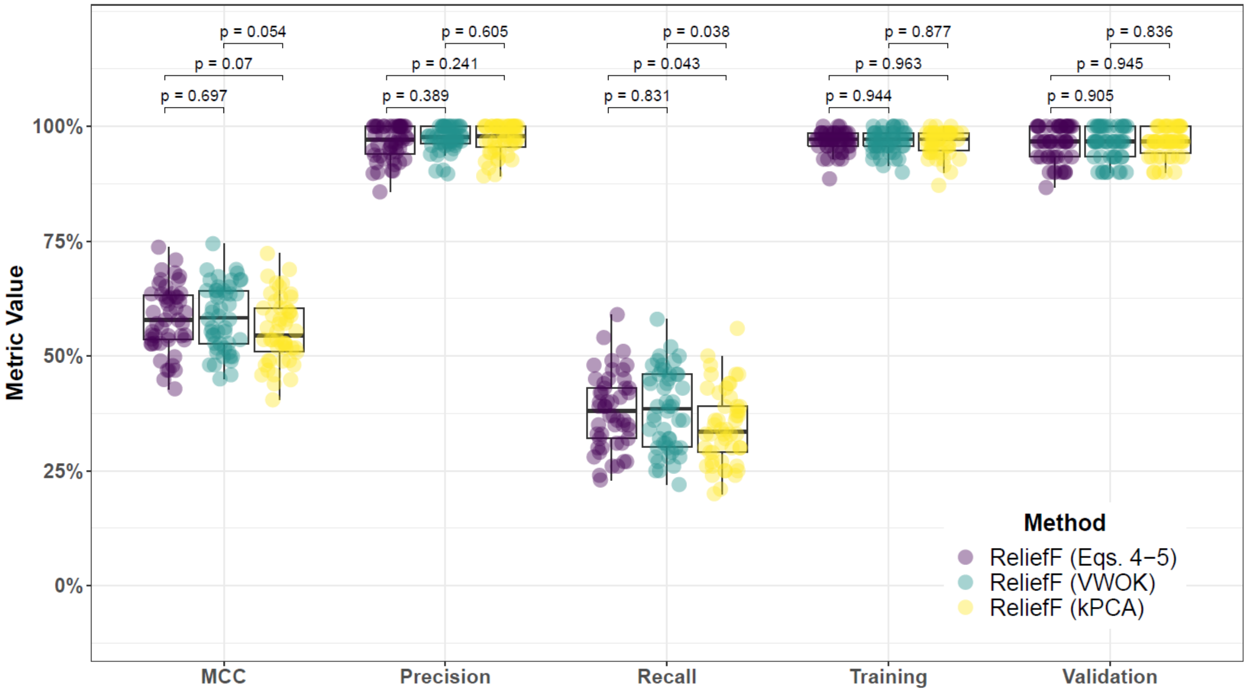


**Supplementary Fig. 26. Performance comparison for hit-miss-k (Eqs. 4 – 5), VWOK (Eqs. 6 – 7), and kPCA with ReliefF feature scoring and consensus-features nested Cross-Validation (cnCV) on balanced data.** Performance of feature selection was measured for 50 simulation replicates. Each simulated data set had $m = 100$ instances, $p = 1000$ features with 100 functional, and balanced class groups with 50 'case' and 50 'control'. Functional features included 50 with main effect ($\text{bias}_{\text{main}} = 0.8$) only and the remaining 50 were involved in network interactions ($\text{bias}_{\text{int}} = 0.4$) and had no main effect. Data was first split into training and test (70% train/30% test) sets and 5 folds were used for inner and outer training loops. The top 30% of features, ranked in decreasing order of importance, were selected within each inner training fold. Matthew's Correlation Coefficient (MCC), precision, and recall were calculated based on detection of functional features. Training and validation represent the balanced classification accuracy of the random forest model that was fit using consensus features from cnCV on full training data (70% of full dataset) and independent test data (30% of full dataset), respectively. Comparisons were made with Mann-Whitney U test.

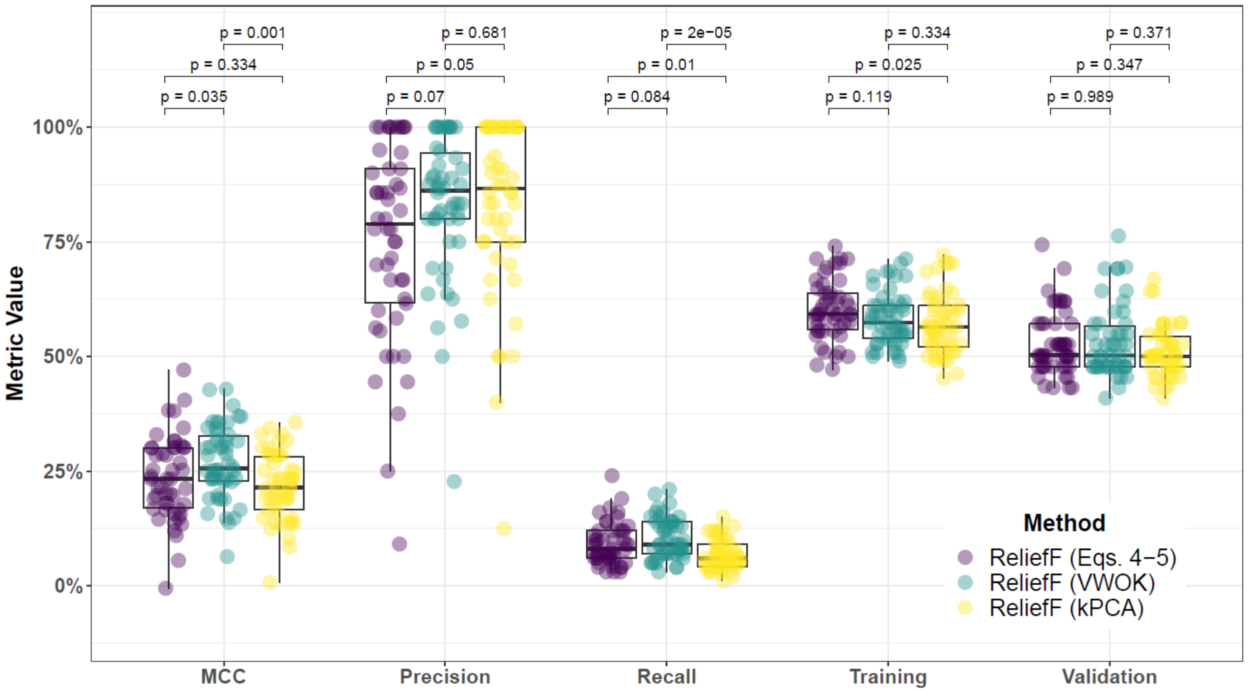


**Supplementary Fig. 27. Performance comparison for hit-miss-k (Eqs. 4 – 5), VWOK (Eqs. 6 – 7), and kPCA with ReliefF feature scoring and consensus-features nested Cross-Validation (cnCV) on imbalanced data.** Performance of feature selection was measured for 50 simulation replicates. Each simulated data set had $m = 100$ instances, $p = 1000$ features with 100 functional, and imbalanced class groups with 25 'case' and 75 'control'. Functional features included 50 with main effect ($\text{bias}_{\text{main}} = 0.8$) only and the remaining 50 were involved in network interactions ($\text{bias}_{\text{int}} = 0.4$) and had no main effect. Data was first split into training and test (70% train/30% test) sets and 5 folds were used for inner and outer training loops. The top 30% of features, ranked in decreasing order of importance, were selected within each inner training fold. Matthew's Correlation Coefficient (MCC), precision, and recall were calculated based on detection of functional features. Training and validation represent the balanced classification accuracy of the random forest model that was fit using consensus features from cnCV on full training data (70% of full dataset) and independent test data (30% of full dataset), respectively. Comparisons were made with Mann-Whitney U test.

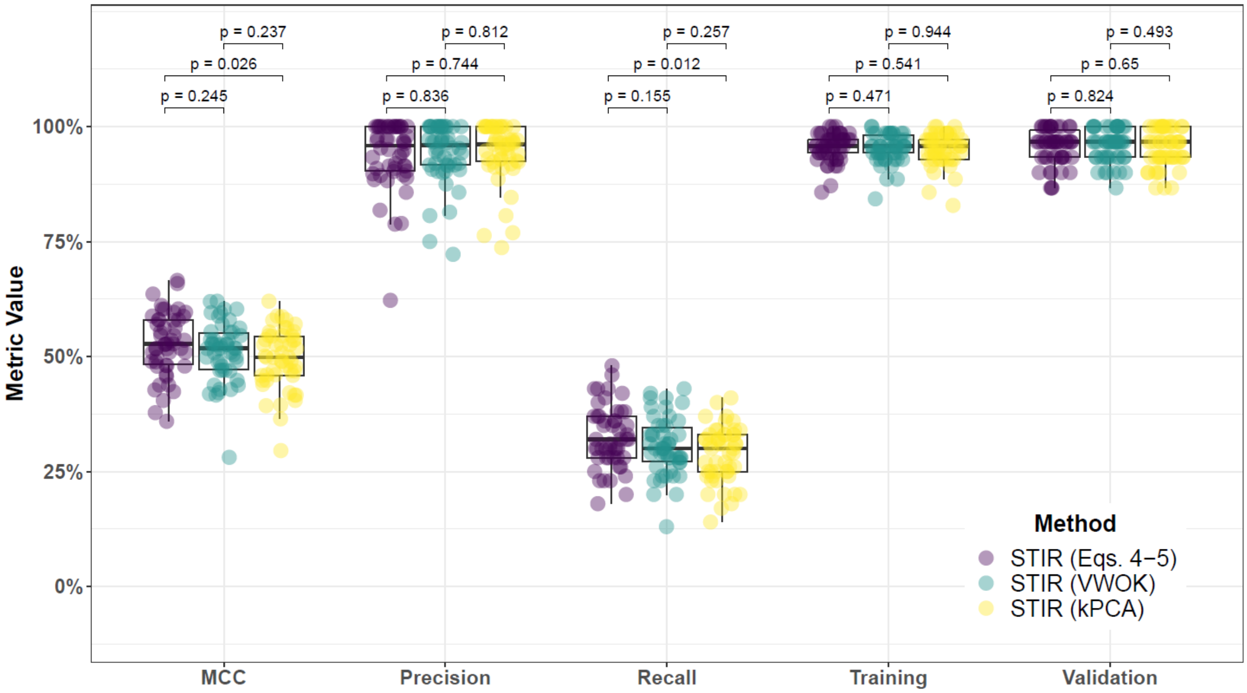


**Supplementary Fig. 28. Performance comparison for hit-miss-k (Eqs. 4 – 5), VWOK (Eqs. 6 – 7), and kPCA with STIR feature scoring and consensus-features nested Cross-Validation (cnCV) on balanced data.** Performance of feature selection was measured for 50 simulation replicates. Each simulated data set had $m = 100$ instances, $p = 1000$ features with 100 functional, and balanced class groups with 50 'case' and 50 'control'. Functional features included 50 with main effect ($\text{bias}_{\text{main}} = 0.8$) only and the remaining 50 were involved in network interactions ($\text{bias}_{\text{int}} = 0.4$) and had no main effect. Data was first split into training and test (70% train/30% test) sets and 5 folds were used for inner and outer training loops. The top 30% of features, ranked in decreasing order of importance, were selected within each inner training fold. Matthew's Correlation Coefficient (MCC), precision, and recall were calculated based on detection of functional features. Training and validation represent the balanced classification accuracy of the random forest model that was fit using consensus features from cnCV on full training data (70% of full dataset) and independent test data (30% of full dataset), respectively. Comparisons were made with Mann-Whitney U test.

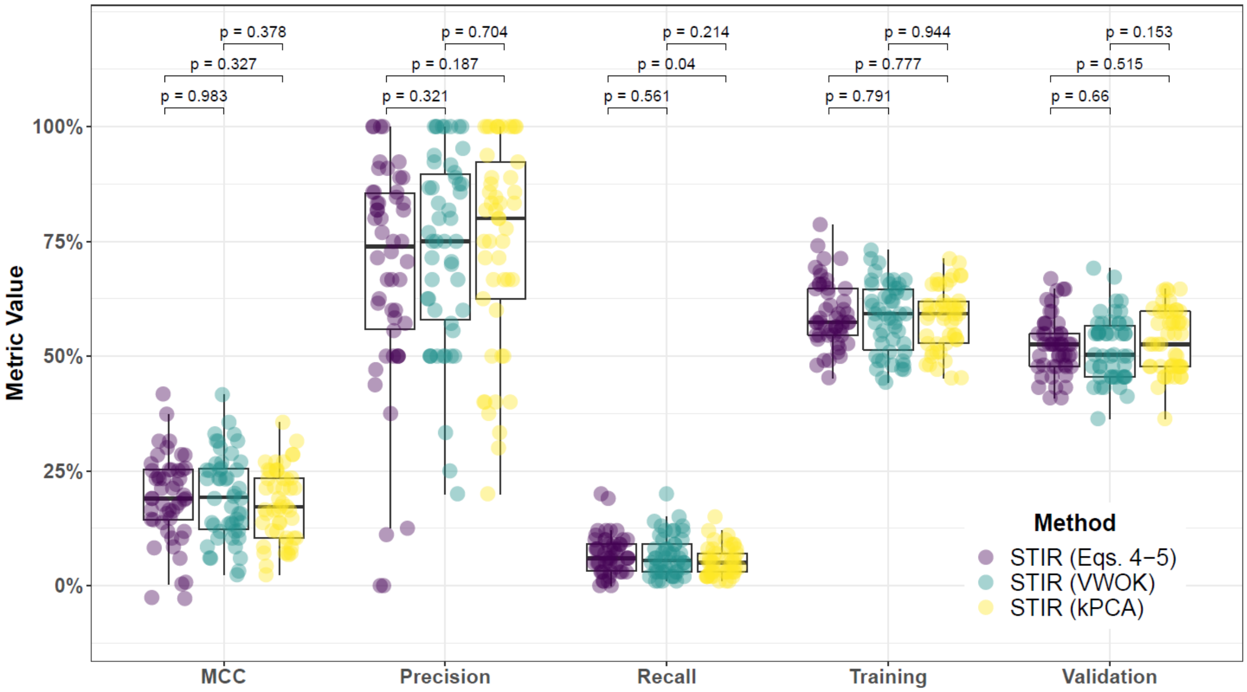


**Supplementary Fig. 29. Performance comparison for hit-miss-k (Eqs. 4 – 5), VWOK (Eqs. 6 – 7), and kPCA with STIR feature scoring and consensus-features nested Cross-Validation (cnCV) on imbalanced data.** Performance of feature selection was measured for 50 simulation replicates. Each simulated data set had $m = 100$ instances, $p = 1000$ features with 100 functional, and imbalanced class groups with 25 'case' and 75 'control'. Functional features included 50 with main effect ($\text{bias}_{\text{main}} = 0.8$) only and the remaining 50 were involved in network interactions ($\text{bias}_{\text{int}} = 0.4$) and had no main effect. Data was first split into training and test (70% train/30% test) sets and 5 folds were used for inner and outer training loops. The top 30% of features, ranked in decreasing order of importance, were selected within each inner training fold. Matthew's Correlation Coefficient (MCC), precision, and recall were calculated based on detection of functional features. Training and validation represent the balanced classification accuracy of the random forest model that was fit using consensus features from cnCV on full training data (70% of full dataset) and independent test data (30% of full dataset), respectively. Comparisons were made with Mann-Whitney U test.

#### 4.2 Comparing imbalance-adjusted fixed-k, regular fixed-k, and MultiSURF

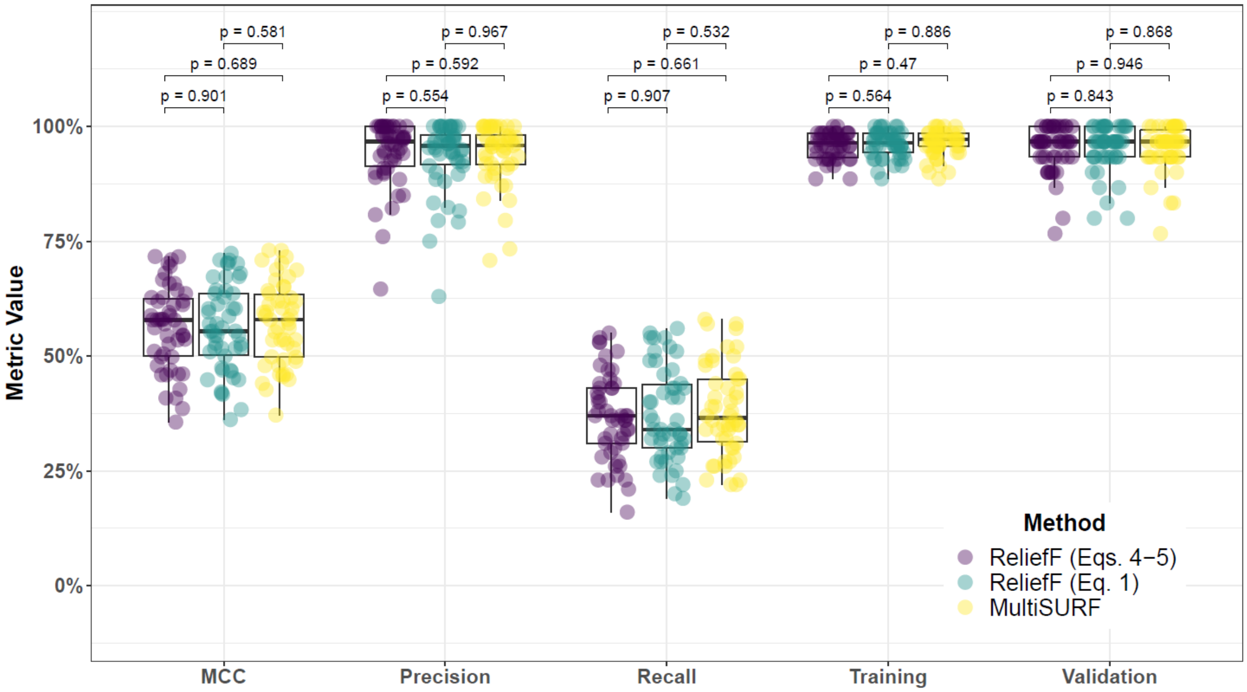


**Supplementary Fig. 30. Performance comparison for hit-miss-k (Eqs. 4 – 5), non-adjusted fixed-k (Eq. 1), and MultiSURF with ReliefF feature scoring and consensus-features nested Cross-Validation (cnCV) on balanced data.** Performance of feature selection was measured for 50 simulation replicates. Each simulated data set had $m = 100$ instances, $p = 1000$ features with 100 functional, and balanced class groups with 50 'case' and 50 'control'. Functional features included 50 with main effect ($\text{bias}_{\text{main}} = 0.8$) only and the remaining 50 were involved in network interactions ($\text{bias}_{\text{int}} = 0.4$) and had no main effect. Data was first split into training and test (70% train/30% test) sets and 5 folds were used for inner and outer training loops. The top 30% of features, ranked in decreasing order of importance, were selected within each inner training fold. Matthew's Correlation Coefficient (MCC), precision, and recall were calculated based on detection of functional features. Training and validation represent the balanced classification accuracy of the random forest model that was fit using consensus features from cnCV on full training data (70% of full dataset) and independent test data (30% of full dataset), respectively. Comparisons were made with Mann-Whitney U test.

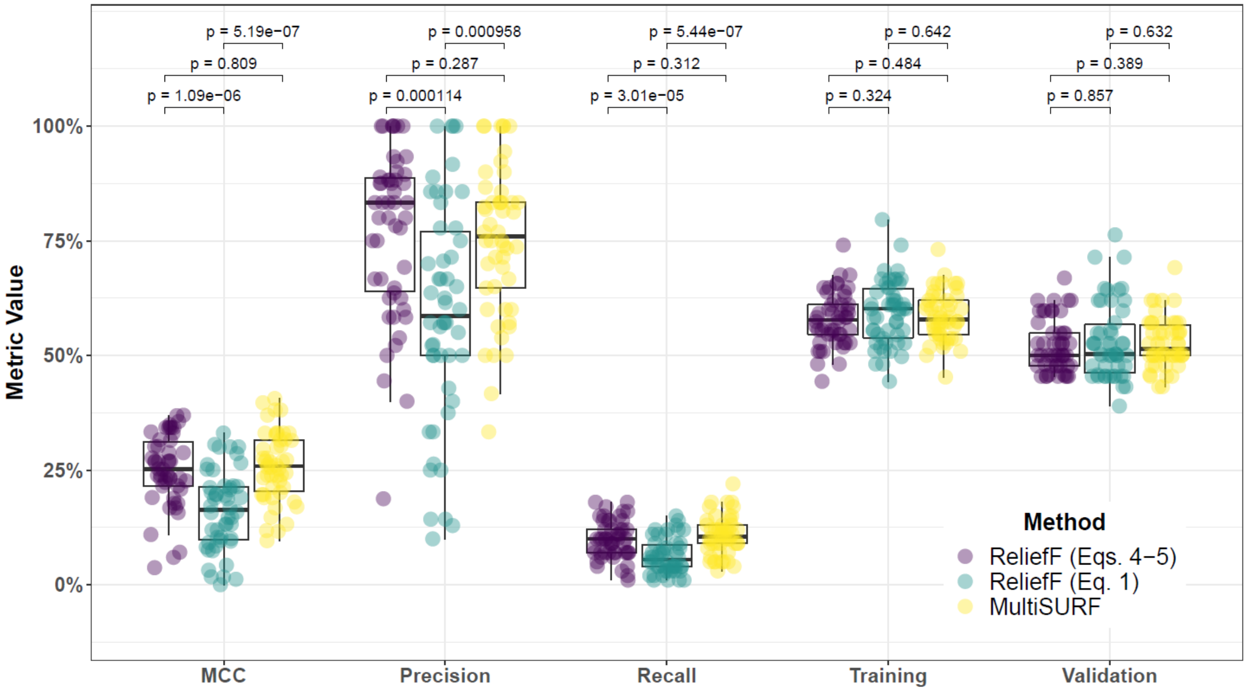


**Supplementary Fig. 31. Performance comparison for hit-miss-k (Eqs. 4 – 5), non-adjusted fixed-k (Eq. 1), and MultiSURF with ReliefF feature scoring and consensus-features nested Cross-Validation (cnCV) on imbalanced data.** Performance of feature selection was measured for 50 simulation replicates. Each simulated data set had $m = 100$ instances, $p = 1000$ features with 100 functional, and imbalanced class groups with 25 'case' and 75 'control'. Functional features included 50 with main effect ($\text{bias}_{\text{main}} = 0.8$) only and the remaining 50 were involved in network interactions ($\text{bias}_{\text{int}} = 0.4$) and had no main effect. Data was first split into training and test (70% train/30% test) sets and 5 folds were used for inner and outer training loops. The top 30% of features, ranked in decreasing order of importance, were selected within each inner training fold. Matthew's Correlation Coefficient (MCC), precision, and recall were calculated based on detection of functional features. Training and validation represent the balanced classification accuracy of the random forest model that was fit using consensus features from cnCV on full training data (70% of full dataset) and independent test data (30% of full dataset), respectively. Comparisons were made with Mann-Whitney U test.

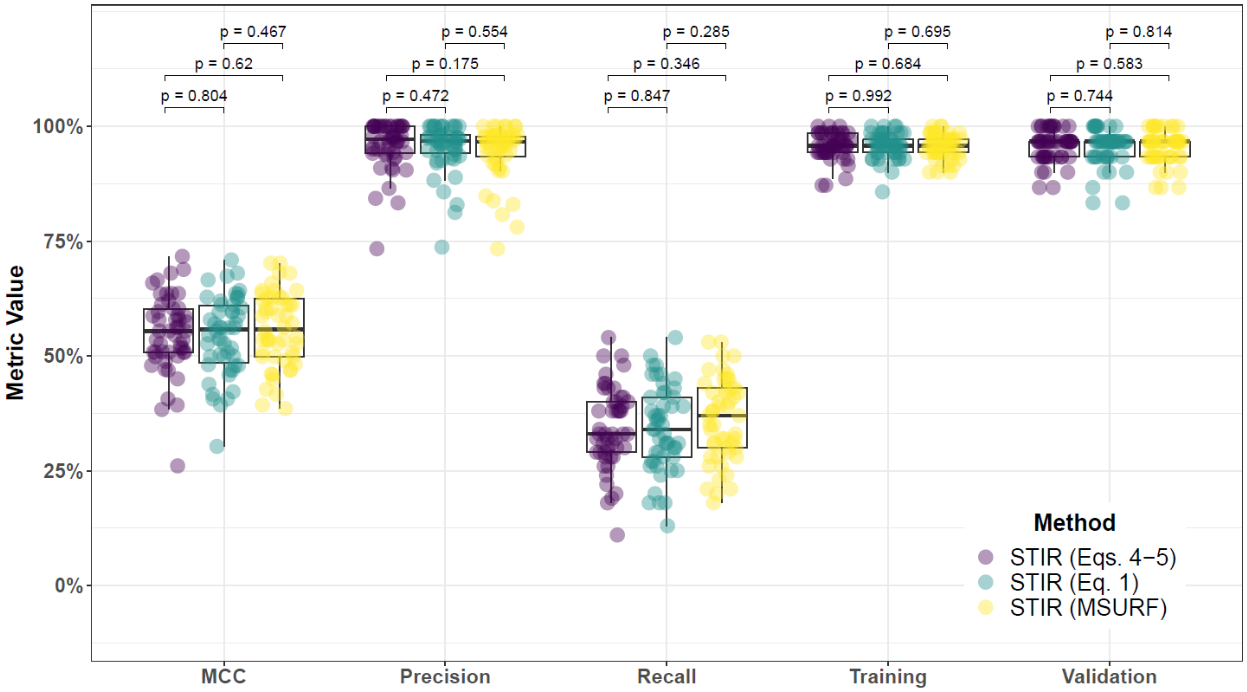


**Supplementary Fig. 32. Performance comparison for hit-miss-k (Eqs. 4 – 5), non-adjusted fixed-k (Eq. 1), and MultiSURF with STIR feature scoring and consensus-features nested Cross-Validation (cnCV) on balanced data.** Performance of feature selection was measured for 50 simulation replicates. Each simulated data set had $m = 100$ instances, $p = 1000$ features with 100 functional, and balanced class groups with 50 'case' and 50 'control'. Functional features included 50 with main effect ($\text{bias}_{\text{main}} = 0.8$) only and the remaining 50 were involved in network interactions ($\text{bias}_{\text{int}} = 0.4$) and had no main effect. Data was first split into training and test (70% train/30% test) sets and 5 folds were used for inner and outer training loops. The top 30% of features, ranked in decreasing order of importance, were selected within each inner training fold. Matthew's Correlation Coefficient (MCC), precision, and recall were calculated based on detection of functional features. Training and validation represent the balanced classification accuracy of the random forest model that was fit using consensus features from cnCV on full training data (70% of full dataset) and independent test data (30% of full dataset), respectively. Comparisons were made with Mann-Whitney U test.

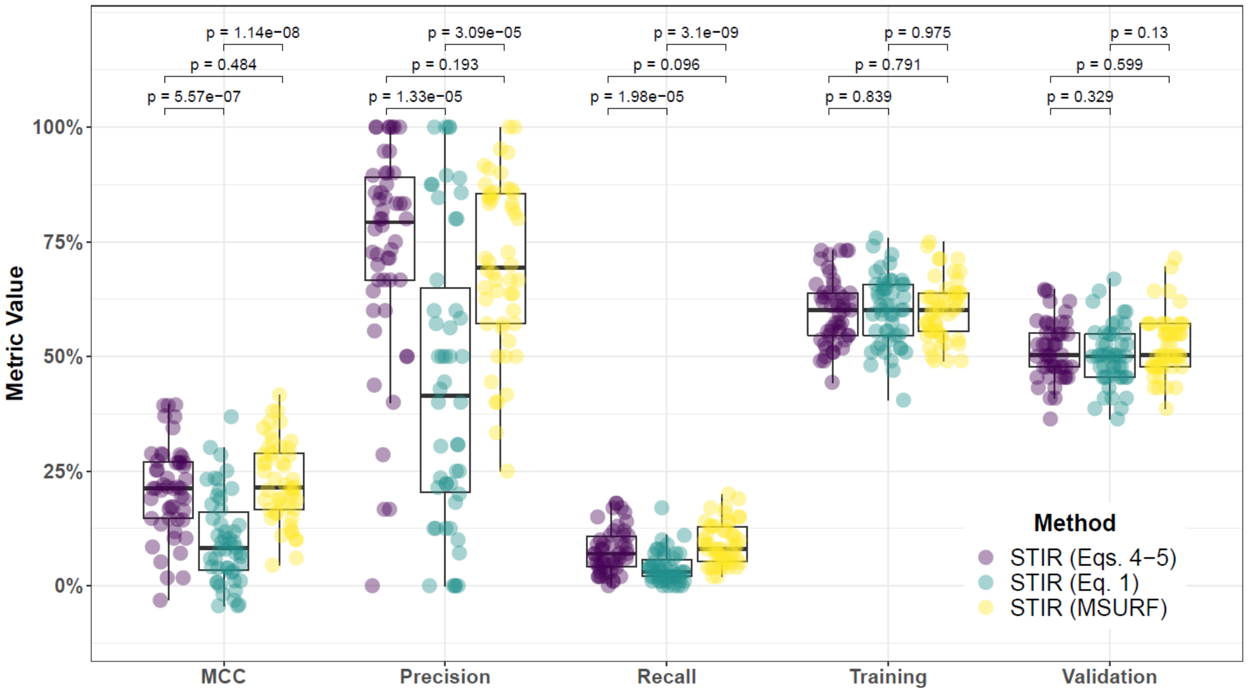


**Supplementary Fig. 33. Performance comparison for hit-miss-k (Eqs. 4 – 5), non-adjusted fixed-k (Eq. 1), and MultiSURF with STIR feature scoring and consensus-features nested Cross-Validation (cnCV) on imbalanced data.** Performance of feature selection was measured for 50 simulation replicates. Each simulated data set had $m = 100$ instances, $p = 1000$ features with 100 functional, and imbalanced class groups with 25 'case' and 75 'control'. Functional features included 50 with main effect ($\text{bias}_{\text{main}} = 0.8$) only and the remaining 50 were involved in network interactions ($\text{bias}_{\text{int}} = 0.4$) and had no main effect. Data was first split into training and test (70% train/30% test) sets and 5 folds were used for inner and outer training loops. The top 30% of features, ranked in decreasing order of importance, were selected within each inner training fold. Matthew's Correlation Coefficient (MCC), precision, and recall were calculated based on detection of functional features. Training and validation represent the balanced classification accuracy of the random forest model that was fit using consensus features from cnCV on full training data (70% of full dataset) and independent test data (30% of full dataset), respectively. Comparisons were made with Mann-Whitney U test.

##### 4.3 Comparing imbalance-adjusted fixed-k, random forest, and ridge regression

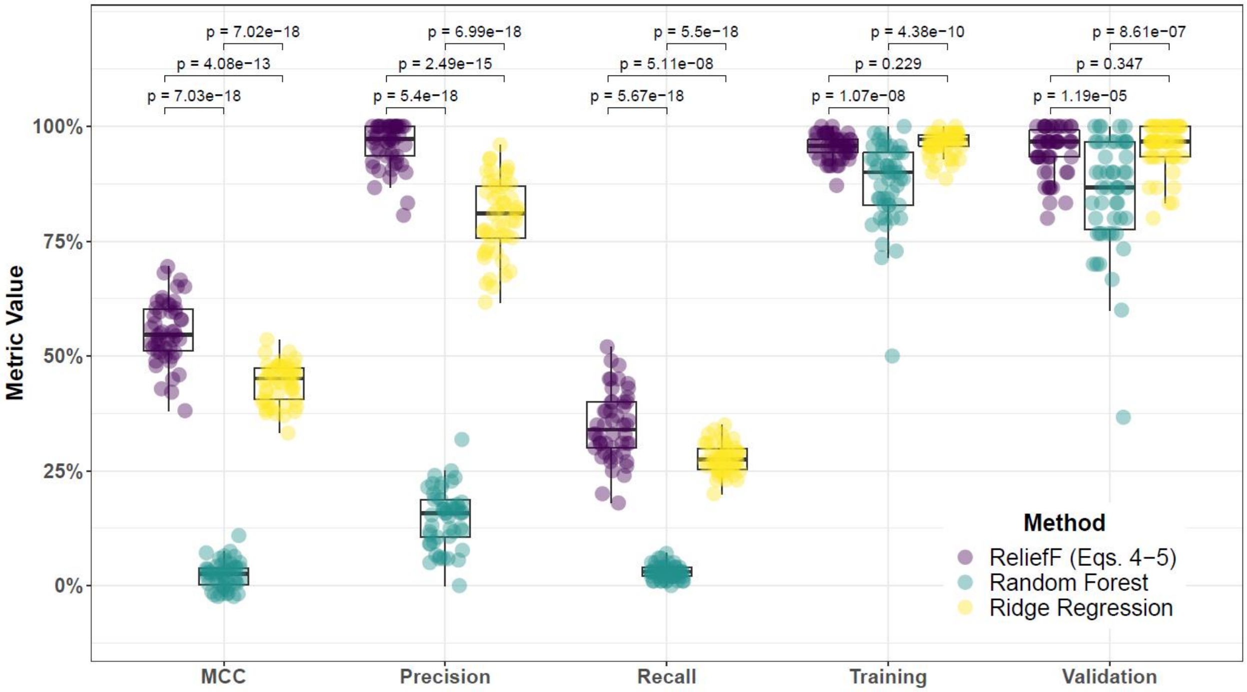


**Supplementary Fig. 34. Performance comparison for hit-miss-k (Eqs. 4 – 5) ReliefF, Random Forest (RF), and Ridge Regression (RR) within consensus-features nested Cross-Validation (cnCV) on balanced data.** Performance of feature selection was measured for 50 simulation replicates. Each simulated data set had $m = 100$ instances, $p = 1000$ features with 100 functional, and balanced class groups with 50 'case' and 50 'control'. Functional features included 50 with main effect ($\text{bias}_{\text{main}} = 0.8$) only and the remaining 50 were involved in network interactions ($\text{bias}_{\text{int}} = 0.4$) and had no main effect. Data was first split into training and test (70% train/30% test) sets and 5 folds were used for inner and outer training loops. The top 30% of features, ranked in decreasing order of importance, were selected within each inner training fold. Matthew's Correlation Coefficient (MCC), precision, and recall were calculated based on detection of functional features. Training and validation represent the balanced classification accuracy of the random forest model that was fit using consensus features from cnCV on full training data (70% of full dataset) and independent test data (30% of full dataset), respectively. Comparisons were made with Mann-Whitney U test.

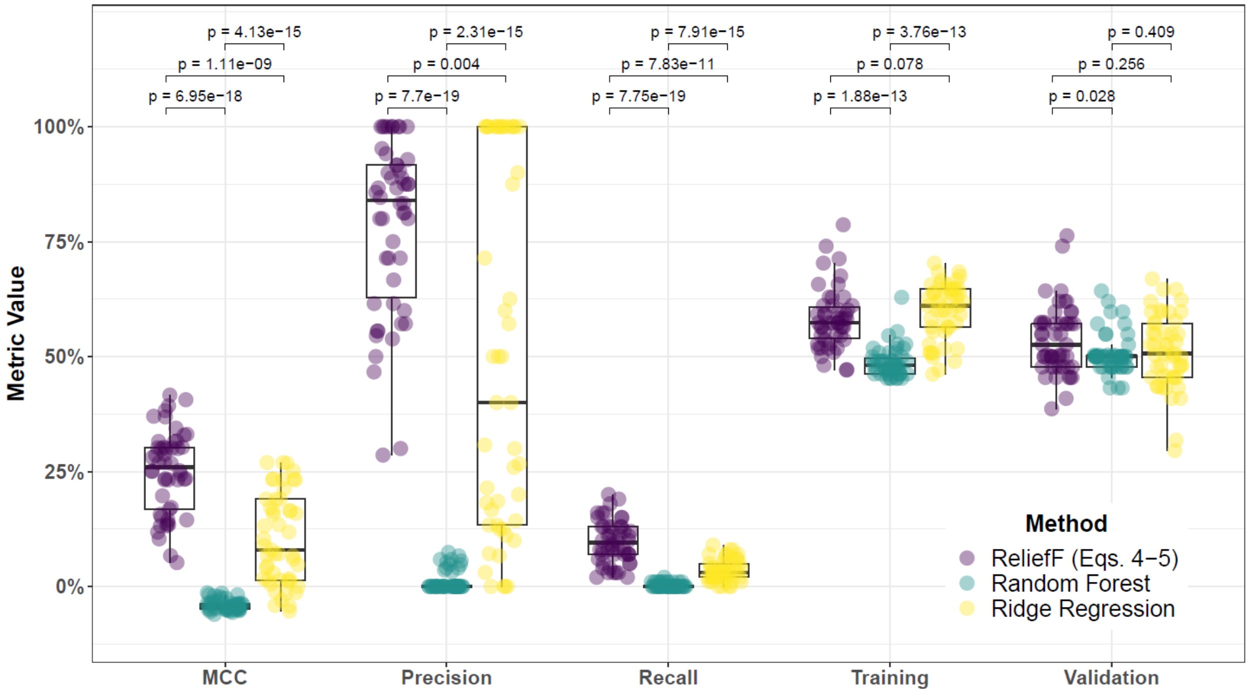


**Supplementary Fig. 35. Performance comparison for hit-miss-k (Eqs. 4 – 5) ReliefF, Random Forest (RF), and Ridge Regression (RR) within consensus-features nested Cross-Validation (cnCV) on imbalanced data.** Performance of feature selection was measured for 50 simulation replicates. Each simulated data set had $m = 100$ instances, $p = 1000$ features with 100 functional, and imbalanced class groups with 25 'case' and 75 'control'. Functional features included 50 with main effect ($\text{bias}_{\text{main}} = 0.8$) only and the remaining 50 were involved in network interactions ($\text{bias}_{\text{int}} = 0.4$) and had no main effect. Data was first split into training and test (70% train/30% test) sets and 5 folds were used for inner and outer training loops. The top 30% of features, ranked in decreasing order of importance, were selected within each inner training fold. Matthew's Correlation Coefficient (MCC), precision, and recall were calculated based on detection of functional features. Training and validation represent the balanced classification accuracy of the random forest model that was fit using consensus features from cnCV on full training data (70% of full dataset) and independent test data (30% of full dataset), respectively. Comparisons were made with Mann-Whitney U test.

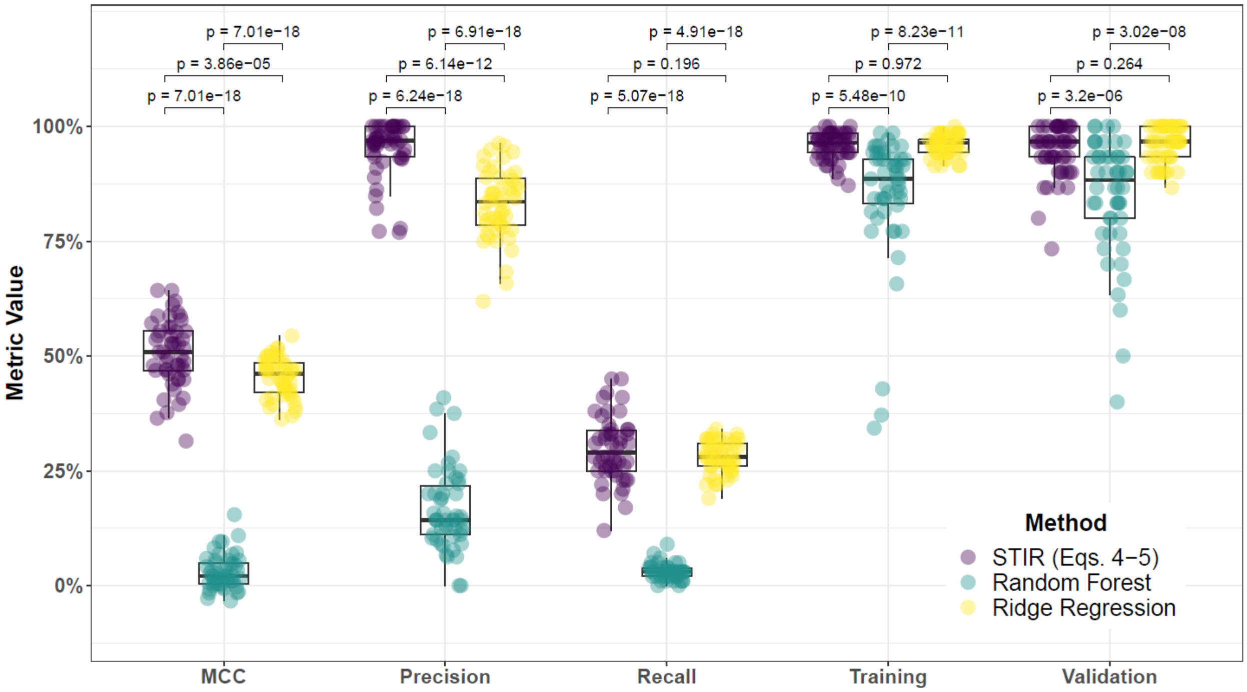


**Supplementary Fig. 36. Performance comparison for hit-miss-k (Eqs. 4 – 5) STIR, Random Forest (RF), and Ridge Regression (RR) within consensus-features nested Cross-Validation (cnCV) on balanced data.** Performance of feature selection was measured for 50 simulation replicates. Each simulated data set had $m = 100$ instances, $p = 1000$ features with 100 functional, and balanced class groups with 50 'case' and 50 'control'. Functional features included 50 with main effect ($\text{bias}_{\text{main}} = 0.8$) only and the remaining 50 were involved in network interactions ($\text{bias}_{\text{int}} = 0.4$) and had no main effect. Data was first split into training and test (70% train/30% test) sets and 5 folds were used for inner and outer training loops. The top 30% of features, ranked in decreasing order of importance, were selected within each inner training fold. Matthew's Correlation Coefficient (MCC), precision, and recall were calculated based on detection of functional features. Training and validation represent the balanced classification accuracy of the random forest model that was fit using consensus features from cnCV on full training data (70% of full dataset) and independent test data (30% of full dataset), respectively. Comparisons were made with Mann-Whitney U test.

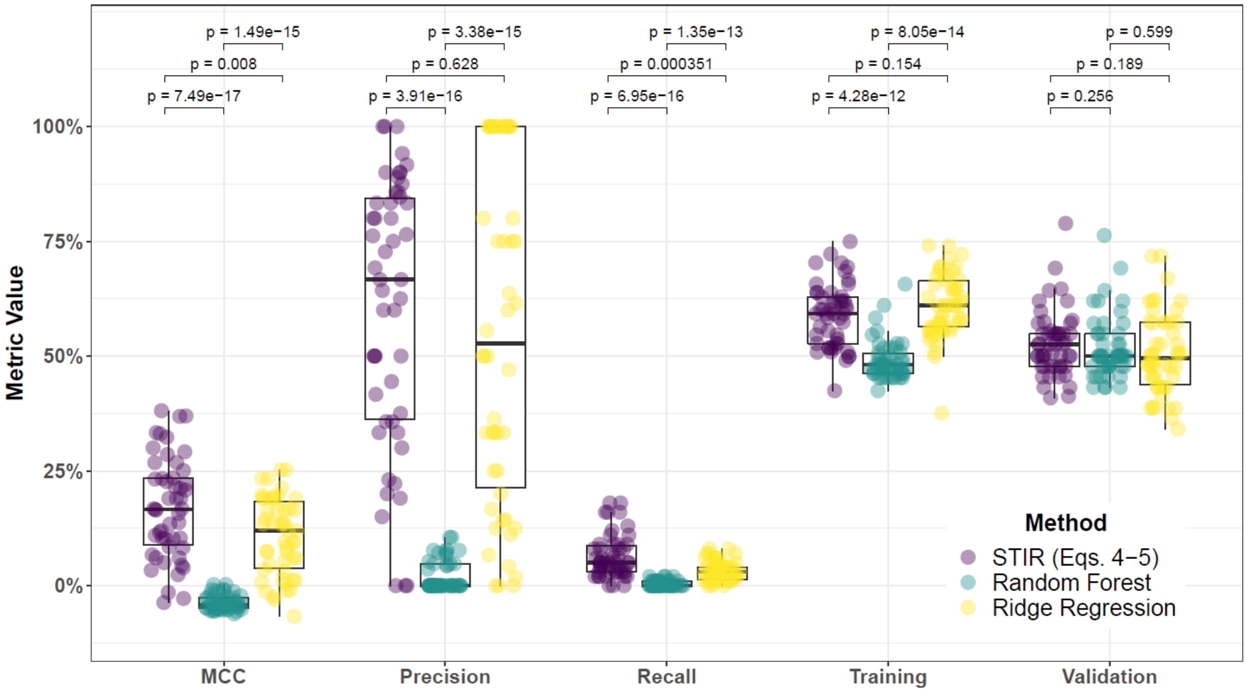


**Supplementary Fig. 37. Performance comparison for hit-miss-k (Eqs. 4 – 5) STIR, Random Forest (RF), and Ridge Regression (RR) within consensus-features nested Cross-Validation (cnCV) on imbalanced data.** Performance of feature selection was measured for 50 simulation replicates. Each simulated data set had $m = 100$ instances, $p = 1000$ features with 100 functional, and imbalanced class groups with 25 'case' and 75 'control'. Functional features included 50 with main effect ($\text{bias}_{\text{main}} = 0.8$) only and the remaining 50 were involved in network interactions ($\text{bias}_{\text{int}} = 0.4$) and had no main effect. Data was first split into training and test (70% train/30% test) sets and 5 folds were used for inner and outer training loops. The top 30% of features, ranked in decreasing order of importance, were selected within each inner training fold. Matthew's Correlation Coefficient (MCC), precision, and recall were calculated based on detection of functional features. Training and validation represent the balanced classification accuracy of the random forest model that was fit using consensus features from cnCV on full training data (70% of full dataset) and independent test data (30% of full dataset), respectively. Comparisons were made with Mann-Whitney U test.

### 5 Feature selection performance comparisons within consensus-features nested cross-validation (cnCV): 75% interaction effect/25% main effect

#### 5.1 Comparing imbalance-adjusted fixed-k, VWOK, and kPCA

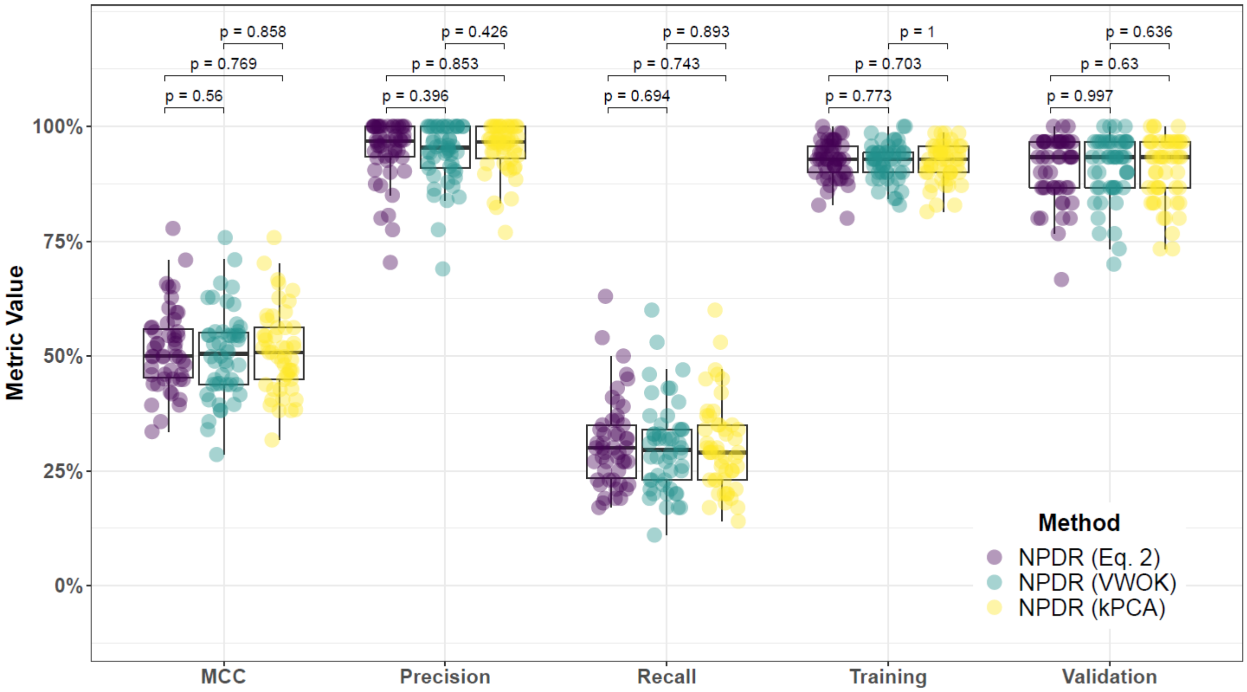


**Supplementary Fig. 38. Performance comparison for minority-class-k (Eq. 2), VWOK (Eq. 3), and kPCA with NPDR feature scoring and consensus-features nested Cross-Validation (cnCV) on balanced data.** Performance of feature selection was measured for 50 simulation replicates. Each simulated data set had $m = 100$ instances and $p = 1000$ features with 100 functional. Each simulated data set had balanced class groups with 50 'case' and 50 'control'. Functional features included 25 with main effect ($\text{bias}_{\text{main}} = 0.8$) only and the remaining 75 were involved in network interactions ($\text{bias}_{\text{int}} = 0.4$) and had no main effect. Matthew's Correlation Coefficient (MCC), precision, and recall were calculated based on detection of functional features (i.e. true positives). Training and validation represent the balanced classification accuracy of the random forest model that was fit using consensus features from the cnCV on full training data (70 samples) and independent test data (30 samples), respectively.

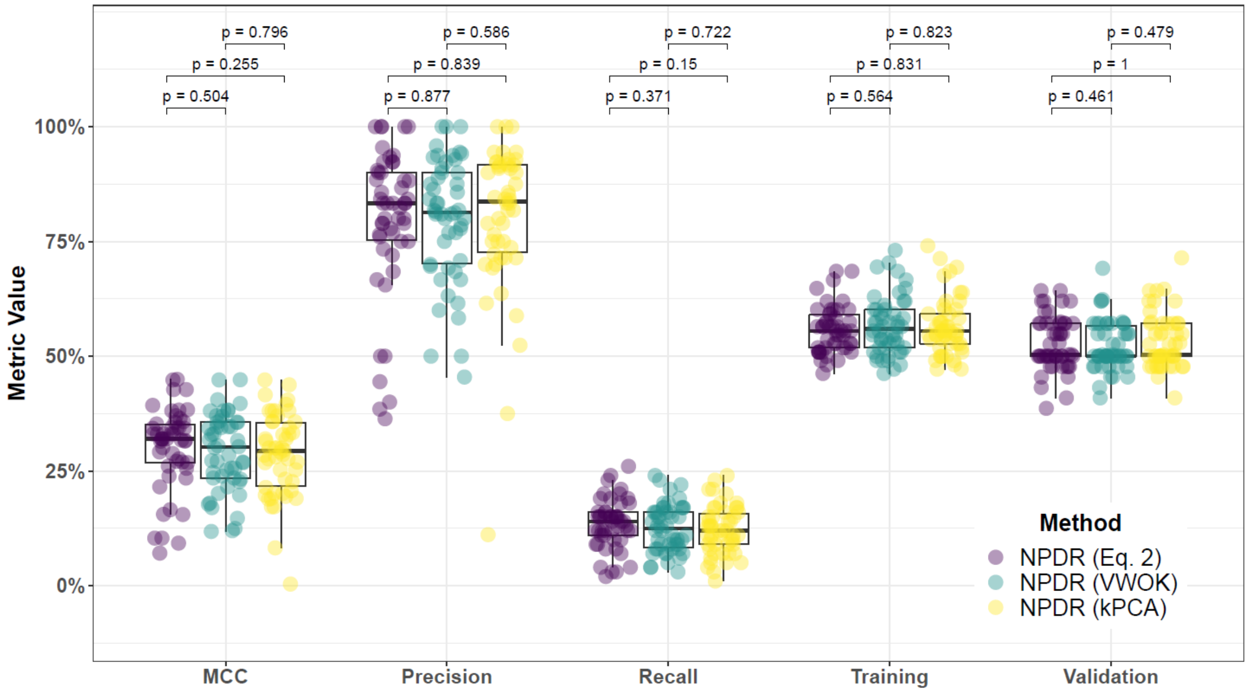


**Supplementary Fig. 39. Performance comparison for minority-class-k (Eq. 2), VWOK (Eq. 3), and kPCA with NPDR feature scoring and consensus-features nested Cross-Validation (cnCV) on imbalanced data.** Performance of feature selection was measured for 50 simulation replicates. Each simulated data set had $m = 100$ instances and $p = 1000$ features with 100 functional. Each simulated data set had imbalanced class groups with 25 'case' and 75 'control'. Functional features included 25 with main effect ($\text{bias}_{\text{main}} = 0.8$) only and the remaining 75 were involved in network interactions ($\text{bias}_{\text{int}} = 0.4$) and had no main effect. Matthew's Correlation Coefficient (MCC), precision, and recall were calculated based on detection of functional features (i.e. true positives). Training and validation represent the balanced classification accuracy of the random forest model that was fit using consensus features from the cnCV on full training data (70 samples) and independent test data (30 samples), respectively.

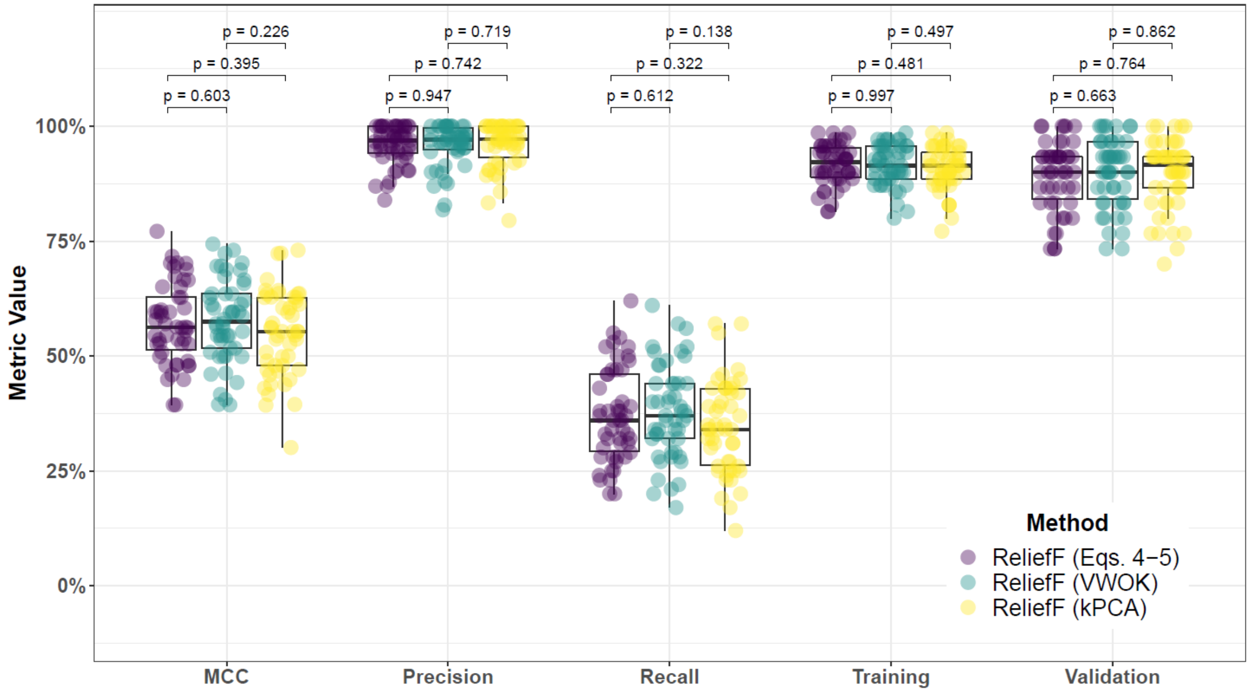


**Supplementary Fig. 40. Performance comparison for hit-miss-k (Eqs. 4 – 5), VWOK (Eqs. 6 – 7), and kPCA with ReliefF feature scoring and consensus-features nested Cross-Validation (cnCV) on balanced data.** Performance of feature selection was measured for 50 simulation replicates. Each simulated data set had $m = 100$ instances and $p = 1000$ features with 100 functional. Each simulated data set had balanced class groups with 50 'case' and 50 'control'. Functional features included 25 with main effect ($\text{bias}_{\text{main}} = 0.8$) only and the remaining 75 were involved in network interactions ($\text{bias}_{\text{int}} = 0.4$) and had no main effect. Matthew's Correlation Coefficient (MCC), precision, and recall were calculated based on detection of functional features (i.e. true positives). Training and validation represent the balanced classification accuracy of the random forest model that was fit using consensus features from the cnCV on full training data (70 samples) and independent test data (30 samples), respectively.

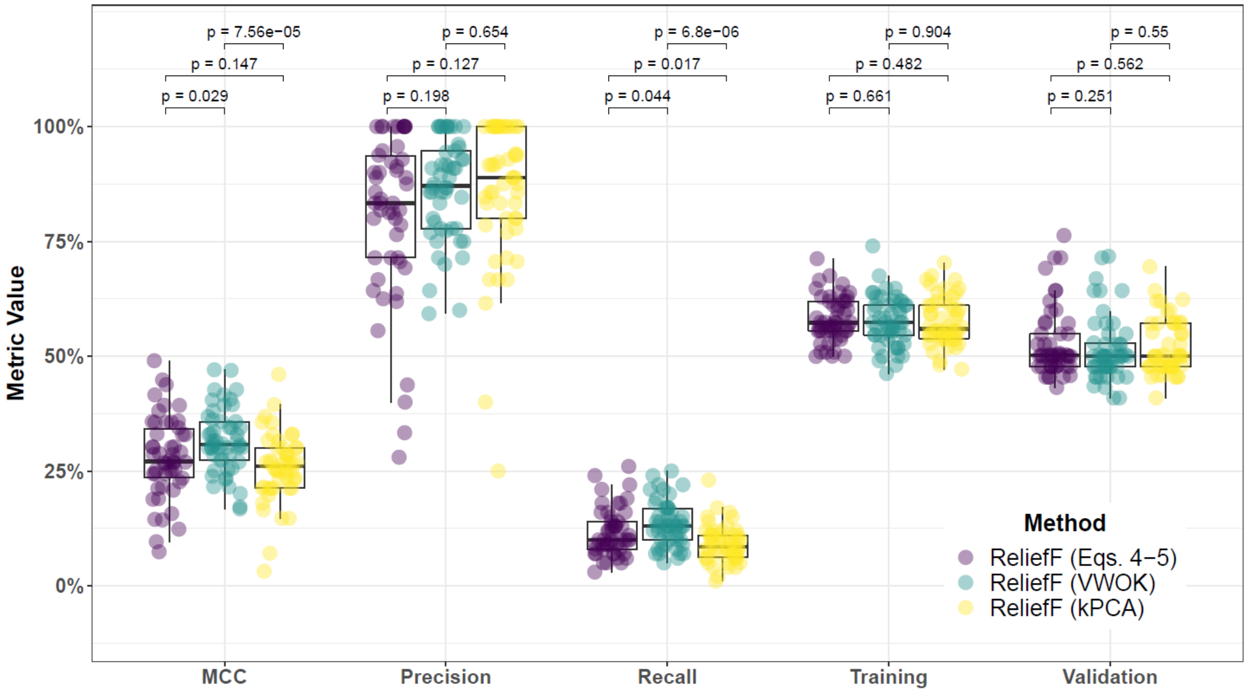


**Supplementary Fig. 41. Performance comparison for hit-miss-k (Eqs. 4 – 5), VWOK (Eqs. 6 – 7), and kPCA with ReliefF feature scoring and consensus-features nested Cross-Validation (cnCV) on imbalanced data.** Performance of feature selection was measured for 50 simulation replicates. Each simulated data set had $m = 100$ instances and $p = 1000$ features with 100 functional. Each simulated data set had imbalanced class groups with 25 'case' and 75 'control'. Functional features included 25 with main effect ($\text{bias}_{\text{main}} = 0.8$) only and the remaining 75 were involved in network interactions ($\text{bias}_{\text{int}} = 0.4$) and had no main effect. Matthew's Correlation Coefficient (MCC), precision, and recall were calculated based on detection of functional features (i.e. true positives). Training and validation represent the balanced classification accuracy of the random forest model that was fit using consensus features from the cnCV on full training data (70 samples) and independent test data (30 samples), respectively.

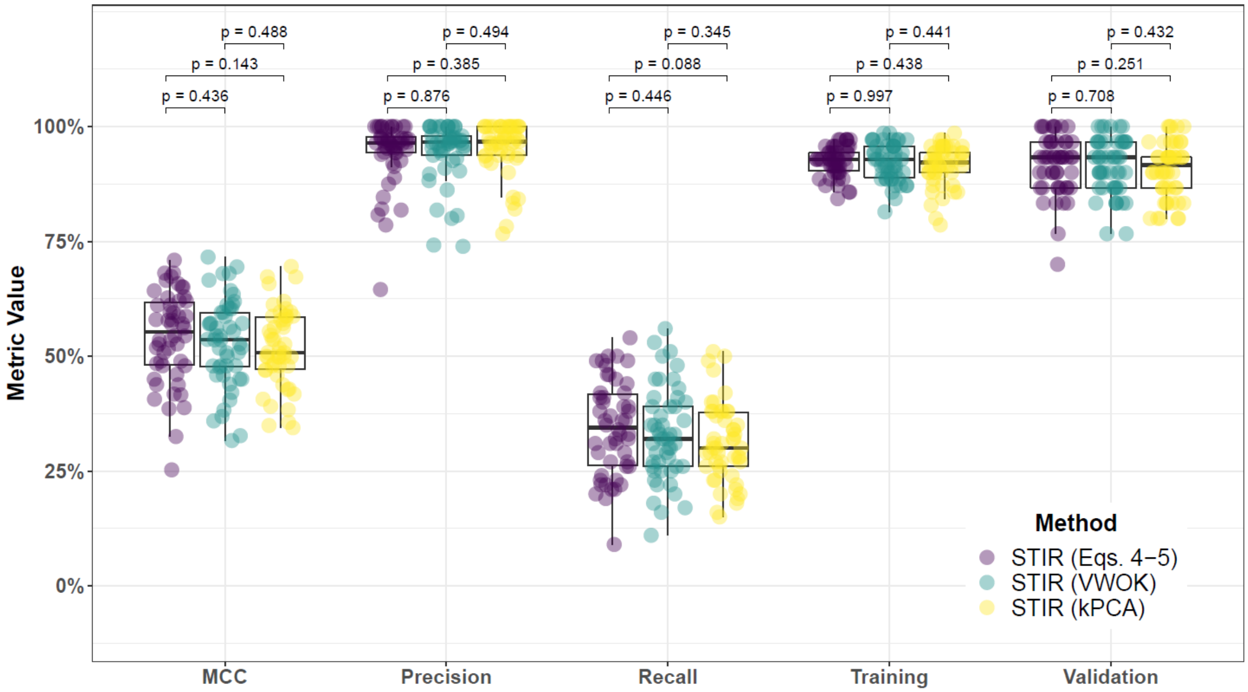


**Supplementary Fig. 42. Performance comparison for hit-miss-k (Eqs. 4 – 5), VWOK (Eqs. 6 – 7), and kPCA with STIR feature scoring and consensus-features nested Cross-Validation (cnCV) on balanced data.** Performance of feature selection was measured for 50 simulation replicates. Each simulated data set had $m = 100$ instances and $p = 1000$ features with 100 functional. Each simulated data set had balanced class groups with 50 'case' and 50 'control'. Functional features included 25 with main effect ($\text{bias}_{\text{main}} = 0.8$) only and the remaining 75 were involved in network interactions ($\text{bias}_{\text{int}} = 0.4$) and had no main effect. Matthew's Correlation Coefficient (MCC), precision, and recall were calculated based on detection of functional features (i.e. true positives). Training and validation represent the balanced classification accuracy of the random forest model that was fit using consensus features from the cnCV on full training data (70 samples) and independent test data (30 samples), respectively.

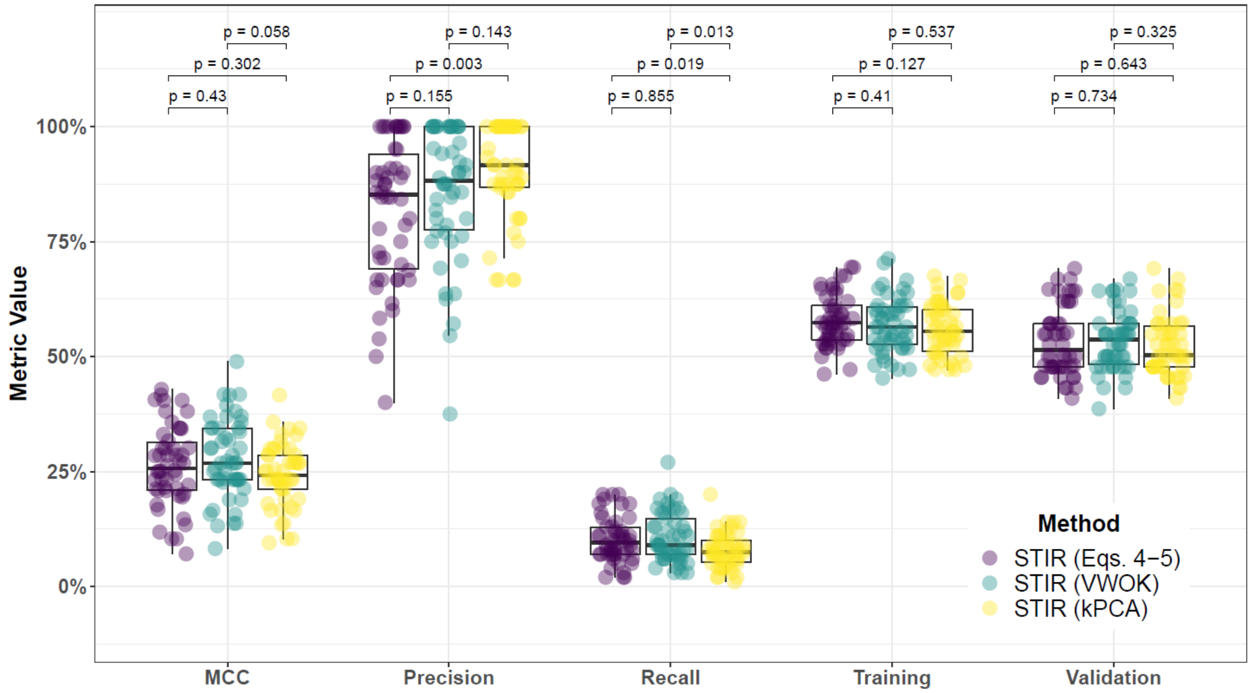


**Supplementary Fig. 43. Performance comparison for hit-miss-k (Eqs. 4 – 5), VWOK (Eqs. 6 – 7), and kPCA with STIR feature scoring and consensus-features nested Cross-Validation (cnCV) on imbalanced data.** Performance of feature selection was measured for 50 simulation replicates. Each simulated data set had $m = 100$ instances and $p = 1000$ features with 100 functional. Each simulated data set had imbalanced class groups with 25 'case' and 75 'control'. Functional features included 25 with main effect ($\text{bias}_{\text{main}} = 0.8$) only and the remaining 75 were involved in network interactions ($\text{bias}_{\text{int}} = 0.4$) and had no main effect. Matthew's Correlation Coefficient (MCC), precision, and recall were calculated based on detection of functional features (i.e. true positives). Training and validation represent the balanced classification accuracy of the random forest model that was fit using consensus features from the cnCV on full training data (70 samples) and independent test data (30 samples), respectively.

##### 5.2 Comparing imbalance-adjusted fixed-k, regular fixed-k, and MultiSURF

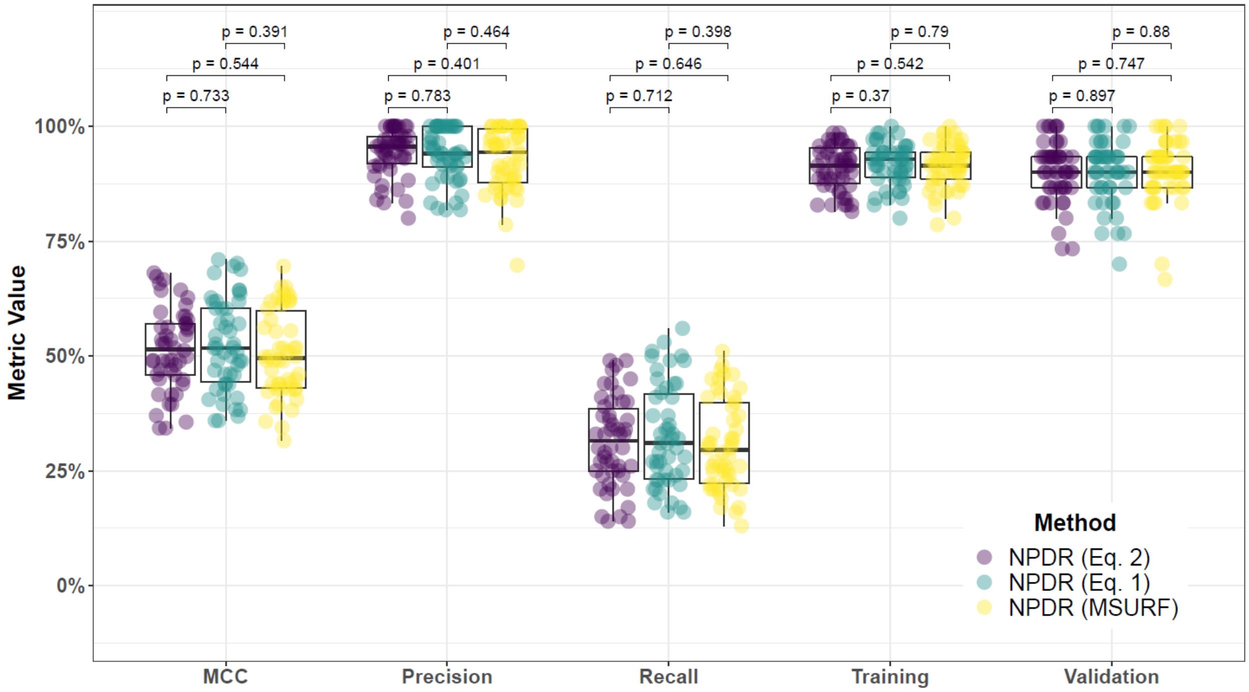


**Supplementary Fig. 44. Performance comparison for minority-class-k (Eq. 2), non-adjusted fixed-k (Eq. 1), and MultiSURF with NPDR feature scoring and consensus-features nested Cross-Validation (cnCV) on balanced data.** Performance of feature selection was measured for 50 simulation replicates. Each simulated data set had $m = 100$ instances and $p = 1000$ features with 100 functional. Each simulated data set had balanced class groups with 50 'case' and 50 'control'. Functional features included 25 with main effect ($\text{bias}_{\text{main}} = 0.8$) only and the remaining 75 were involved in network interactions ($\text{bias}_{\text{int}} = 0.4$) and had no main effect. Matthew's Correlation Coefficient (MCC), precision, and recall were calculated based on detection of functional features (i.e. true positives). Training and validation represent the balanced classification accuracy of the random forest model that was fit using consensus features from the cnCV on full training data (70 samples) and independent test data (30 samples), respectively.

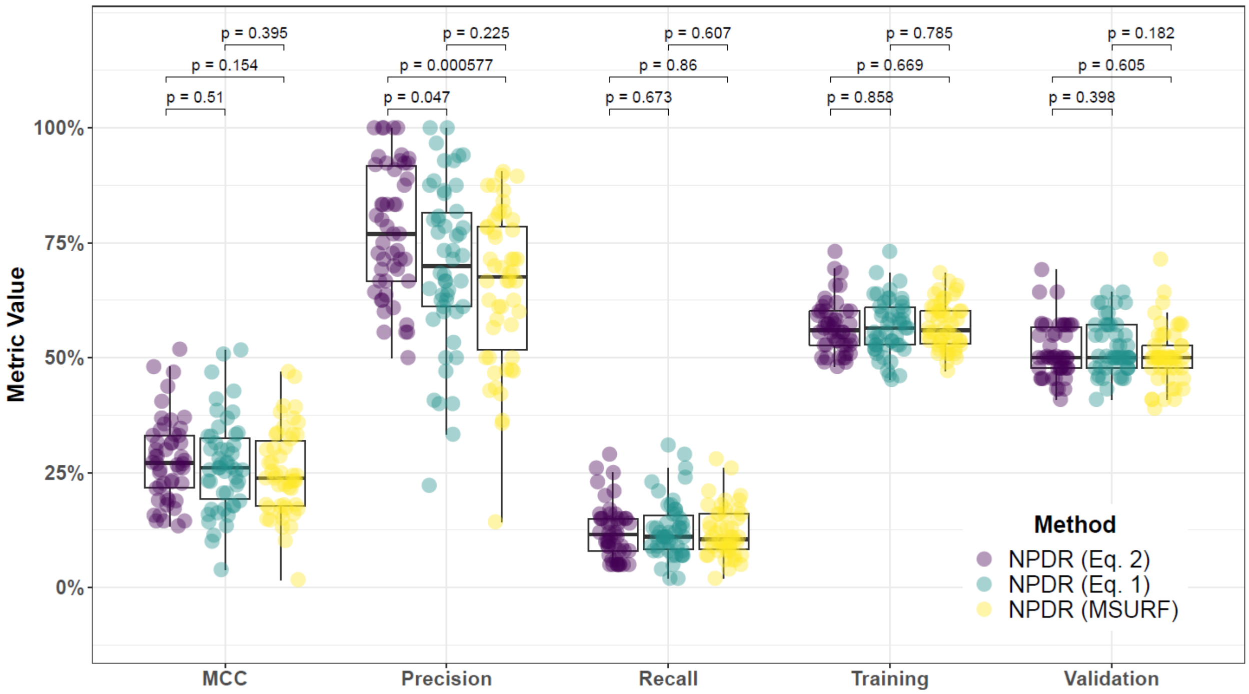


**Supplementary Fig. 45. Performance comparison for minority-class-k (Eq. 2), non-adjusted fixed-k (Eq. 1), and MultiSURF with NPDR feature scoring and consensus-features nested Cross-Validation (cnCV) on imbalanced data.** Performance of feature selection was measured for 50 simulation replicates. Each simulated data set had $m = 100$ instances and $p = 1000$ features with 100 functional. Each simulated data set had imbalanced class groups with 25 'case' and 75 'control'. Functional features included 25 with main effect ($\text{bias}_{\text{main}} = 0.8$) only and the remaining 75 were involved in network interactions ($\text{bias}_{\text{int}} = 0.4$) and had no main effect. Matthew's Correlation Coefficient (MCC), precision, and recall were calculated based on detection of functional features (i.e. true positives). Training and validation represent the balanced classification accuracy of the random forest model that was fit using consensus features from the cnCV on full training data (70 samples) and independent test data (30 samples), respectively.

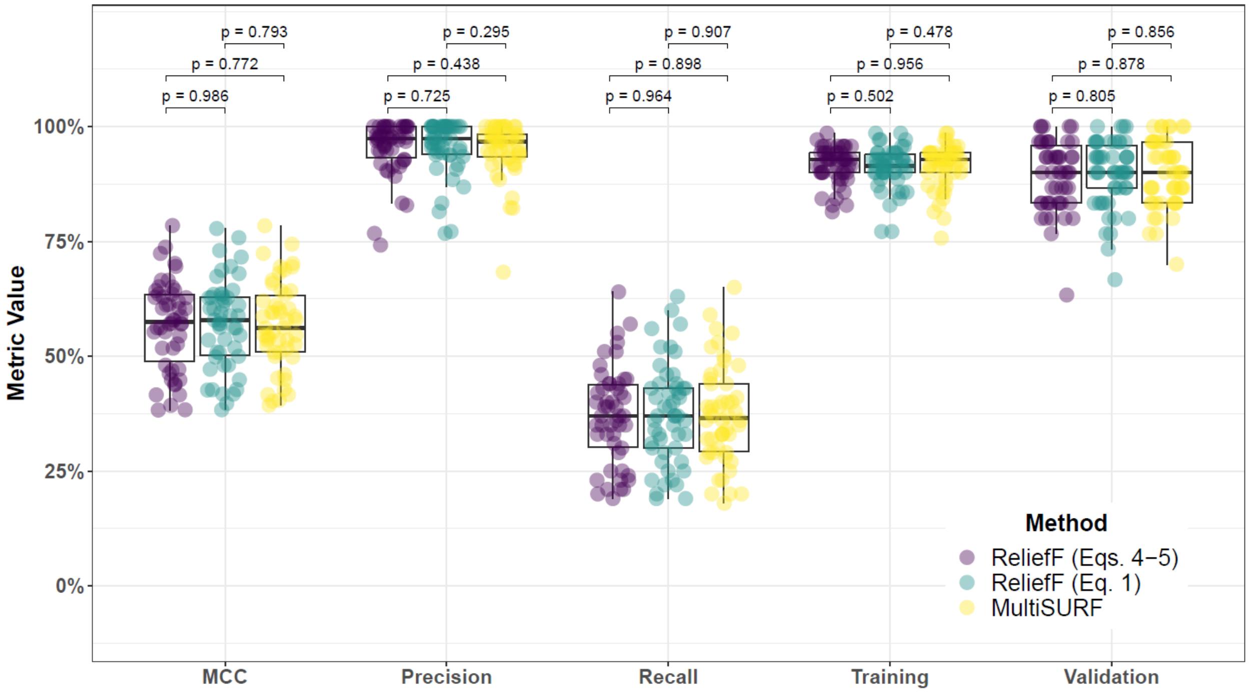


**Supplementary Fig. 46. Performance comparison for hit-miss-k (Eqs. 4 – 5) ReliefF, non-adjusted fixed-k (Eq. 1) ReliefF, and MultiSURF and consensus-features nested Cross-Validation (cnCV) on balanced data.** Performance of feature selection was measured for 50 simulation replicates. Each simulated data set had $m = 100$ instances and $p = 1000$ features with 100 functional. Each simulated data set had balanced class groups with 50 'case' and 50 'control'. Functional features included 25 with main effect ($\text{bias}_{\text{main}} = 0.8$) only and the remaining 75 were involved in network interactions ($\text{bias}_{\text{int}} = 0.4$) and had no main effect. Matthew's Correlation Coefficient (MCC), precision, and recall were calculated based on detection of functional features (i.e. true positives). Training and validation represent the balanced classification accuracy of the random forest model that was fit using consensus features from the cnCV on full training data (70 samples) and independent test data (30 samples), respectively.

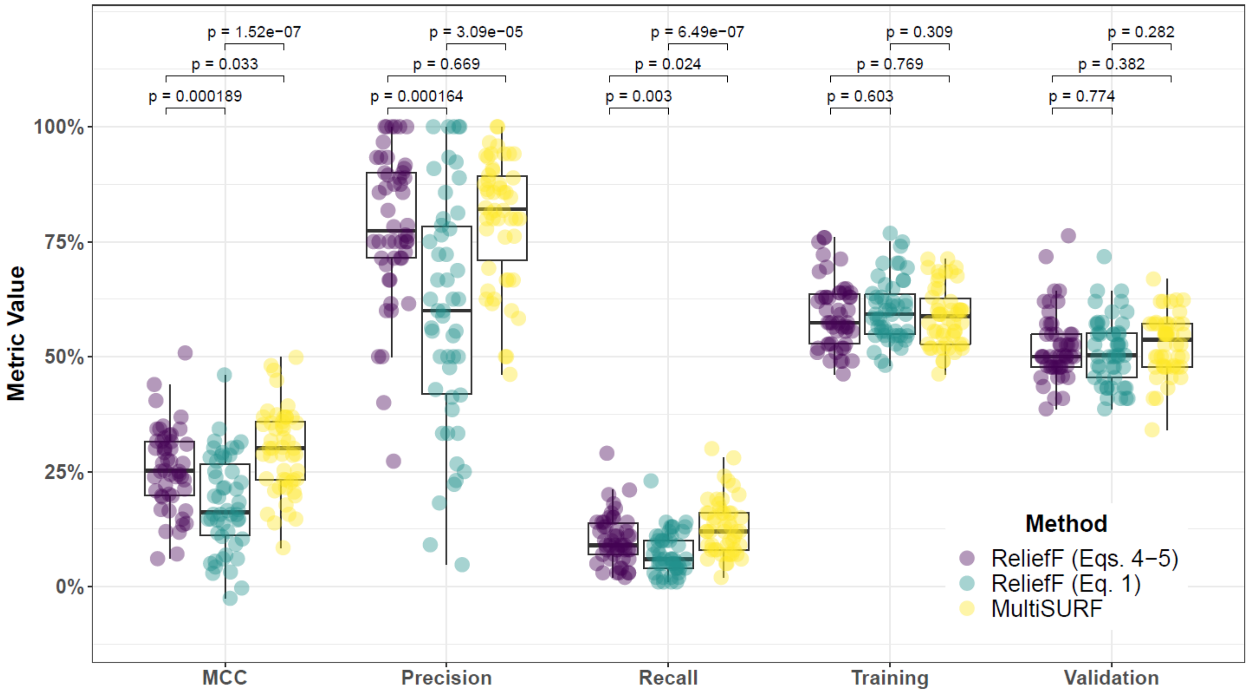


**Supplementary Fig. 47. Performance comparison for hit-miss-k (Eqs. 4 – 5) ReliefF, non-adjusted fixed-k (Eq. 1) ReliefF, and MultiSURF and consensus-features nested Cross-Validation (cnCV) on imbalanced data.** Performance of feature selection was measured for 50 simulation replicates. Each simulated data set had $m = 100$ instances and $p = 1000$ features with 100 functional. Each simulated data set had imbalanced class groups with 25 'case' and 75 'control'. Functional features included 25 with main effect ($\text{bias}_{\text{main}} = 0.8$) only and the remaining 75 were involved in network interactions ($\text{bias}_{\text{int}} = 0.4$) and had no main effect. Matthew's Correlation Coefficient (MCC), precision, and recall were calculated based on detection of functional features (i.e. true positives). Training and validation represent the balanced classification accuracy of the random forest model that was fit using consensus features from the cnCV on full training data (70 samples) and independent test data (30 samples), respectively.

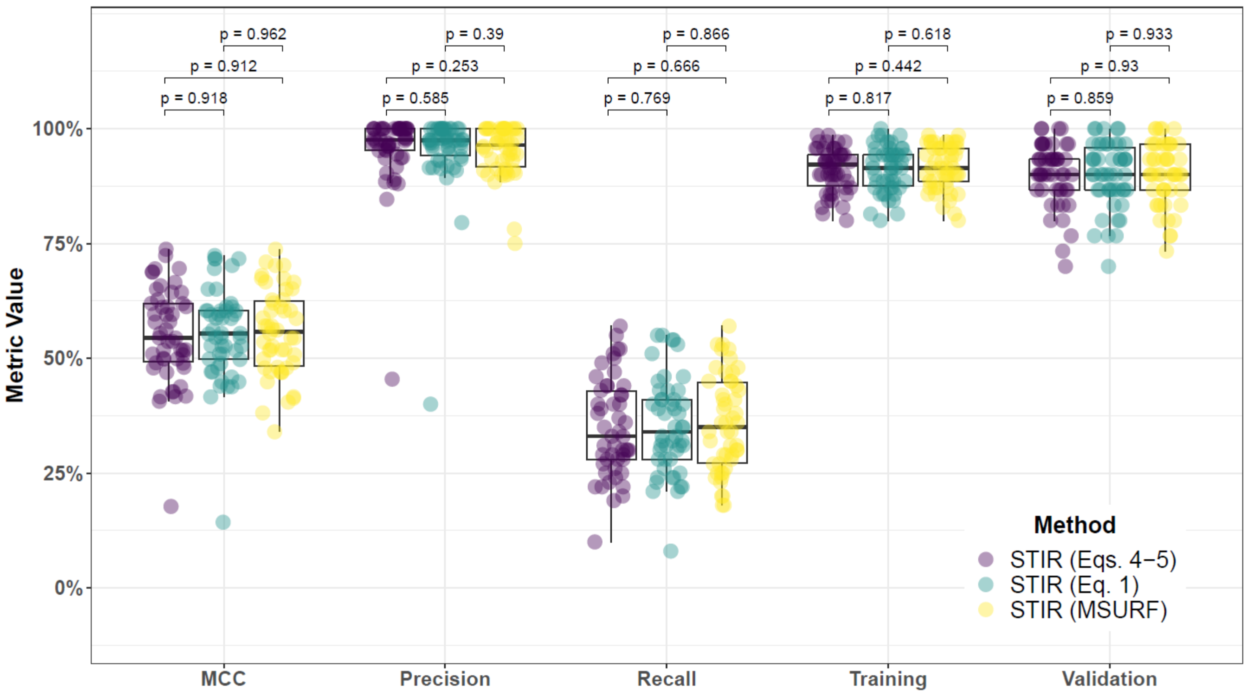


**Supplementary Fig. 48. Performance comparison for hit-miss-k (Eqs. 4 – 5), non-adjusted fixed-k (Eq. 1), and MultiSURF with STIR feature scoring and consensus-features nested Cross-Validation (cnCV) on balanced data.** Performance of feature selection was measured for 50 simulation replicates. Each simulated data set had $m = 100$ instances and $p = 1000$ features with 100 functional. Each simulated data set had balanced class groups with 50 'case' and 50 'control'. Functional features included 25 with main effect ($\text{bias}_{\text{main}} = 0.8$) only and the remaining 75 were involved in network interactions ($\text{bias}_{\text{int}} = 0.4$) and had no main effect. Matthew's Correlation Coefficient (MCC), precision, and recall were calculated based on detection of functional features (i.e. true positives). Training and validation represent the balanced classification accuracy of the random forest model that was fit using consensus features from the cnCV on full training data (70 samples) and independent test data (30 samples), respectively.

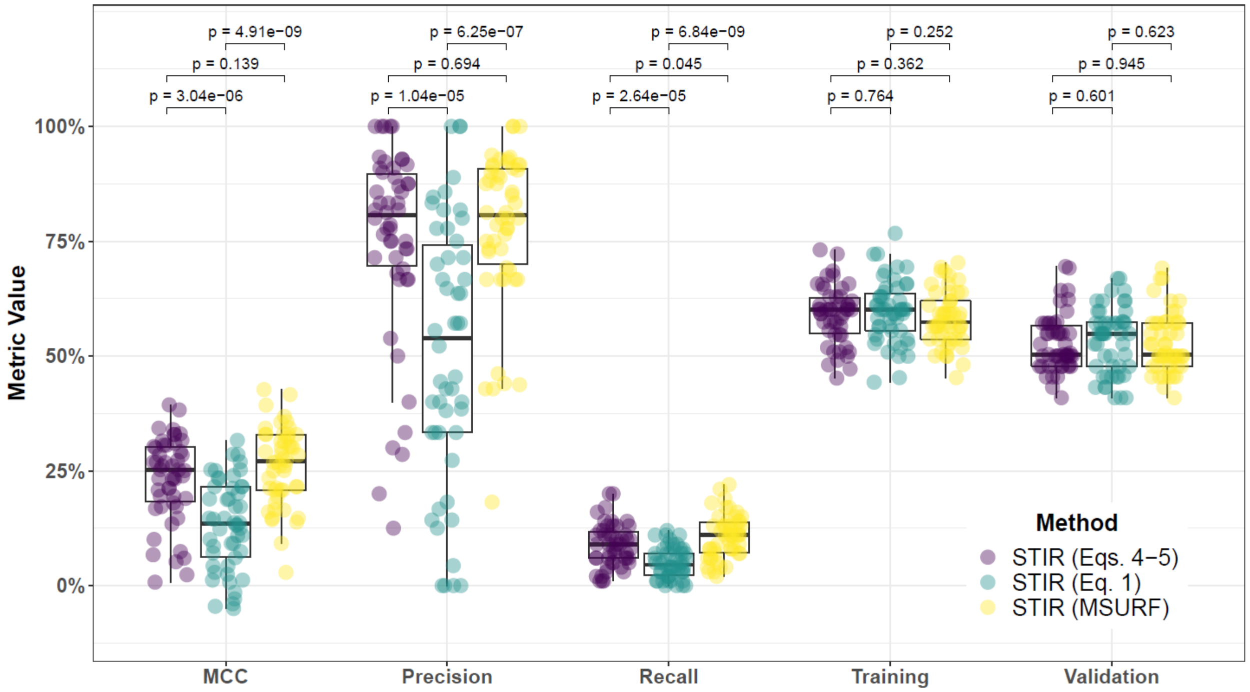


**Supplementary Fig. 49. Performance comparison for hit-miss-k (Eqs. 4 – 5), non-adjusted fixed-k (Eq. 1), and MultiSURF with STIR feature scoring and consensus-features nested Cross-Validation (cnCV) on imbalanced data.** Performance of feature selection was measured for 50 simulation replicates. Each simulated data set had $m = 100$ instances and $p = 1000$ features with 100 functional. Each simulated data set had imbalanced class groups with 25 'case' and 75 'control'. Functional features included 25 with main effect ($\text{bias}_{\text{main}} = 0.8$) only and the remaining 75 were involved in network interactions ($\text{bias}_{\text{int}} = 0.4$) and had no main effect. Matthew's Correlation Coefficient (MCC), precision, and recall were calculated based on detection of functional features (i.e. true positives). Training and validation represent the balanced classification accuracy of the random forest model that was fit using consensus features from the cnCV on full training data (70 samples) and independent test data (30 samples), respectively.

#### 5.3 Comparing imbalance-adjusted fixed-k, random forest, and ridge regression

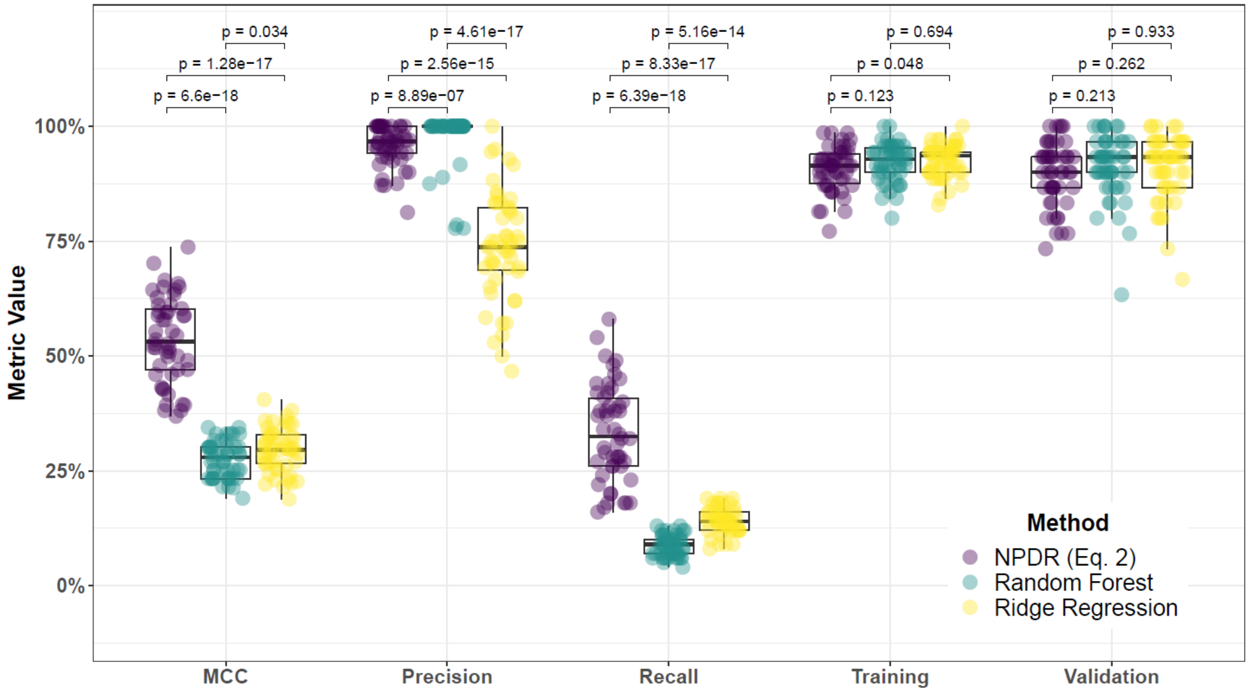


**Supplementary Fig. 50. Performance comparison of NPDR with minority-class-k (Eq. 2), Random Forest (RF), and Ridge Regression (RR) and consensus-features nested Cross-Validation (cnCV) on balanced data.** Performance of feature selection was measured for 50 simulation replicates. Each simulated data set had $m = 100$ instances and $p = 1000$ features with 100 functional. Each simulated data set had balanced class groups with 50 'case' and 50 'control'. Functional features included 25 with main effect ($\text{bias}_{\text{main}} = 0.8$) only and the remaining 75 were involved in network interactions ($\text{bias}_{\text{int}} = 0.4$) and had no main effect. Matthew's Correlation Coefficient (MCC), precision, and recall were calculated based on detection of functional features (i.e. true positives). Training and validation represent the balanced classification accuracy of the random forest model that was fit using consensus features from the cnCV on full training data (70 samples) and independent test data (30 samples), respectively.

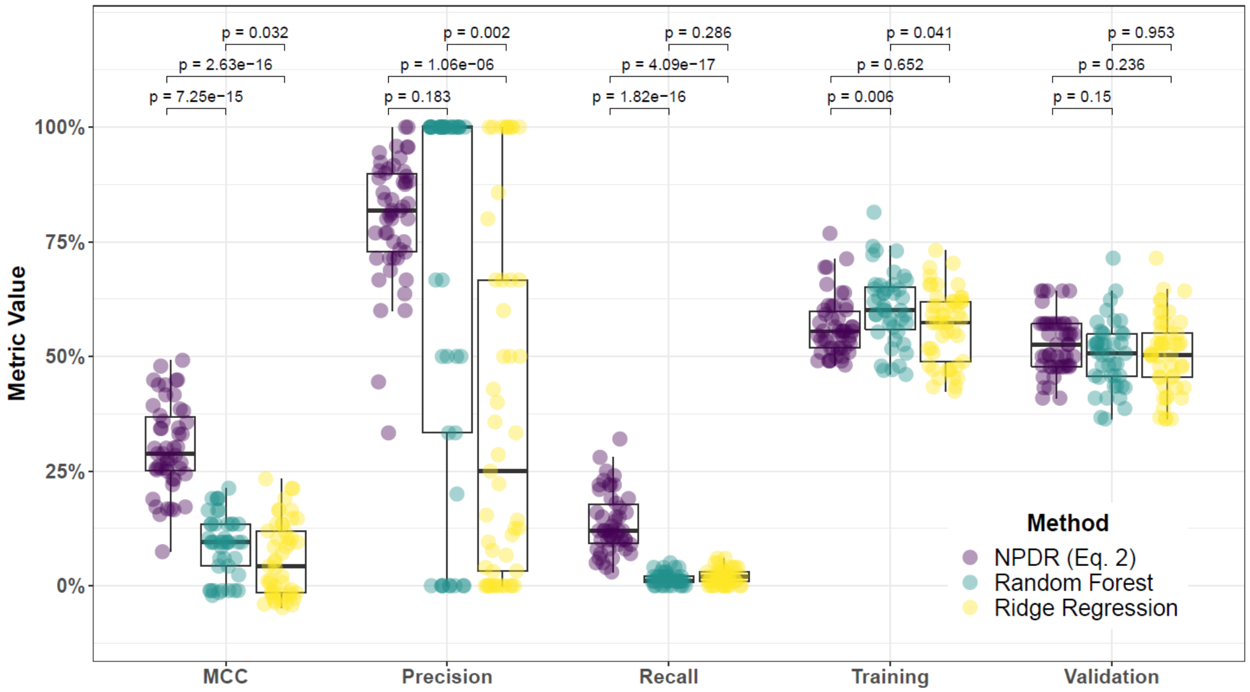


**Supplementary Fig. 51. Performance comparison of NPDR with minority-class-k (Eq. 2), Random Forest (RF), and Ridge Regression (RR) and consensus-features nested Cross-Validation (cnCV) on imbalanced data.** Performance of feature selection was measured for 50 simulation replicates. Each simulated data set had $m = 100$ instances and $p = 1000$ features with 100 functional. Each simulated data set had imbalanced class groups with 25 'case' and 75 'control'. Functional features included 25 with main effect ($\text{bias}_{\text{main}} = 0.8$) only and the remaining 75 were involved in network interactions ($\text{bias}_{\text{int}} = 0.4$) and had no main effect. Matthew's Correlation Coefficient (MCC), precision, and recall were calculated based on detection of functional features (i.e. true positives). Training and validation represent the balanced classification accuracy of the random forest model that was fit using consensus features from the cnCV on full training data (70 samples) and independent test data (30 samples), respectively.

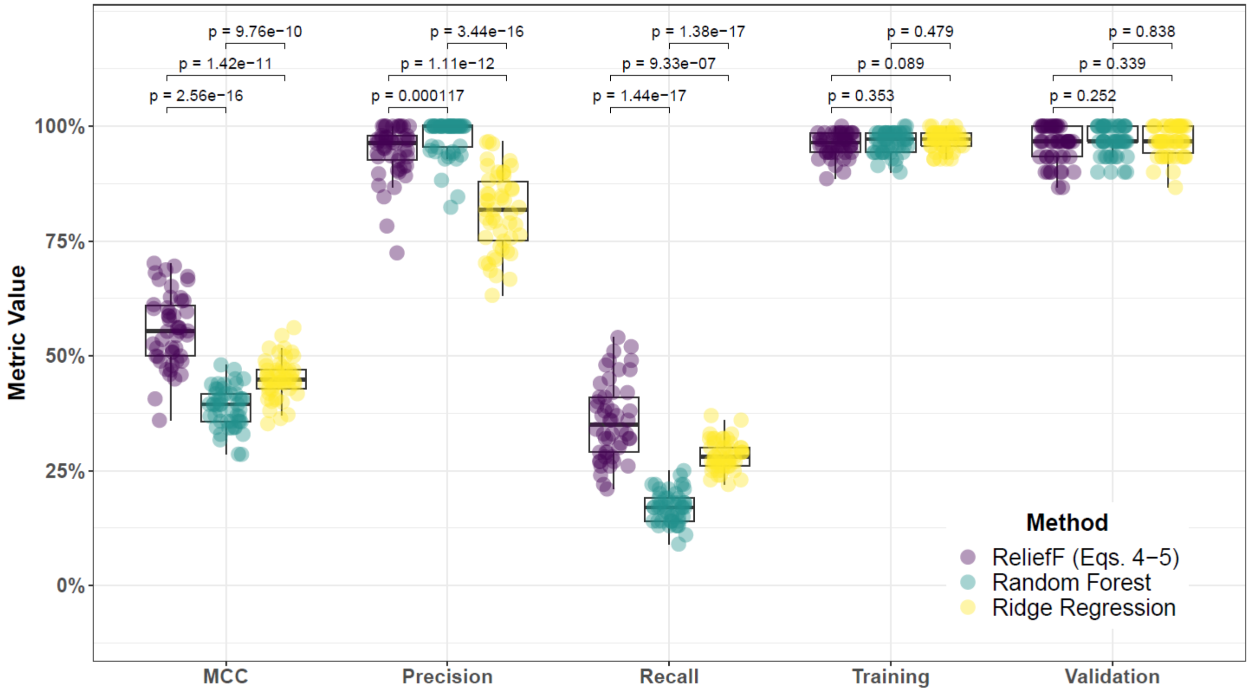


**Supplementary Fig. 52. Performance comparison for hit-miss-k (Eqs. 4 – 5) ReliefF, Random Forest (RF), and Ridge Regression (RR) and consensus-features nested Cross-Validation (cnCV) on balanced data.** Performance of feature selection was measured for 50 simulation replicates. Each simulated data set had $m = 100$ instances and $p = 1000$ features with 100 functional. Each simulated data set had balanced class groups with 50 'case' and 50 'control'. Functional features included 25 with main effect ($\mathrm{bias}_{\mathrm{main}} = 0.8$) only and the remaining 75 were involved in network interactions ($\mathrm{bias}_{\mathrm{int}} = 0.4$) and had no main effect. Matthew's Correlation Coefficient (MCC), precision, and recall were calculated based on detection of functional features (i.e. true positives). Training and validation represent the balanced classification accuracy of the random forest model that was fit using consensus features from the cnCV on full training data (70 samples) and independent test data (30 samples), respectively.

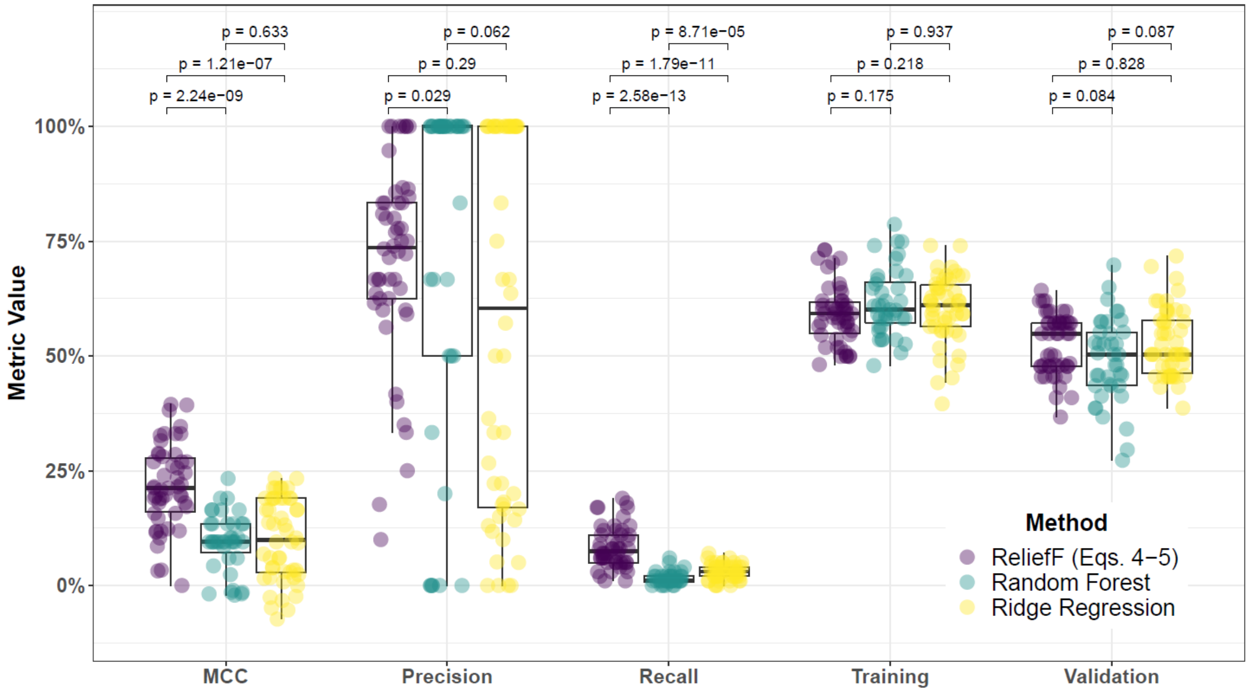


**Supplementary Fig. 53. Performance comparison for hit-miss-k (Eqs. 4 – 5) ReliefF, Random Forest (RF), and Ridge Regression (RR) and consensus-features nested Cross-Validation (cnCV) on imbalanced data.** Performance of feature selection was measured for 50 simulation replicates. Each simulated data set had $m = 100$ instances and $p = 1000$ features with 100 functional. Each simulated data set had imbalanced class groups with 25 'case' and 75 'control'. Functional features included 25 with main effect ($\text{bias}_{\text{main}} = 0.8$) only and the remaining 75 were involved in network interactions ($\text{bias}_{\text{int}} = 0.4$) and had no main effect. Matthew's Correlation Coefficient (MCC), precision, and recall were calculated based on detection of functional features (i.e. true positives). Training and validation represent the balanced classification accuracy of the random forest model that was fit using consensus features from the cnCV on full training data (70 samples) and independent test data (30 samples), respectively.

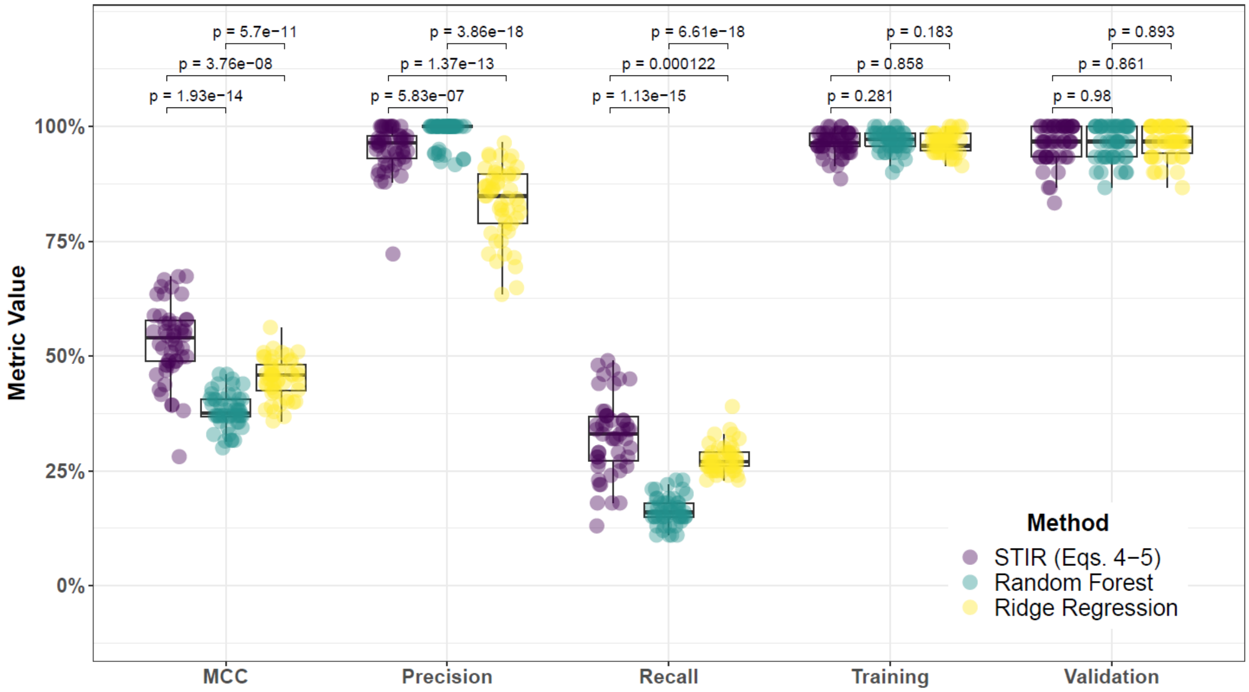


**Supplementary Fig. 54. Performance comparison for hit-miss-k (Eqs. 4 – 5) STIR, Random Forest (RF), and Ridge Regression (RR) and consensus-features nested Cross-Validation (cnCV) on balanced data.** Performance of feature selection was measured for 50 simulation replicates. Each simulated data set had $m = 100$ instances and $p = 1000$ features with 100 functional. Each simulated data set had balanced class groups with 50 'case' and 50 'control'. Functional features included 25 with main effect ($\text{bias}_{\text{main}} = 0.8$) only and the remaining 75 were involved in network interactions ($\text{bias}_{\text{int}} = 0.4$) and had no main effect. Matthew's Correlation Coefficient (MCC), precision, and recall were calculated based on detection of functional features (i.e. true positives). Training and validation represent the balanced classification accuracy of the random forest model that was fit using consensus features from the cnCV on full training data (70 samples) and independent test data (30 samples), respectively.

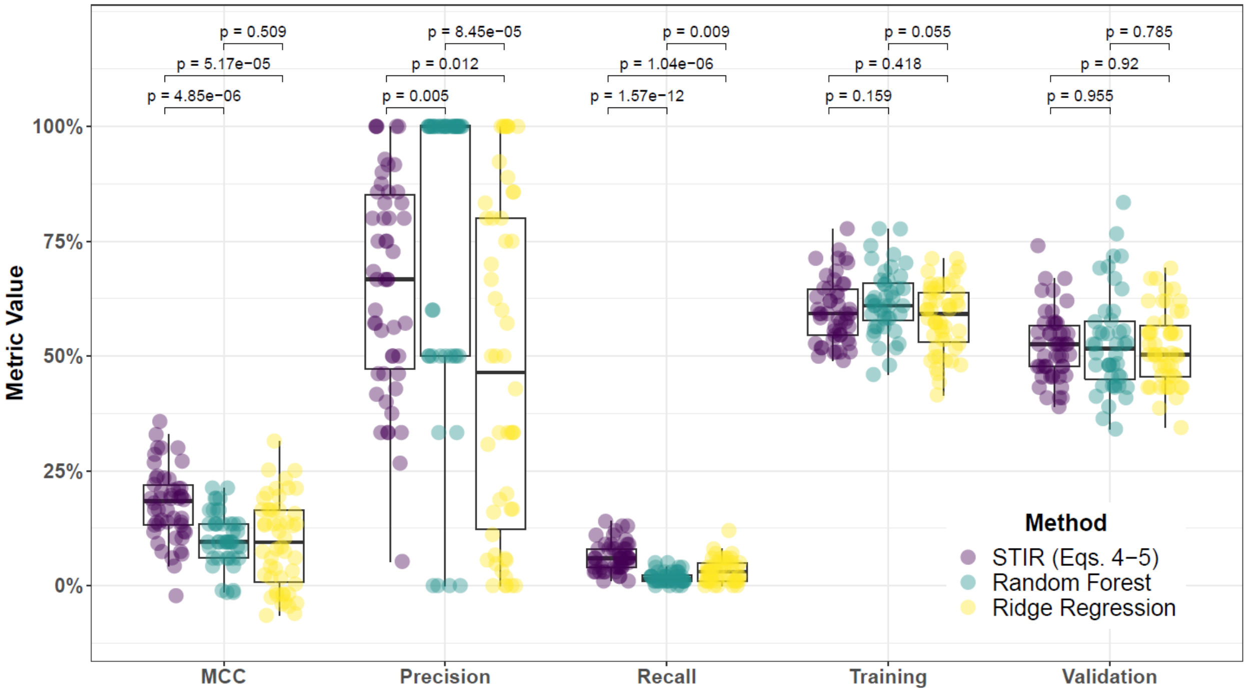


**Supplementary Fig. 55. Performance comparison for hit-miss-k (Eqs. 4 – 5) STIR, Random Forest (RF), and Ridge Regression (RR) and consensus-features nested Cross-Validation (cnCV) on imbalanced data.** Performance of feature selection was measured for 50 simulation replicates. Each simulated data set had $m = 100$ instances and $p = 1000$ features with 100 functional. Each simulated data set had imbalanced class groups with 25 'case' and 75 'control'. Functional features included 25 with main effect ($\text{bias}_{\text{main}} = 0.8$) only and the remaining 75 were involved in network interactions ($\text{bias}_{\text{int}} = 0.4$) and had no main effect. Matthew's Correlation Coefficient (MCC), precision, and recall were calculated based on detection of functional features (i.e. true positives). Training and validation represent the balanced classification accuracy of the random forest model that was fit using consensus features from the cnCV on full training data (70 samples) and independent test data (30 samples), respectively.

### 6 Feature selection performance comparisons within consensus-features nested cross-validation (cnCV): 25% interaction effect/75% main effect

#### 6.1 Comparing imbalance-adjusted fixed-k, VWOK, and kPCA

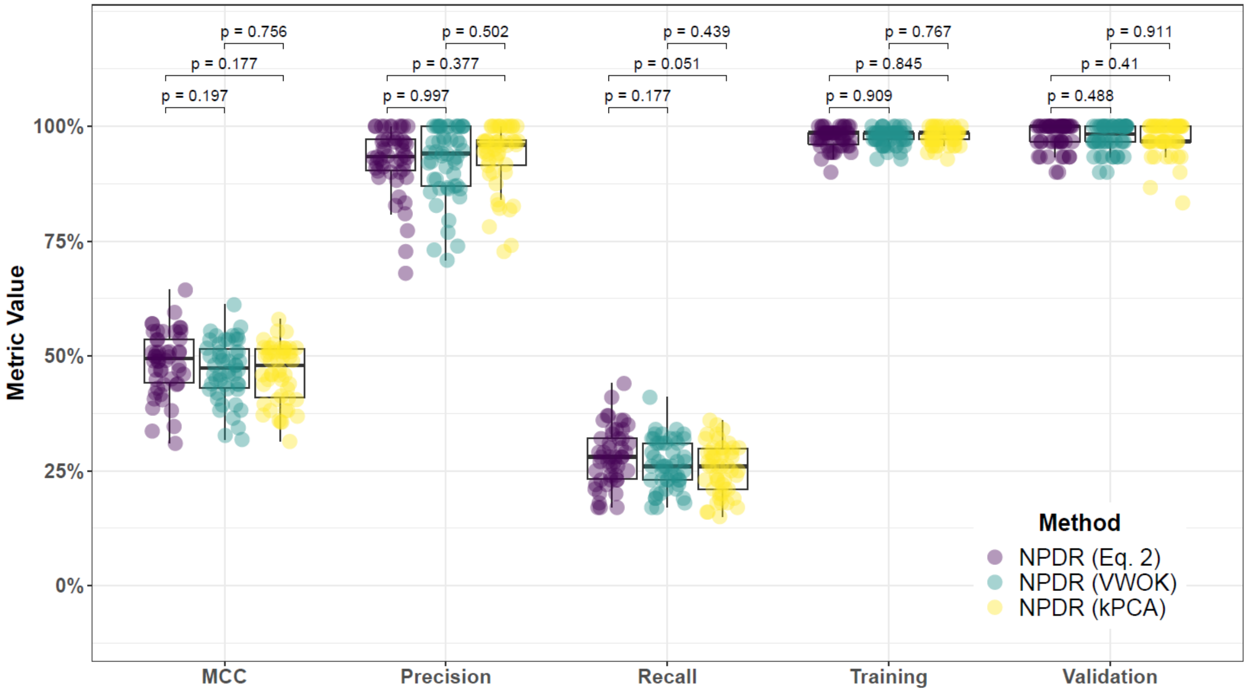


**Supplementary Fig. 56. Performance comparison for minority-class-k (Eq. 2), VWOK (Eq. 3), and kPCA with NPDR feature scoring and consensus-features nested Cross-Validation (cnCV) on balanced data.** Performance of feature selection was measured for 50 simulation replicates. Each simulated data set had $m = 100$ instances and $p = 1000$ features with 100 functional. Each simulated data set had balanced class groups with 50 'case' and 50 'control'. Functional features included 75 with main effect ($\text{bias}_{\text{main}} = 0.8$) only and the remaining 25 were involved in network interactions ($\text{bias}_{\text{int}} = 0.4$) and had no main effect. Matthew's Correlation Coefficient (MCC), precision, and recall were calculated based on detection of functional features (i.e. true positives). Training and validation represent the balanced classification accuracy of the random forest model that was fit using consensus features from the cnCV on full training data (70 samples) and independent test data (30 samples), respectively.

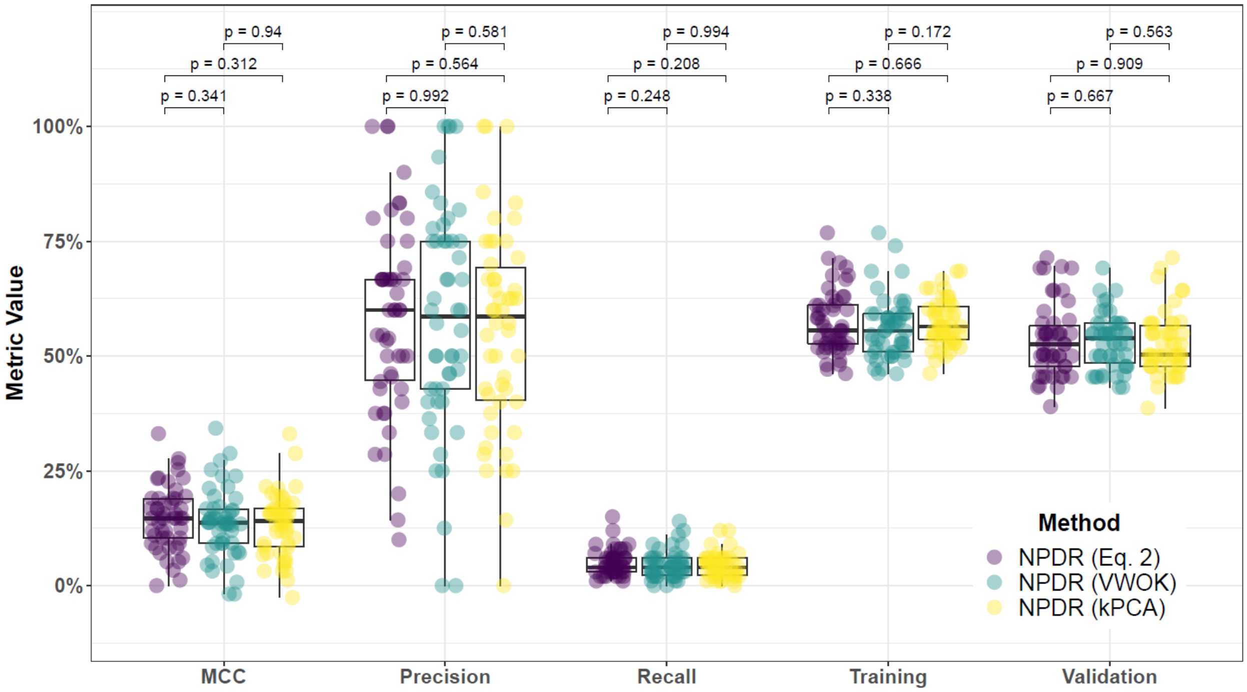


**Supplementary Fig. 57. Performance comparison for minority-class-k (Eq. 2), VWOK (Eq. 3), and kPCA with NPDR feature scoring and consensus-features nested Cross-Validation (cnCV) on imbalanced data.** Performance of feature selection was measured for 50 simulation replicates. Each simulated data set had $m = 100$ instances and $p = 1000$ features with 100 functional. Each simulated data set had imbalanced class groups with 25 'case' and 75 'control'. Functional features included 75 with main effect ($\text{bias}_{\text{main}} = 0.8$) only and the remaining 25 were involved in network interactions ($\text{bias}_{\text{int}} = 0.4$) and had no main effect. Matthew's Correlation Coefficient (MCC), precision, and recall were calculated based on detection of functional features (i.e. true positives). Training and validation represent the balanced classification accuracy of the random forest model that was fit using consensus features from the cnCV on full training data (70 samples) and independent test data (30 samples), respectively.

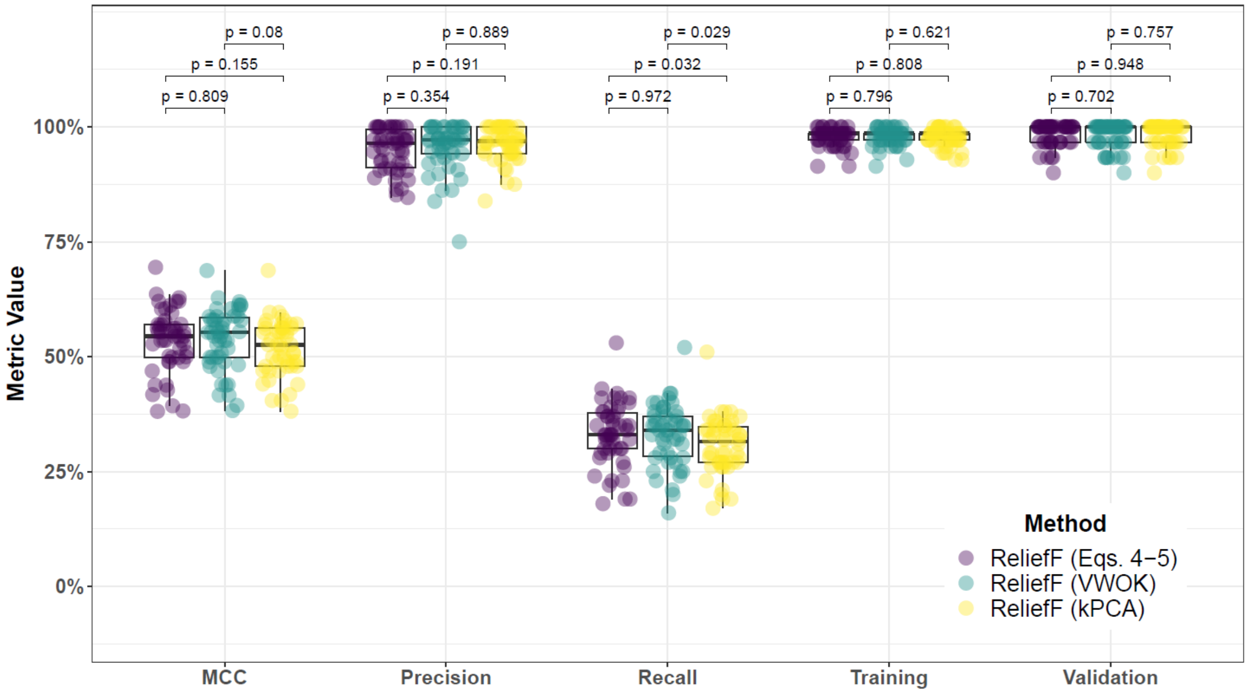


**Supplementary Fig. 58. Performance comparison for hit-miss-k (Eqs. 4 – 5), VWOK (Eqs. 6 – 7), and kPCA with ReliefF feature scoring and consensus-features nested Cross-Validation (cnCV) on balanced data.** Performance of feature selection was measured for 50 simulation replicates. Each simulated data set had $m = 100$ instances and $p = 1000$ features with 100 functional. Each simulated data set had balanced class groups with 50 'case' and 50 'control'. Functional features included 75 with main effect ($\text{bias}_{\text{main}} = 0.8$) only and the remaining 25 were involved in network interactions ($\text{bias}_{\text{int}} = 0.4$) and had no main effect. Matthew's Correlation Coefficient (MCC), precision, and recall were calculated based on detection of functional features (i.e. true positives). Training and validation represent the balanced classification accuracy of the random forest model that was fit using consensus features from the cnCV on full training data (70 samples) and independent test data (30 samples), respectively.

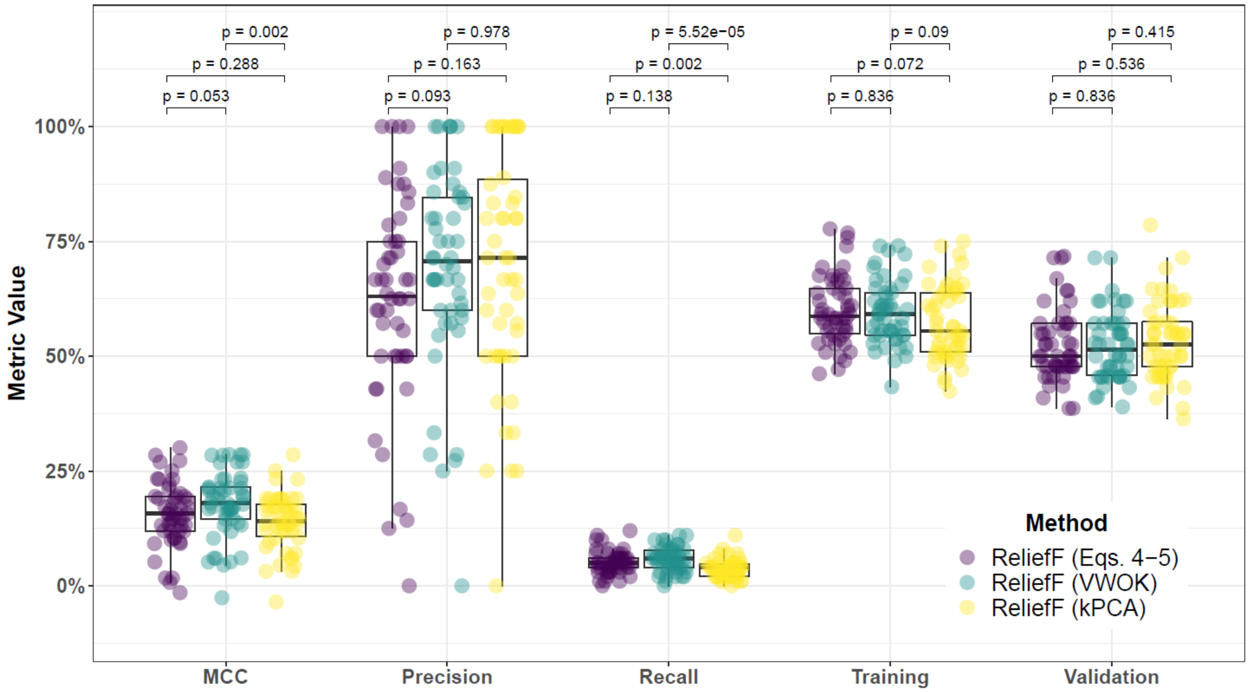


**Supplementary Fig. 59. Performance comparison for hit-miss-k (Eqs. 4 – 5), VWOK (Eqs. 6 – 7), and kPCA with ReliefF feature scoring and consensus-features nested Cross-Validation (cnCV) on imbalanced data.** Performance of feature selection was measured for 50 simulation replicates. Each simulated data set had $m = 100$ instances and $p = 1000$ features with 100 functional. Each simulated data set had imbalanced class groups with 25 'case' and 75 'control'. Functional features included 75 with main effect ($\mathrm{bias}_{\mathrm{main}} = 0.8$) only and the remaining 25 were involved in network interactions ($\mathrm{bias}_{\mathrm{int}} = 0.4$) and had no main effect. Matthew's Correlation Coefficient (MCC), precision, and recall were calculated based on detection of functional features (i.e. true positives). Training and validation represent the balanced classification accuracy of the random forest model that was fit using consensus features from the cnCV on full training data (70 samples) and independent test data (30 samples), respectively.

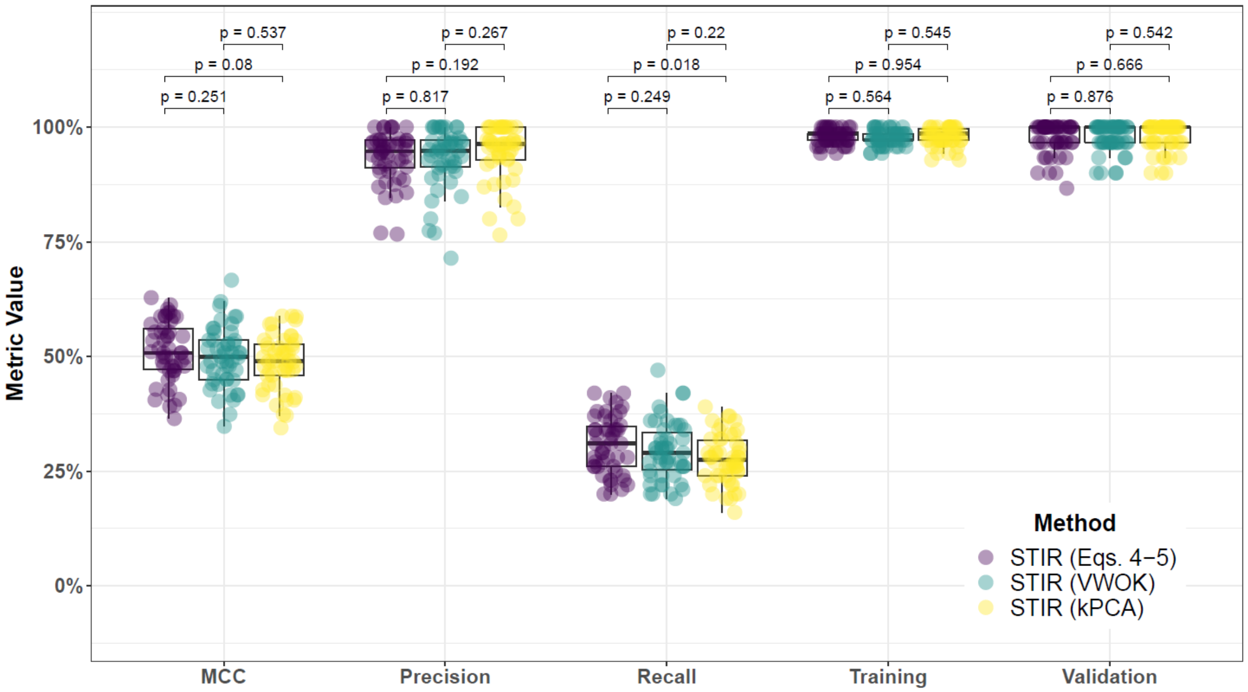


**Supplementary Fig. 60. Performance comparison for hit-miss-k (Eqs. 4 – 5), VWOK (Eqs. 6 – 7), and kPCA with STIR feature scoring and consensus-features nested Cross-Validation (cnCV) on balanced data.** Performance of feature selection was measured for 50 simulation replicates. Each simulated data set had $m = 100$ instances and $p = 1000$ features with 100 functional. Each simulated data set had balanced class groups with 50 'case' and 50 'control'. Functional features included 75 with main effect ($\text{bias}_{\text{main}} = 0.8$) only and the remaining 25 were involved in network interactions ($\text{bias}_{\text{int}} = 0.4$) and had no main effect. Matthew's Correlation Coefficient (MCC), precision, and recall were calculated based on detection of functional features (i.e. true positives). Training and validation represent the balanced classification accuracy of the random forest model that was fit using consensus features from the cnCV on full training data (70 samples) and independent test data (30 samples), respectively.

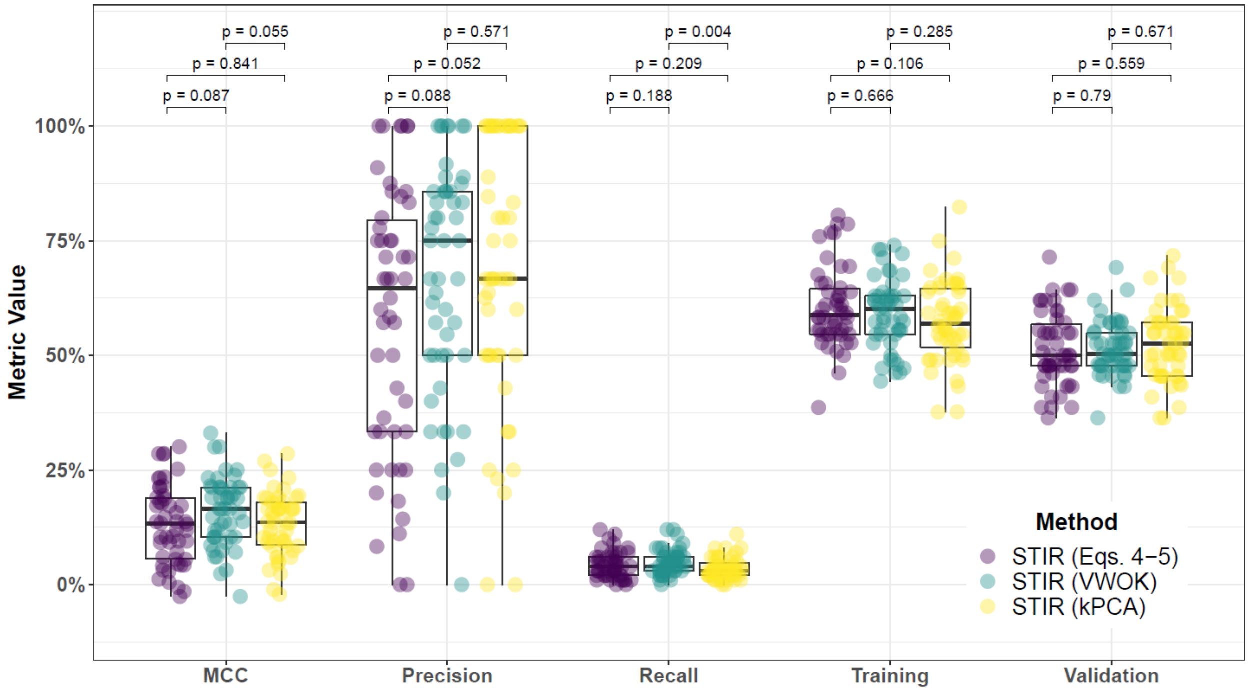


**Supplementary Fig. 61. Performance comparison for hit-miss-k (Eqs. 4 – 5), VWOK (Eqs. 6 – 7), and kPCA with STIR feature scoring and consensus-features nested Cross-Validation (cnCV) on imbalanced data.** Performance of feature selection was measured for 50 simulation replicates. Each simulated data set had $m = 100$ instances and $p = 1000$ features with 100 functional. Each simulated data set had imbalanced class groups with 25 'case' and 75 'control'. Functional features included 75 with main effect ($\text{bias}_{\text{main}} = 0.8$) only and the remaining 25 were involved in network interactions ($\text{bias}_{\text{int}} = 0.4$) and had no main effect. Matthew's Correlation Coefficient (MCC), precision, and recall were calculated based on detection of functional features (i.e. true positives). Training and validation represent the balanced classification accuracy of the random forest model that was fit using consensus features from the cnCV on full training data (70 samples) and independent test data (30 samples), respectively.

##### 6.2 Comparing imbalance-adjusted fixed-k, regular fixed-k, and MultiSURF

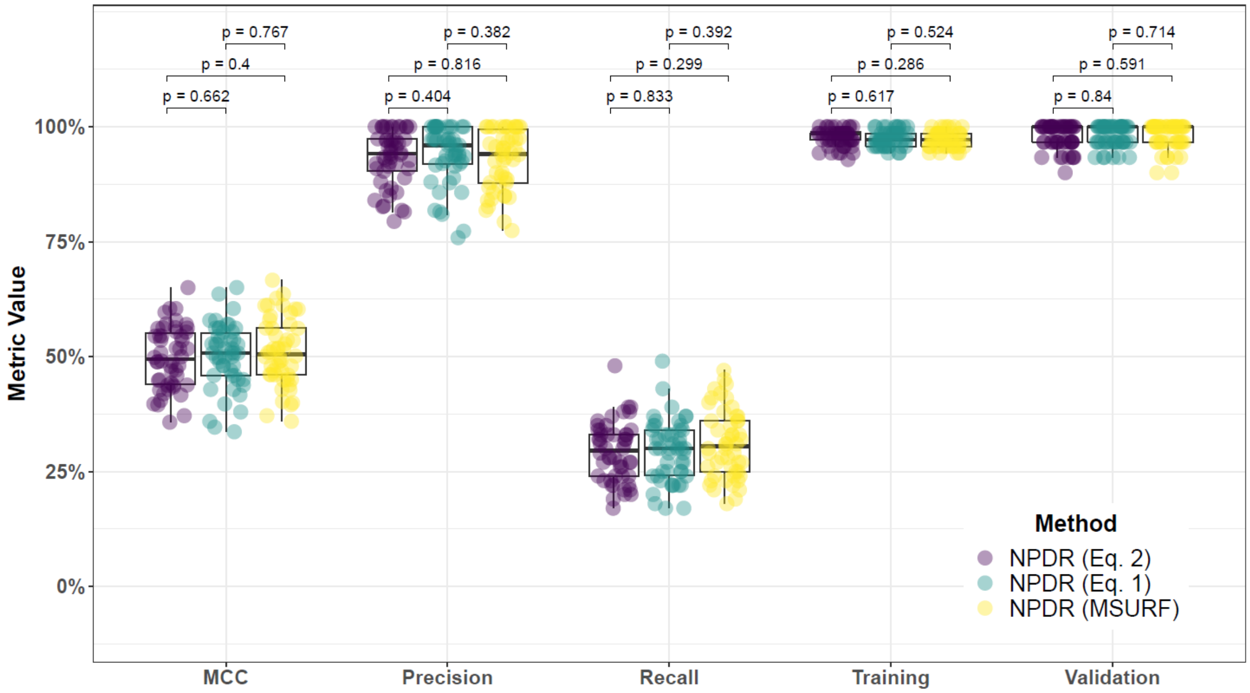


**Supplementary Fig. 62. Performance comparison for minority-class-k (Eq. 2), non-adjusted fixed-k (Eq. 1), and MultiSURF with NPDR feature scoring and consensus-features nested Cross-Validation (cnCV) on balanced data.** Performance of feature selection was measured for 50 simulation replicates. Each simulated data set had $m = 100$ instances and $p = 1000$ features with 100 functional. Each simulated data set had balanced class groups with 50 'case' and 50 'control'. Functional features included 75 with main effect ($\text{bias}_{\text{main}} = 0.8$) only and the remaining 25 were involved in network interactions ($\text{bias}_{\text{int}} = 0.4$) and had no main effect. Matthew's Correlation Coefficient (MCC), precision, and recall were calculated based on detection of functional features (i.e. true positives). Training and validation represent the balanced classification accuracy of the random forest model that was fit using consensus features from the cnCV on full training data (70 samples) and independent test data (30 samples), respectively.

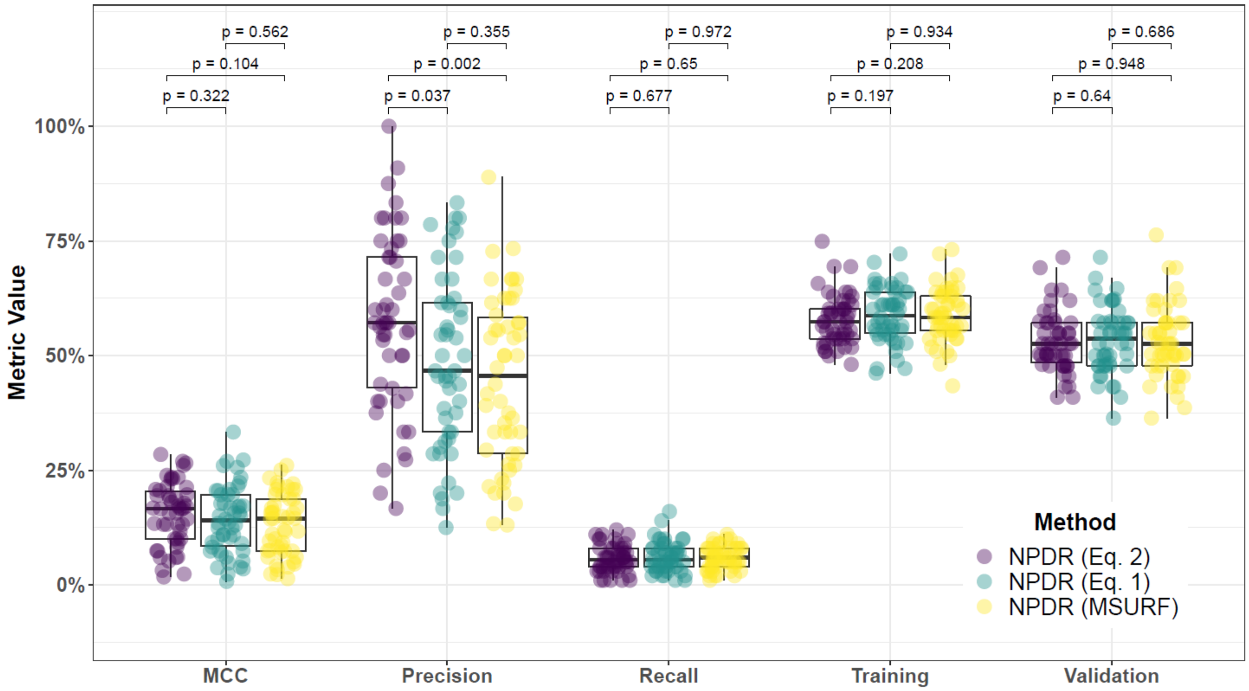


**Supplementary Fig. 63. Performance comparison for minority-class-k (Eq. 2), non-adjusted fixed-k (Eq. 1), and MultiSURF with NPDR feature scoring and consensus-features nested Cross-Validation (cnCV) on imbalanced data.** Performance of feature selection was measured for 50 simulation replicates. Each simulated data set had $m = 100$ instances and $p = 1000$ features with 100 functional. Each simulated data set had imbalanced class groups with 25 'case' and 75 'control'. Functional features included 75 with main effect ($\text{bias}_{\text{main}} = 0.8$) only and the remaining 25 were involved in network interactions ($\text{bias}_{\text{int}} = 0.4$) and had no main effect. Matthew's Correlation Coefficient (MCC), precision, and recall were calculated based on detection of functional features (i.e. true positives). Training and validation represent the balanced classification accuracy of the random forest model that was fit using consensus features from the cnCV on full training data (70 samples) and independent test data (30 samples), respectively.

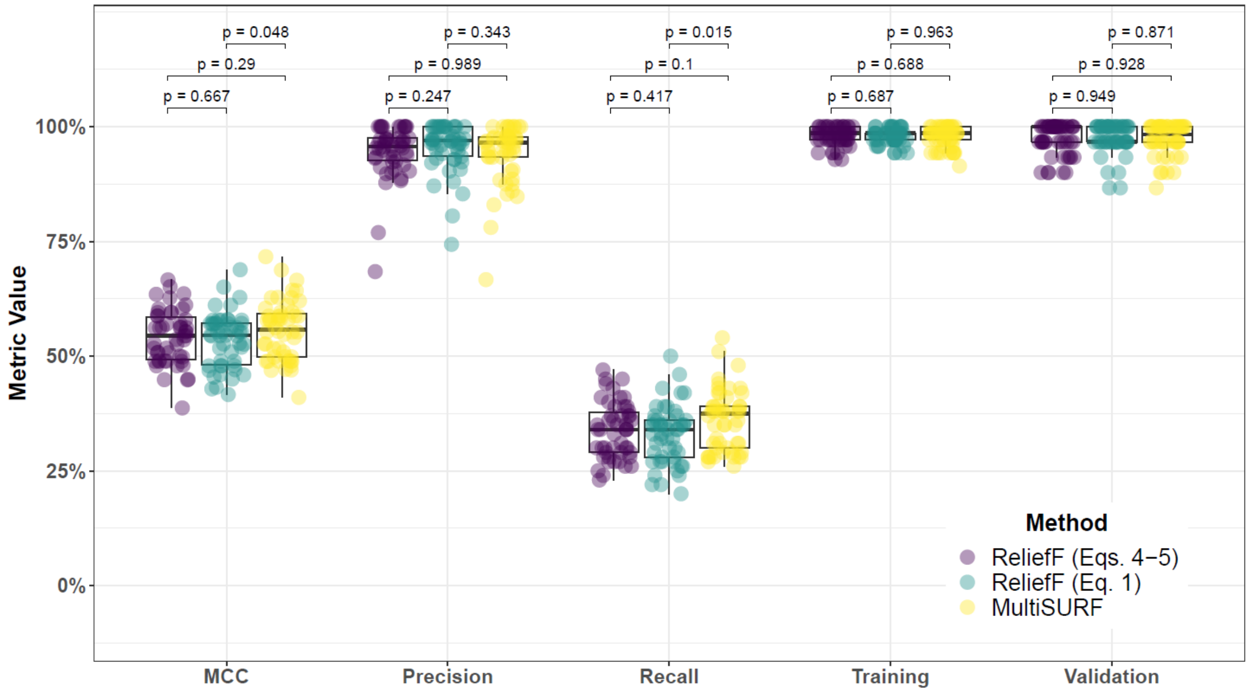


**Supplementary Fig. 64. Performance comparison for hit-miss-k (Eqs. 4 – 5) ReliefF, non-adjusted fixed-k (Eq. 1) ReliefF, and MultiSURF and consensus-features nested Cross-Validation (cnCV) on balanced data.** Performance of feature selection was measured for 50 simulation replicates. Each simulated data set had $m = 100$ instances and $p = 1000$ features with 100 functional. Each simulated data set had balanced class groups with 50 'case' and 50 'control'. Functional features included 75 with main effect ($\text{bias}_{\text{main}} = 0.8$) only and the remaining 25 were involved in network interactions ($\text{bias}_{\text{int}} = 0.4$) and had no main effect. Matthew's Correlation Coefficient (MCC), precision, and recall were calculated based on detection of functional features (i.e. true positives). Training and validation represent the balanced classification accuracy of the random forest model that was fit using consensus features from the cnCV on full training data (70 samples) and independent test data (30 samples), respectively.

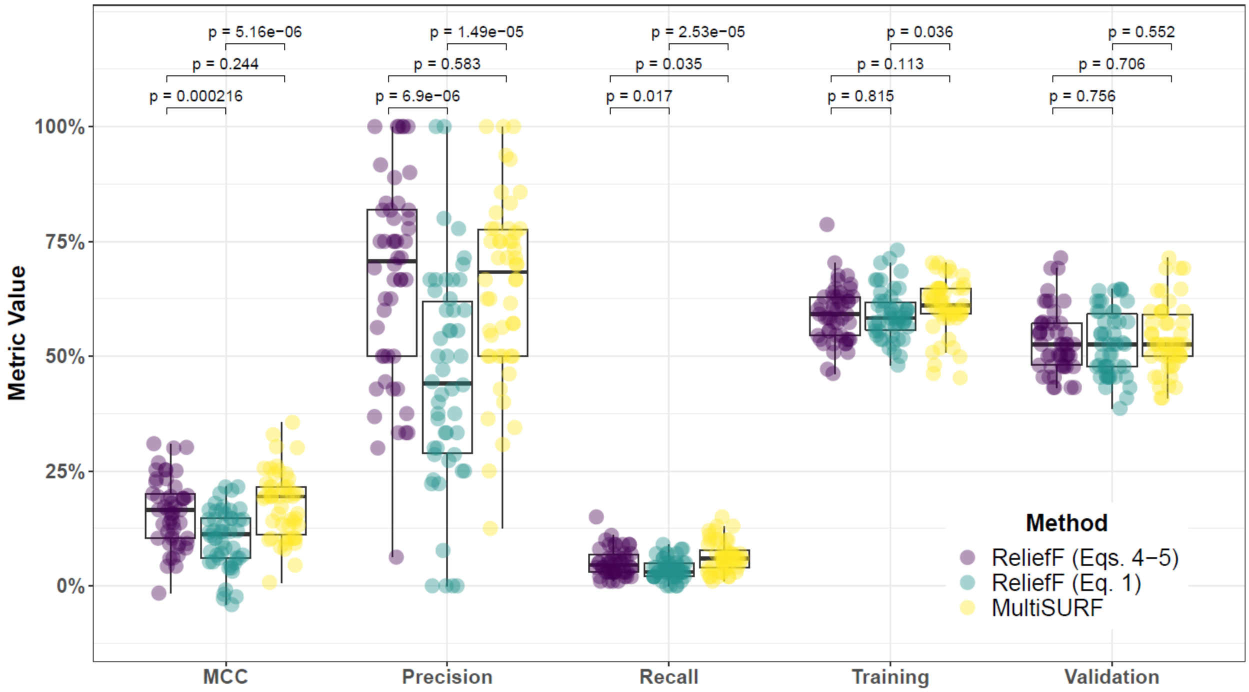


**Supplementary Fig. 65. Performance comparison for hit-miss-k (Eqs. 4 – 5) ReliefF, non-adjusted fixed-k (Eq. 1) ReliefF, and MultiSURF and consensus-features nested Cross-Validation (cnCV) on imbalanced data.** Performance of feature selection was measured for 50 simulation replicates. Each simulated data set had $m = 100$ instances and $p = 1000$ features with 100 functional. Each simulated data set had imbalanced class groups with 25 'case' and 75 'control'. Functional features included 75 with main effect ($\text{bias}_{\text{main}} = 0.8$) only and the remaining 25 were involved in network interactions ($\text{bias}_{\text{int}} = 0.4$) and had no main effect. Matthew's Correlation Coefficient (MCC), precision, and recall were calculated based on detection of functional features (i.e. true positives). Training and validation represent the balanced classification accuracy of the random forest model that was fit using consensus features from the cnCV on full training data (70 samples) and independent test data (30 samples), respectively.

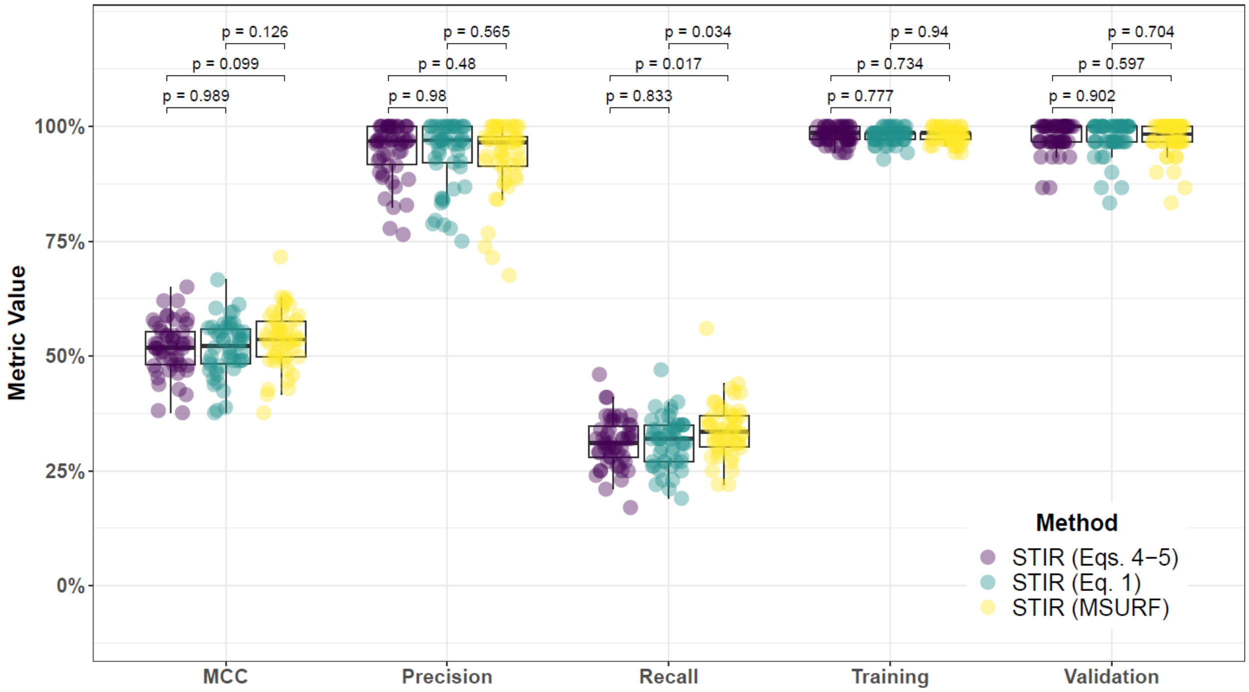


**Supplementary Fig. 66. Performance comparison for hit-miss-k (Eqs. 4 – 5) STIR, non-adjusted fixed-k (Eq. 1) STIR, and MultiSURF and consensus-features nested Cross-Validation (cnCV) on balanced data.** Performance of feature selection was measured for 50 simulation replicates. Each simulated data set had $m = 100$ instances and $p = 1000$ features with 100 functional. Each simulated data set had balanced class groups with 50 ‘case’ and 50 ‘control’. Functional features included 75 with main effect ($\text{bias}_{\text{main}} = 0.8$) only and the remaining 25 were involved in network interactions ($\text{bias}_{\text{int}} = 0.4$) and had no main effect. Matthew’s Correlation Coefficient (MCC), precision, and recall were calculated based on detection of functional features (i.e. true positives). Training and validation represent the balanced classification accuracy of the random forest model that was fit using consensus features from the cnCV on full training data (70 samples) and independent test data (30 samples), respectively.

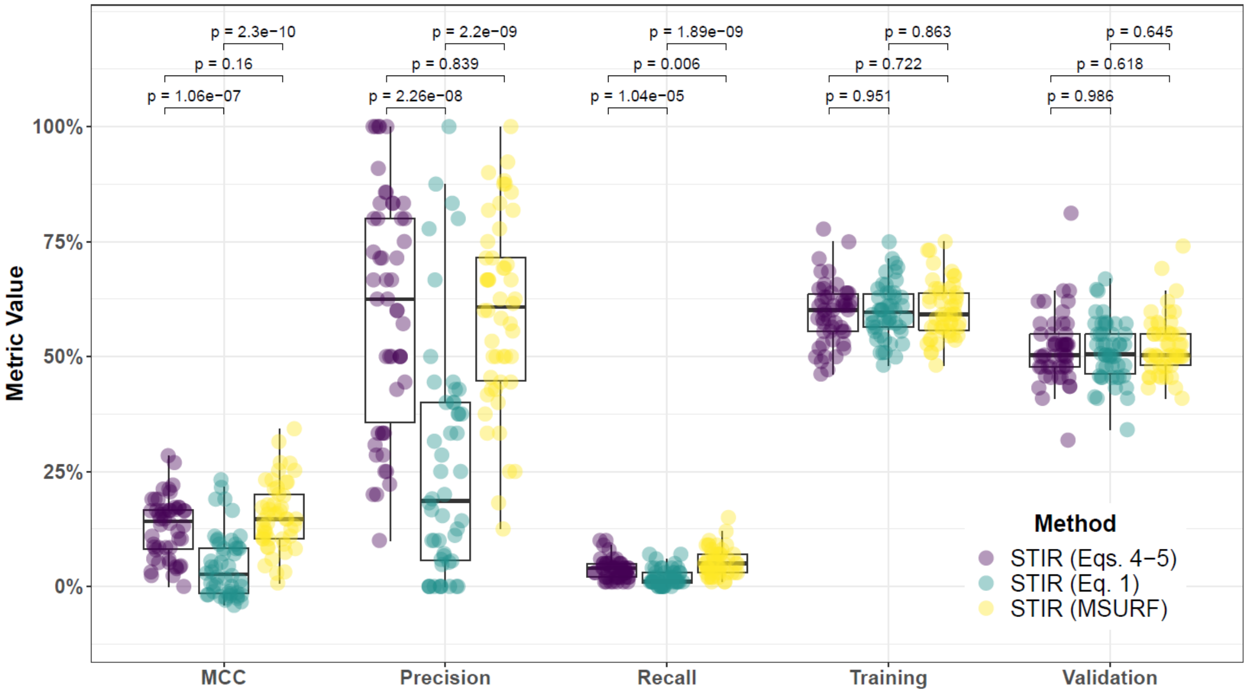


**Supplementary Fig. 67. Performance comparison for hit-miss-k (Eqs. 4 – 5) STIR, non-adjusted fixed-k (Eq. 1) STIR, and MultiSURF and consensus-features nested Cross-Validation (cnCV) on imbalanced data.** Performance of feature selection was measured for 50 simulation replicates. Each simulated data set had $m = 100$ instances and $p = 1000$ features with 100 functional. Each simulated data set had imbalanced class groups with 25 'case' and 75 'control'. Functional features included 75 with main effect ($\text{bias}_{\text{main}} = 0.8$) only and the remaining 25 were involved in network interactions ($\text{bias}_{\text{int}} = 0.4$) and had no main effect. Matthew's Correlation Coefficient (MCC), precision, and recall were calculated based on detection of functional features (i.e. true positives). Training and validation represent the balanced classification accuracy of the random forest model that was fit using consensus features from the cnCV on full training data (70 samples) and independent test data (30 samples), respectively.

##### 6.3 Comparing imbalance-adjusted fixed-k, random forest, and ridge regression

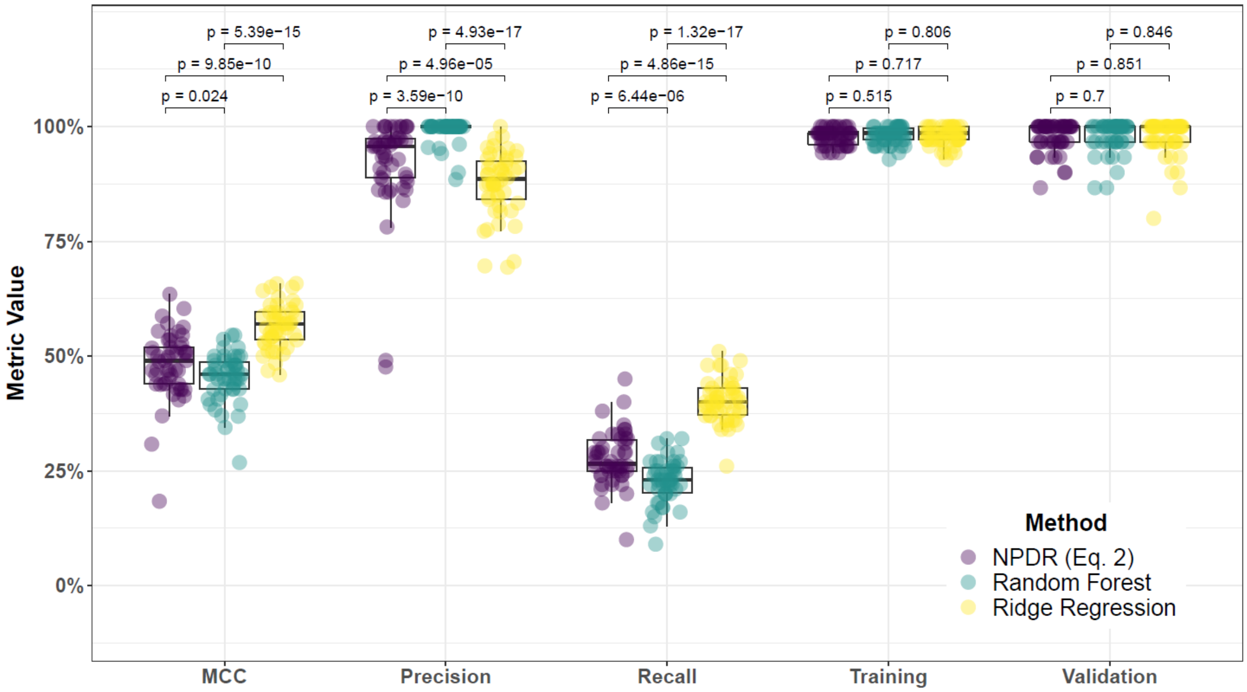


**Supplementary Fig. 68. Performance comparison of NPDR with minority-class-k (Eq. 2), Random Forest (RF), and Ridge Regression (RR) and consensus-features nested Cross-Validation (cnCV) on balanced data.** Performance of feature selection was measured for 50 simulation replicates. Each simulated data set had $m = 100$ instances and $p = 1000$ features with 100 functional. Each simulated data set had balanced class groups with 50 'case' and 50 'control'. Functional features included 75 with main effect ($\text{bias}_{\text{main}} = 0.8$) only and the remaining 25 were involved in network interactions ($\text{bias}_{\text{int}} = 0.4$) and had no main effect. Matthew's Correlation Coefficient (MCC), precision, and recall were calculated based on detection of functional features (i.e. true positives). Training and validation represent the balanced classification accuracy of the random forest model that was fit using consensus features from the cnCV on full training data (70 samples) and independent test data (30 samples), respectively.

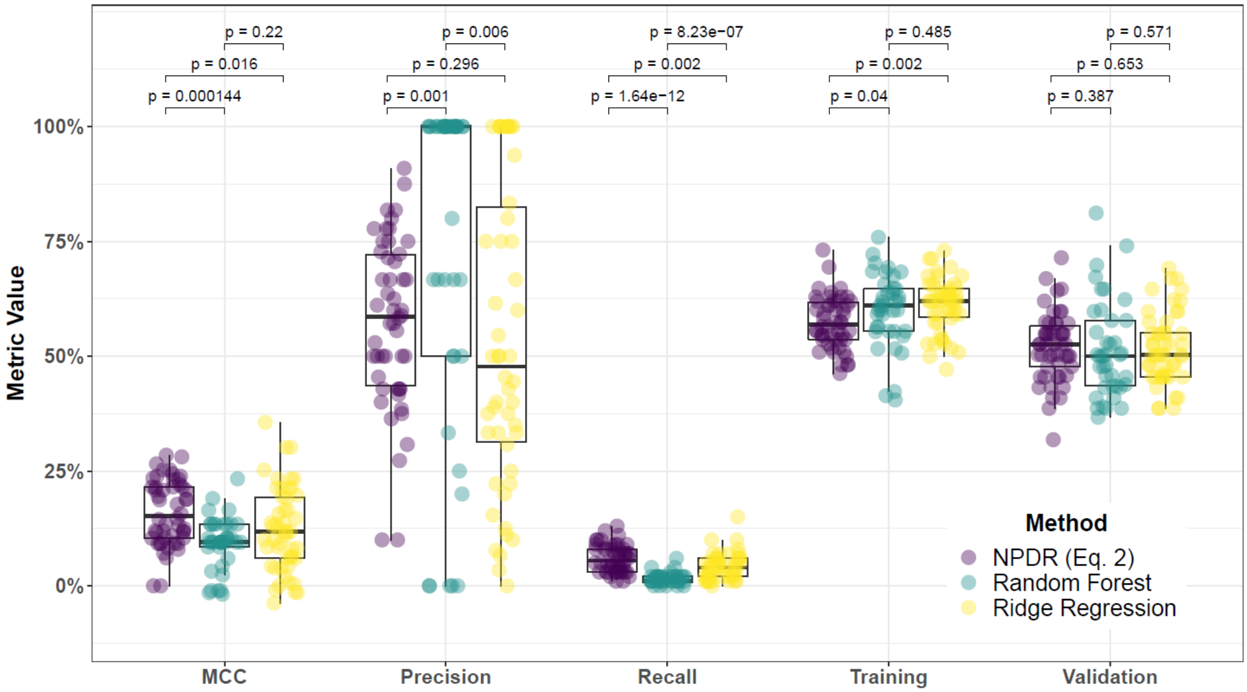


**Supplementary Fig. 69. Performance comparison of NPDR with minority-class-k (Eq. 2), Random Forest (RF), and Ridge Regression (RR) and consensus-features nested Cross-Validation (cnCV) on imbalanced data.** Performance of feature selection was measured for 50 simulation replicates. Each simulated data set had $m = 100$ instances and $p = 1000$ features with 100 functional. Each simulated data set had imbalanced class groups with 25 'case' and 75 'control'. Functional features included 75 with main effect ($\text{bias}_{\text{main}} = 0.8$) only and the remaining 25 were involved in network interactions ($\text{bias}_{\text{int}} = 0.4$) and had no main effect. Matthew's Correlation Coefficient (MCC), precision, and recall were calculated based on detection of functional features (i.e. true positives). Training and validation represent the balanced classification accuracy of the random forest model that was fit using consensus features from the cnCV on full training data (70 samples) and independent test data (30 samples), respectively.

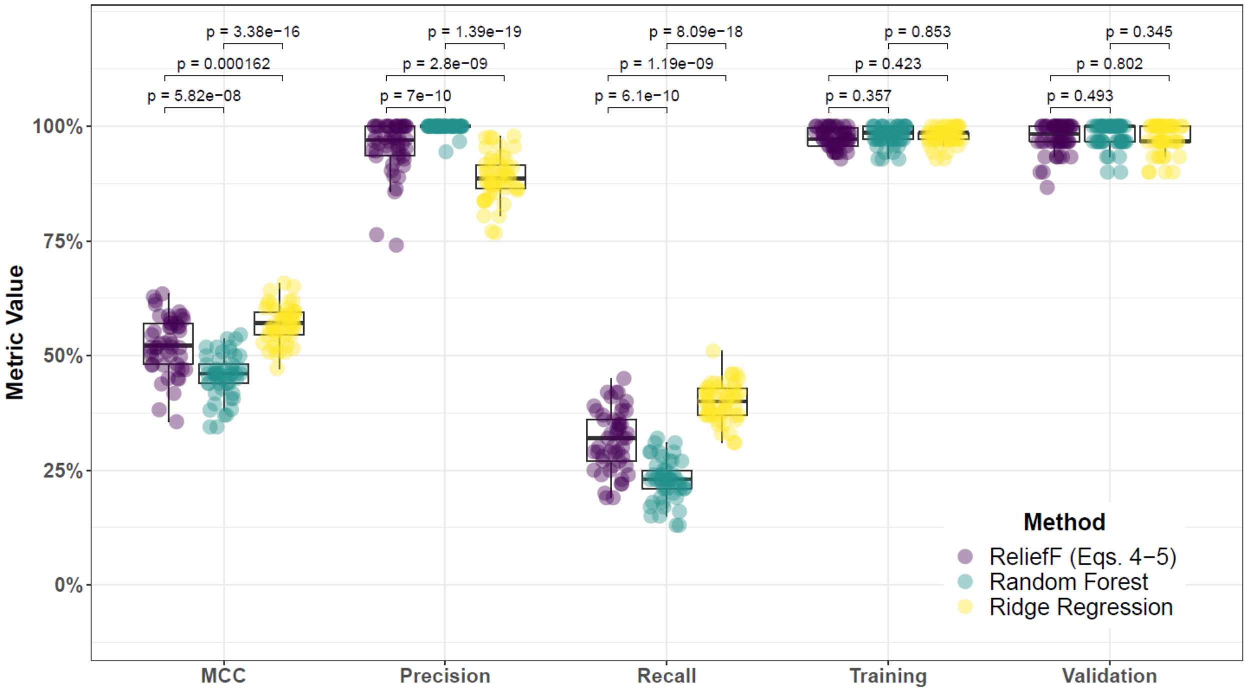


**Supplementary Fig. 70. Performance comparison for hit-miss-k (Eqs. 4 – 5) ReliefF, Random Forest (RF), and Ridge Regression (RR) and consensus-features nested Cross-Validation (cnCV) on balanced data.** Performance of feature selection was measured for 50 simulation replicates. Each simulated data set had $m = 100$ instances and $p = 1000$ features with 100 functional. Each simulated data set had balanced class groups with 50 ‘case’ and 50 ‘control’. Functional features included 75 with main effect ($\text{bias}_{\text{main}} = 0.8$) only and the remaining 25 were involved in network interactions ($\text{bias}_{\text{int}} = 0.4$) and had no main effect. Matthew’s Correlation Coefficient (MCC), precision, and recall were calculated based on detection of functional features (i.e. true positives). Training and validation represent the balanced classification accuracy of the random forest model that was fit using consensus features from the cnCV on full training data (70 samples) and independent test data (30 samples), respectively.

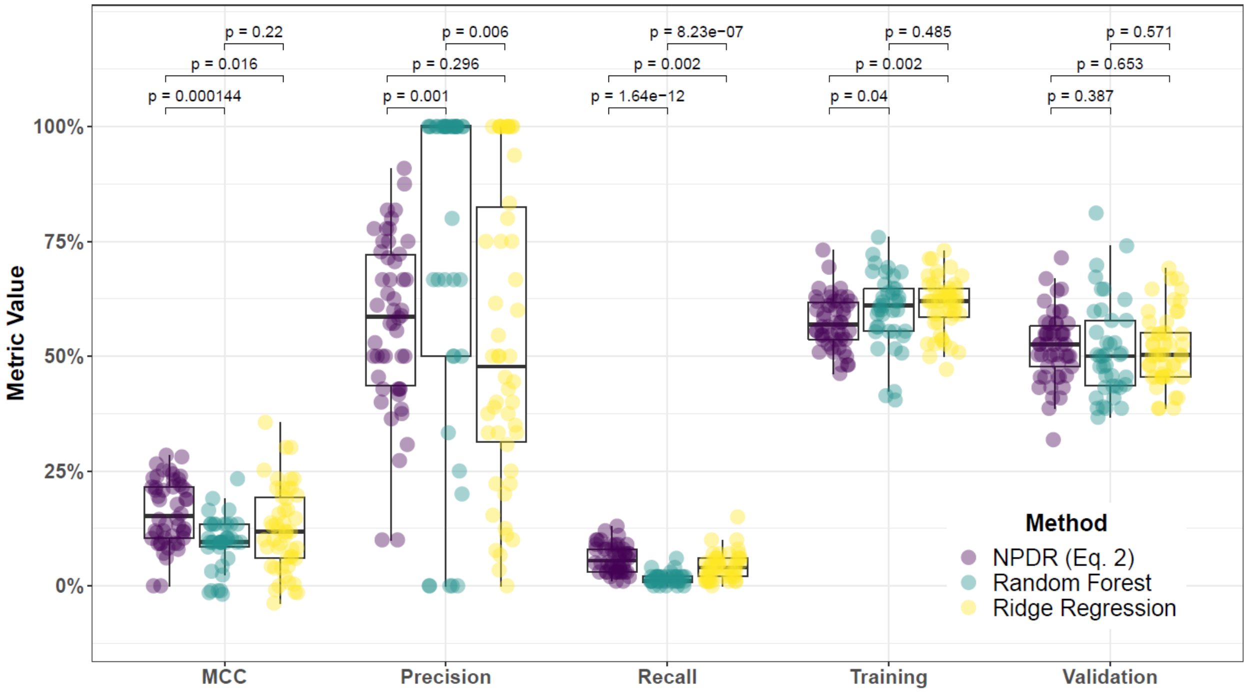


**Supplementary Fig. 71. Performance comparison for hit-miss-k (Eqs. 4 – 5) ReliefF, Random Forest (RF), and Ridge Regression (RR) and consensus-features nested Cross-Validation (cnCV) on imbalanced data.** Performance of feature selection was measured for 50 simulation replicates. Each simulated data set had $m = 100$ instances and $p = 1000$ features with 100 functional. Each simulated data set had imbalanced class groups with 25 'case' and 75 'control'. Functional features included 75 with main effect ($\text{bias}_{\text{main}} = 0.8$) only and the remaining 25 were involved in network interactions ($\text{bias}_{\text{int}} = 0.4$) and had no main effect. Matthew's Correlation Coefficient (MCC), precision, and recall were calculated based on detection of functional features (i.e. true positives). Training and validation represent the balanced classification accuracy of the random forest model that was fit using consensus features from the cnCV on full training data (70 samples) and independent test data (30 samples), respectively.

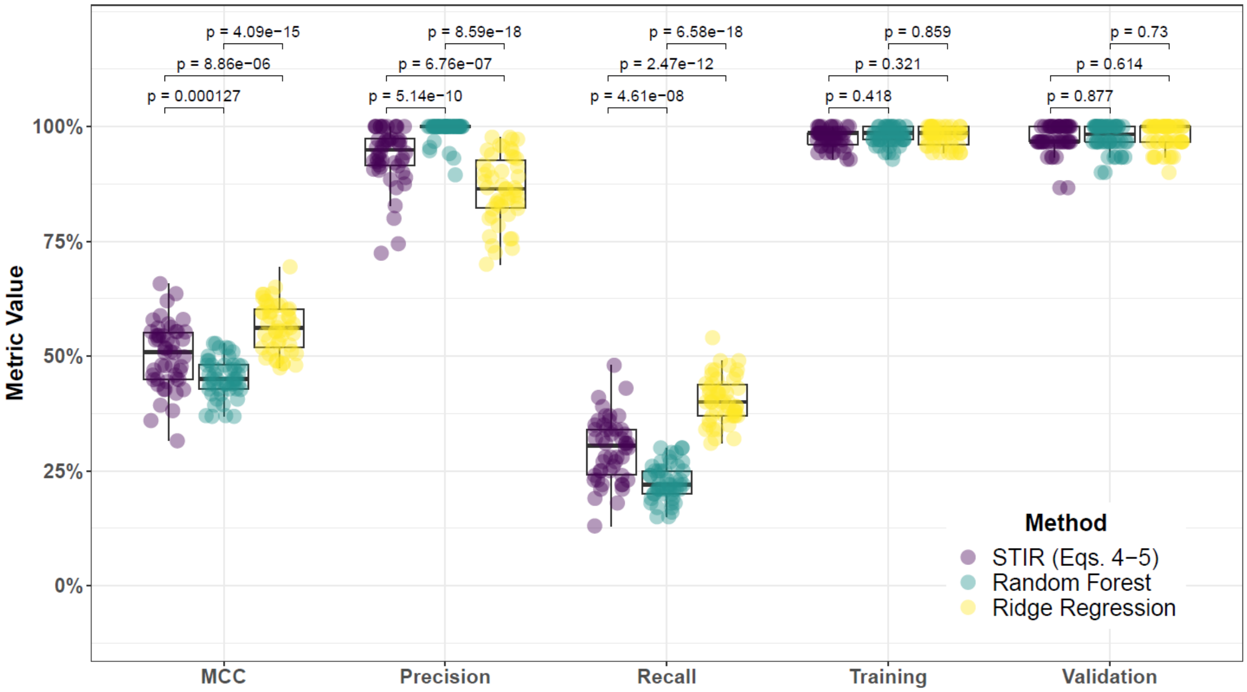


**Supplementary Fig. 72. Performance comparison for hit-miss-k (Eqs. 4 – 5) STIR, Random Forest (RF), and Ridge Regression (RR) and consensus-features nested Cross-Validation (cnCV) on balanced data.** Performance of feature selection was measured for 50 simulation replicates. Each simulated data set had $m = 100$ instances and $p = 1000$ features with 100 functional. Each simulated data set had balanced class groups with 50 'case' and 50 'control'. Functional features included 75 with main effect ($\text{bias}_{\text{main}} = 0.8$) only and the remaining 25 were involved in network interactions ($\text{bias}_{\text{int}} = 0.4$) and had no main effect. Matthew's Correlation Coefficient (MCC), precision, and recall were calculated based on detection of functional features (i.e. true positives). Training and validation represent the balanced classification accuracy of the random forest model that was fit using consensus features from the cnCV on full training data (70 samples) and independent test data (30 samples), respectively.

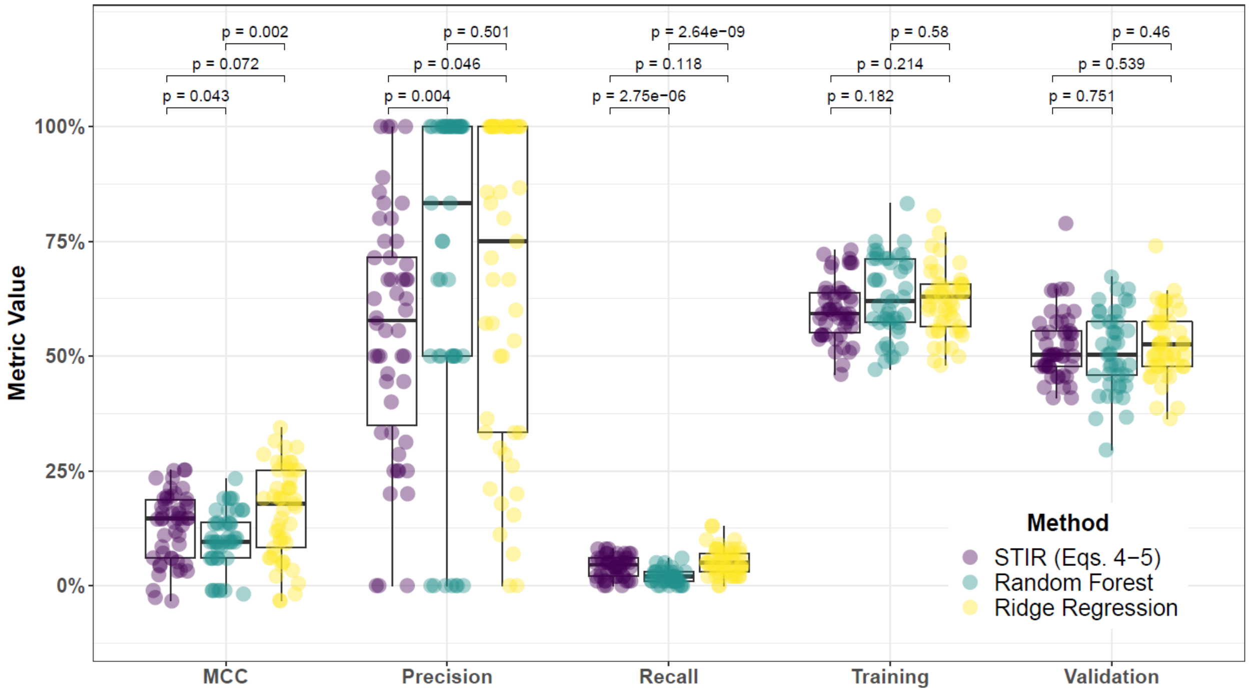


**Supplementary Fig. 73. Performance comparison for hit-miss-k (Eqs. 4 – 5) STIR, Random Forest (RF), and Ridge Regression (RR) and consensus-features nested Cross-Validation (cnCV) on imbalanced data.** Performance of feature selection was measured for 50 simulation replicates. Each simulated data set had $m = 100$ instances and $p = 1000$ features with 100 functional. Each simulated data set had imbalanced class groups with 25 'case' and 75 'control'. Functional features included 75 with main effect ($\text{bias}_{\text{main}} = 0.8$) only and the remaining 25 were involved in network interactions ($\text{bias}_{\text{int}} = 0.4$) and had no main effect. Matthew's Correlation Coefficient (MCC), precision, and recall were calculated based on detection of functional features (i.e. true positives). Training and validation represent the balanced classification accuracy of the random forest model that was fit using consensus features from the cnCV on full training data (70 samples) and independent test data (30 samples), respectively.

**Supplementary Table 1. Biological pathways for MDD-associated genes detected using cnCV with importance scores calculated using NPDR with minority-class-k (Eq. 2), NPDR with non-adjusted fixed-k (Eq. 1), and Ridge Regression.**

| Method | Pathway | Source | q-value† (FDR) | Input Overlap‡ | Size ¶ |
|---|---|---|---|---|---|
| cnCV-NPDR (Eq. 2) | Hyperornithinemia with gyrate atrophy (HOGA) | SMPDB | 0.0011 | CPS1; CKB | 20 |
| cnCV-NPDR (Eq. 2) | Creatine deficiency, guanidinoacetate methyltransferase deficiency | SMPDB | 0.0011 | CPS1; CKB | 20 |
| cnCV-NPDR (Eq. 2) | L-arginine:glycine amidinotransferase deficiency | SMPDB | 0.0011 | CPS1; CKB | 20 |
| cnCV-NPDR (Eq. 2) | Hyperornithinemia-hyperammonemia-homocitrullinuria [HHH-syndrome] | SMPDB | 0.0011 | CPS1; CKB | 20 |
| cnCV-NPDR (Eq. 2) | Guanidinoacetate Methyltransferase Deficiency (GAMT Deficiency) | SMPDB | 0.0011 | CPS1; CKB | 20 |
| cnCV-NPDR (Eq. 2) | Prolinemia Type II | SMPDB | 0.0011 | CPS1; CKB | 20 |
| cnCV-NPDR (Eq. 2) | Prolidase Deficiency (PD) | SMPDB | 0.0011 | CPS1; CKB | 20 |
| cnCV-NPDR (Eq. 2) | Arginine and Proline Metabolism | SMPDB | 0.0011 | CPS1; CKB | 20 |
| cnCV-NPDR (Eq. 2) | Hyperprolinemia Type I | SMPDB | 0.0011 | CPS1; CKB | 20 |
| cnCV-NPDR (Eq. 2) | Hyperprolinemia Type II | SMPDB | 0.0011 | CPS1; CKB | 20 |
| cnCV-NPDR (Eq. 2) | Ornithine Aminotransferase Deficiency (OAT Deficiency) | SMPDB | 0.0011 | CPS1; CKB | 20 |
| cnCV-NPDR (Eq. 2) | Arginine: Glycine Amidinotransferase Deficiency (AGAT Deficiency) | SMPDB | 0.0011 | CPS1; CKB | 20 |
| cnCV-NPDR (Eq. 2) | Phase 0 - rapid depolarisation | Reactome | 0.0012 | SCN5A; SCN8A | 22 |
| cnCV-NPDR (Eq. 2) | Interaction between L1 and Ankyrins | Reactome | 0.0020 | SCN5A; SCN8A | 29 |
| cnCV-NPDR (Eq. 2) | Primary focal segmental glomerulosclerosis (FSGS) | Wikipathways | 0.0116 | MKI67; PLCE1 | 73 |
| cnCV-NPDR (Eq. 2) | Platelet activation, signaling and aggregation | Reactome | 0.0132 | DGKK; VWF; CD9 | 263 |
| cnCV-NPDR (Eq. 2) | Phosphatidylinositol signaling system - | KEGG | 0.0171 | DGKK; PLCE1 | 97 |

| | | | | | |
|---|---|---|---|---|---|
| | Homo sapiens (human) | | | | |
| cnCV-NPDR (Eq. 2) | Glycerophospholipid metabolism - Homo sapiens (human) | KEGG | 0.0171 | DGKK; GPD1 | 98 |
| cnCV-NPDR (Eq. 2) | L1CAM interactions | Reactome | 0.0175 | SCN5A; SCN8A | 102 |
| cnCV-NPDR (Eq. 2) | Cardiac conduction | Reactome | 0.0176 | SCN5A; SCN8A | 105 |
| cnCV-NPDR (Eq. 1) | Pyrimidine nucleotides nucleosides metabolism | INOH | 0.0934 | TYMS; CPS1 | 51 |
| cnCV-Ridge | Photodynamic therapy-induced HIF-1 survival signaling | Wikipathways | 0.0170 | IGFBP2; IGFBP3 | 36 |
| cnCV-Ridge | IGF signaling | INOH | 0.0170 | IGFBP2; IGFBP3 | 36 |
| cnCV-Ridge | Myometrial relaxation and contraction pathways | Wikipathways | 0.0170 | IGFBP2; IGFBP3; ADCY6 | 155 |
| cnCV-Ridge | Graft-versus-host disease - Homo sapiens (human) | KEGG | 0.0170 | KIR2DL3; KIR2DL1 | 42 |
| cnCV-Ridge | Transport of small molecules | Reactome | 0.0170 | ABCA10; ANO5; ADCY6; SLC5A11; MCOLN3 | 641 |
| cnCV-Ridge | Antigen processing and presentation - Homo sapiens (human) | KEGG | 0.0404 | KIR2DL3; KIR2DL1 | 78 |
| cnCV-Ridge | Stimuli-sensing channels | Reactome | 0.0497 | ANO5; MCOLN3 | 94 |

†P-value for over-representation calculated from hypergeometric test, based on the number of genes in predefined set and user-input set. Multiple testing correction performed using false discovery rate (q).
‡Genes overlapping between user-input set and predefined set.
¶Size, the number of genes in the predefined gene set.

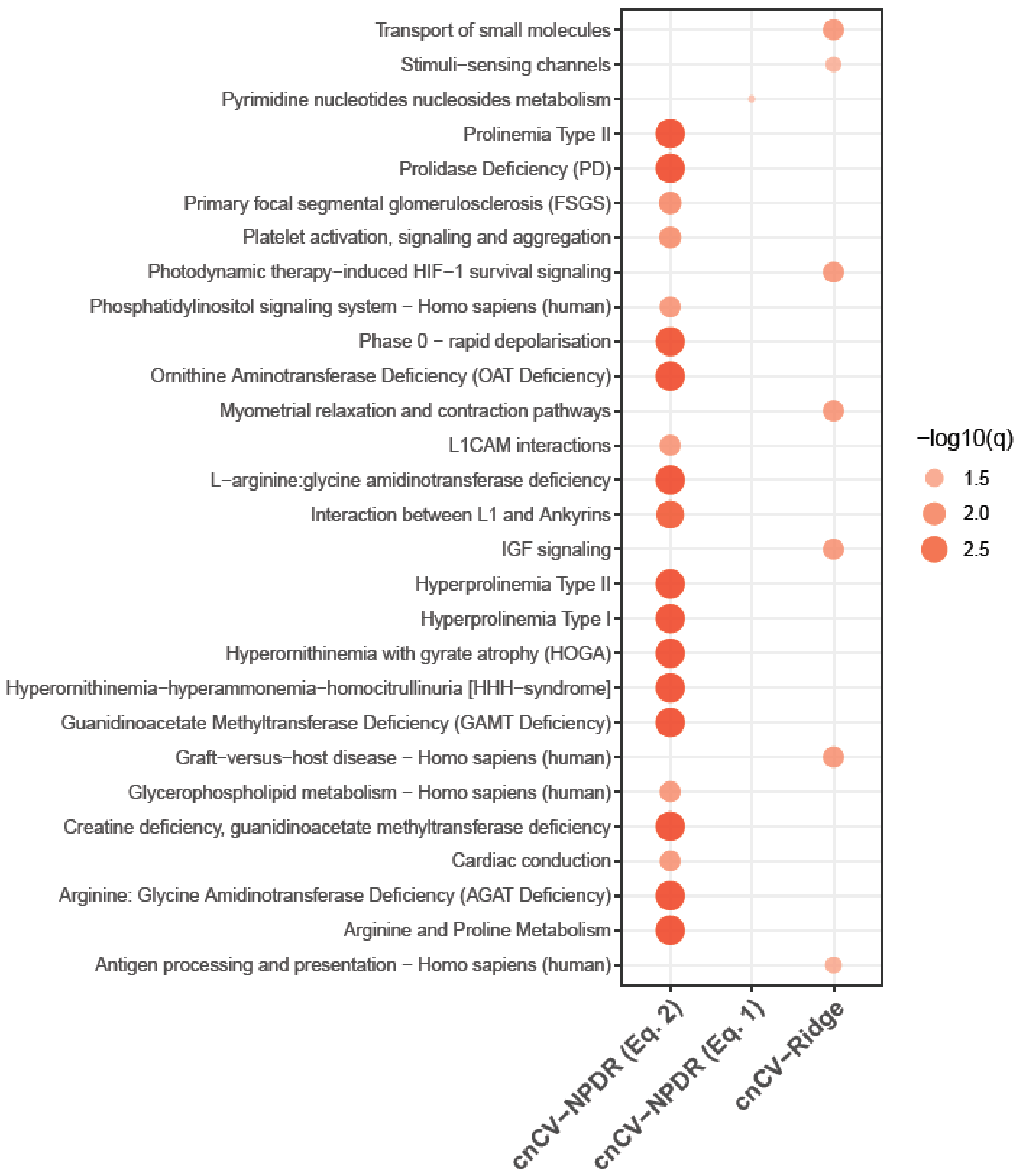


**Supplementary Fig. 74. Comparison of biological pathways for MDD-associated genes detected using cnCV with importance scores calculated by NPDR with non-adjusted fixed-k (Eq. 1), NPDR with minority-class-k (Eq. 2), and Ridge Regression.** Point color and size is proportional to the -log10 False Discovery Rate q-value, denoted by -log10(q). Biological pathways are on the y-axis and feature selection methods are on the x-axis. Pathways were calculated from a variety of databases, including Reactome, Wikipathways, KEGG, INOH, and SMPDB.

**Supplementary Table 2. Genes detected by cnCV, using NPDR non-adjusted fixed-k (Eq. 1), NPDR minority-class-k (Eq. 2), random forest, and ridge regression, on full (balanced) and sub-sampled (imbalanced) RNA-Seq data.**

| cnCV-NPDR (Eq. 1) | | cnCV-NPDR (Eq. 2) | | cnCV-Random Forest | | cnCV-Ridge Regression | |
|---|---|---|---|---|---|---|---|
| Balanced | Imbalanced | Balanced | Imbalanced | Balanced | Imbalanced | Balanced | Imbalanced |
| ADORA3 | ACTA2 | BCORP1 | ARHGEF37 | C1orf101 | RAB26 | ANO5 | ABCA10 |
| BCORP1 | B9D1 | C15orf42 | C8orf39 | GPR15 | | ARMC3 | ADCY6 |
| C9orf84 | C15orf48 | CAV2 | C9orf131 | MCOLN3 | | BEAN1 | AMAC1L2 |
| CAV2 | C8orf39 | CDCA3 | CD300LD | | | C1orf101 | ANO5 |
| CEBPE | C9orf131 | CYorf15A | CD9 | | | C5orf35 | C13orf16 |
| CLC | CD300LD | CYorf15B | CDC6 | | | C6orf174 | CORT |
| CYorf15A | CD9 | CYP2D6 | CHRNG | | | CDCA2 | EXO1 |
| CYorf15B | CDKN3 | DDX3Y | CKB | | | CELF3 | FAM198A |
| DDX3Y | CHRNG | DLGAP5 | CPS1 | | | CUX2 | GLB1L |
| EIF1AY | CPS1 | EIF1AY | DGKK | | | FAM86DP | GLT25D2 |
| FAM69B | CYP27B1 | FAM69B | DYX1C1 | | | GINS1 | GRIK1 |
| FTCD | DGKK | FTCD | FAM26E | | | GLT25D2 | GTF2H2B |
| GPR15 | DYX1C1 | GABRR2 | FLJ27352 | | | GPR15 | HTRA1 |
| HPN | FAM171A2 | GPR15 | GEMIN8P4 | | | IGFBP2 | IGFBP2 |
| HRH3 | FLJ27352 | HRH3 | GLT25D2 | | | ISL2 | IGFBP3 |
| KDM5D | GLT25D2 | ISL2 | GPD1 | | | KIAA1543 | KIR2DL1 |
| LOC347376 | GPD1 | KDM5D | GRM2 | | | LOC100131176 | KIR2DL3 |
| NCRNA00185 | GRM2 | LOC100306975 | KLHL30 | | | LOC220594 | LOC144571 |
| PLAC4 | IQGAP3 | LOC100499467 | LOC220594 | | | LOC338739 | LOC220594 |
| PRKY | KIAA1324 | MAP7D2 | LOC441208 | | | LOC400958 | LOC284648 |
| PRR19 | KLHL30 | NCRNA00185 | MEGF11 | | | LRRC37A4 | LOC81691 |
| RPS4Y1 | LOC220594 | PLAC4 | MKI67 | | | MCOLN3 | MAP1B |
| RPS4Y2 | LOC441208 | POTEKP | MTVR2 | | | MSR1 | MCOLN3 |
| RXFP2 | LOC729375 | PRKY | NPTX2 | | | NRP1 | MIR3677 |
| SLC16A14 | MKI67 | RPS4Y1 | NRXN2 | | | PMS2P4 | NMNAT2 |
| SLC4A10 | MRPL42P5 | SNORD30 | PLCE1 | | | PRSS27 | NRG2 |
| SNORD30 | MTVR2 | SOX30 | PRCD | | | RGPD1 | OIT3 |
| SNORD50A | NPTX2 | TCTEX1D4 | RAB26 | | | SDK1 | OR52K1 |
| SOX30 | NR5A1 | TK1 | SCARNA7 | | | SNORD30 | PI3 |
| STAC | NRXN2 | TPX2 | SCN5A | | | SNORD49B | PRRG2 |
| TCTEX1D4 | PRCD | TSIX | SCN8A | | | TMEM121 | PZP |
| TMSB4Y | PTPRH | TTLL10 | SNORD25 | | | WDR86 | SLC5A11 |
| TTLL10 | RAB26 | TTTY14 | SNORD76 | | | XCL2 | SNORD50B |
| TTTY10 | SCARNA7 | TTTY15 | TMEM98 | | | ZNF442 | TCTEX1D4 |
| TTTY14 | SCN8A | USP9Y | UTY | | | | XCL2 |
| TTTY15 | SKA1 | UTY | VWA5B2 | | | | |
| USP9Y | SLC16A2 | ZFY | VWF | | | | |
| UTY | SLC25A29 | | WDR63 | | | | |
| XIST | SNORD25 | | WFIKKN1 | | | | |
| ZFY | TMSB4Y | | XIST | | | | |
| | TSIX | | ZBTB8A | | | | |
| | TTTY15 | | | | | | |
| | TYMS | | | | | | |
| | UTY | | | | | | |
| | VWA5B2 | | | | | | |
| | VWF | | | | | | |
| | WDR63 | | | | | | |
| | XIST | | | | | | |
| | ZBTB8A | | | | | | |

Highlighted genes are those detected by each respective method on full (balanced) and sub-sampled (imbalanced) RNA-Seq data.
